# Hierarchical, Memory-based Movement Models for Translocated Elk (*Cervus canadensis*)

**DOI:** 10.1101/2021.04.30.442197

**Authors:** Andrea Falcón-Cortés, Denis Boyer, Evelyn Merrill, Jacqueline L. Frair, Juan Manuel Morales

## Abstract

The use of spatial memory is well documented in many animal species and has been shown to be critical for the emergence of spatial learning. Adaptive behaviors based on learning can emerge thanks to an interdependence between the acquisition of information over time and movement decisions. The study of how spatio-ecological knowledge is constructed throughout the life of an individual has not been carried out in a quantitative and comprehensive way, hindered by the lack of knowledge of the information an animal already has of its environment at the time monitoring begins. Identifying how animals use memory to make beneficial decisions is fundamental to developing a general theory of animal movement and space use. Here we propose several mobility models based on memory and perform hierarchical Bayesian inference on 11-month trajectories of 21 elk after they were released in a completely new environment. Almost all the observed animals exhibited preferential returns to previously visited patches, such that memory and random exploration phases occurred. Memory decay was mild or negligible over the study period. The fact that individual elk rapidly become used to a relatively small number of patches was consistent with the hypothesis that they seek places with predictable resources and reduced mortality risks such as predation.

## 1. INTRODUCTION

The use of spatial memory is well documented in many animal species. For example, humans, non-human primates and other large-brained vertebrates make movement decisions based on spatial representations of their environments (Wills et al. 2010). These representations may allow animals to move directly to important sites in their environment that lie outside of their perceptual range (Normand and Boesch 2009, Presotto and Izar 2010), such as resource patches or safe spots to avoid predators, and may also allow them to estimate the travel cost to reach a particular place (Janson 2007, Janson and Byrne 2007, Lanner 1996, Noser and Byrne 2007). Another type of memory, described for the first time by Schacter (1992) and retaken by Fagan et al. (2013), encodes the attributes of landscape features under the name of *attribute memory.* While spatial memory allows animals to reduce uncertainty about the location of geographical features, attribute memory reduces uncertainty concerning location-independent features of objects (Fagan et al. 2013). The information stored as attribute memory may be the abundance or types of food, and can be linked to spatial information. For example, food patch quality can be spatially encoded: patch quality is an attribute and its location is spatial information (Fagan et al. 2013). The combination of these two types of information allows animals to choose among alternative movement paths as has been observed in bumblebees (Lihoreau et al. 2011) or large herbivores (Avgar et al. 2013, Merkle et al. 2014). Identifying how animals use memory to make decisions is fundamental to developing a general theory of animal movement and space use (Gautestad and Mysterud 2005, Morales et al. 2010, Spencer 2012).

Memory is critical in the emergence of spatial learning, which results from interactions with the environment and can be detected through changes in movement patterns Mueller and Fagan (2008). Adaptive behaviors based on learning can occur thanks to an interdependence between the acquisition of information over time and movement decisions (Falcón-Cortés et al. 2017, 2019). For instance, an animal can make decisions based on past successful experiences, resulting in a change of behaviour and improved resource exploitation. Leonard (1990) showed in laboratory studies that exposure to food rewards distributed spatially in mazes can affect movement decisions in rats. Something similar occurs in ungulates that obtain foraging benefits by remembering previous trajectories while migrating under seasonal ranges (Bracis and Mueller 2017, Jesmer et al. 2018, Merkle et al. 2019). On the other hand, non-informed movements may facilitate exploration of unknown profitable areas (Bouton 2007, Miller and Shettleworth 2007). Ecological knowledge is the result of a continuous learning process through the entire life of an individual (Brent et al. 2015). For example, adult seabirds have a better knowledge of possible exploitation zones and forage more efficiently than young individuals, whose movements are more unpredictable (Grecian et al. 2018). Large herbivores that were introduced in a new environment took many years to adopt home range movements, presumably as they built up new spatial memory (Fryxell et al. 2008). Learning is consistent, for example, with frequent visits to certain locations, or site fidelity (Bonnell et al. 2013, Falcón-Cortés et al. 2017), and with the emergence of home range behaviour or preferential travel routes (Boyer and Walsh 2010, Van Moorter et al. 2009). The capability of learning can also bring other benefits beyond improved foraging; e.g. providing advantage in territorial defense (Potts and Lewis 2014, Schlägel and Lewis 2014, Schlägel et al. 2017), more effective escape from predators (Brown 2001), and improving the route choice in migration (Bischof et al. 2012, Poor et al. 2012). Nevertheless, the connections between memory and spatial learning is not well understood. Theoretical models bring useful insights by predicting, for instance, how often memory should be used for the emergence of recurrent movements to a particular resource patch (Boyer et al. 2019, Falcón-Cortés et al. 2017).

Many theoretical studies have attempted to relate spatial learning with movement and with the advantages such learning brings. These theoretical approximations are diverse. Agent-based models can be used to study the connection between cognitive abilities and foraging success (Boyer and Walsh 2010). Several studies have highlighted the role played by memory for home range formation (Berger-Tal and Avgar 2012, Börger et al. 2008, Van Moorter et al. 2009) and paved the way for inferring individual memory capacities from movement and environmental data (Avgar et al. 2013). Models that incorporate distance, resources, and memory into analyses of movement data (Dalziel et al. 2008), as well as models that mix spatial and attribute memory (Merkle et al. 2014), have revealed that large herbivores are likely to choose previously visited patches of high quality, thus offering a promising template for understanding the role of memory in animal movement. The applications of these theoretical approaches to free-ranging animals are varied. For example, predictions of a simple memory model based on linear reinforcement through preferential revisits have been compared with the movements of capuchin monkeys, revealing movement rules found to generate very slow diffusion and heterogeneous space use (Boyer and Solis-Salas 2014a). On the other hand, Merkle et al. (2014) applied a patch-to-patch model to ranging data of American bison, finding that these animals remember valuable information about the location and quality of meadows and use this information to revisit profitable locations.

The study of how spatio-ecological knowledge is constructed throughout the life of an individual has not been developed thoroughly. Data analyses that employ memory based models are promising but are often difficult to implement due to the short observation periods available, and the fact that the animals are observed in an environment already familiar to them. If memory is long-ranged, the above limitations may affect the results. To avoid these shortcomings, we used data from relocated animals. This means that the observed animals explored an unknown landscape at the start of their movement trajectories. In this new environment the spatial locations of different environmental features and patches were initially unknown to them. We analyzed the movement data from 21 relocated elk (*Cervus canadensis*) as described in (Frair et al. 2007, Wolf et al. 2009). We expected elk to show an initial exploratory phase in which the animals were getting familiarized with their new environment and collecting information about the location and quality of different habitat patches. We then expected an exploitation phase showing less random space use, eventually leading to the formation of home ranges. Furthermore, as the relocated animals came from three different sources with different degrees of similarity with the release site (see below), it is possible that some animals would show different strategies.

In a recent study, a memory-based movement model similar to the ones that we propose below was fitted to roe deer reintroduced into a novel environment, showing that home ranges in the absence of territoriality could emerge from the benefits of using memory during foraging (Ranc et al. 2020). Here we followed a similar approach, but placed emphasis on comparisons among alternative movement models. This allowed us to reveal possible differences in behaviours. We also paid special attention to the estimates of certain key parameters characterizing informed movement, such as the rate at which an animal used memory, and whether memory decayed over time and how.

We present four simple patch-to-patch movement models, defined through the probabilities of transiting from one patch to another. The simplest model is memoryless as it assumes that the transition probabilities depend only on the distance between the two patches and on the quality (size) of the target patch. The remaining three models consider the role of memory. The manner in which we introduce memory in the dynamics is very similar to that of (Boyer and Solis-Salas 2014a) and (Falcón-Cortés et al. 2017) in which the patches previously visited by the forager are reinforced. Therefore, in these memory-based models we mix spatial memory (animals remember patch locations) and attribute memory (animals remember, through reinforcement, patch quality). The main difference between these three models is the way in which animals use their memory. In the simplest case we suppose that animals have infinite memory, *i.e.,* they can remember all the patches previously visited, and they use their memory at a constant rate. In another model we assume infinite memory but the rate at which the animal decides to use its memory increases with the number of explored patches. In the last model we relax the assumption of infinite memory by introducing a memory decay associated to each patch visit (McNamara and Houston 1985), whereas the rate of memory use increases as in the previous model.

## 2. METHODS

### 2.1. Ranging data

We used data collected and presented by Frair et al. (2007); see also Wolf et al. (2009). The study area consisted of 15 800 km^2^ along the eastern slopes of the Rocky Mountains in central Alberta, Canada. Approximately 2000 elk inhabited the area during the study period, from December 2000 to September 2002 (Frair et al. 2007). Elevation was 500-1500 *m* and the area was largely forested (68.7% of the total area). Dominant tree species included lodgepole pine *Pinus Contorta,* white spruce *Picea Glauca,* and aspen/poplar *Populus Tremuloides* and *P. Balsamea.* Interspersed throughout the forested matrix were wet and dry meadows (7.1%), cutover forest following timber harvest (4.3%), bare soil/rock outcrops (12.3%), rivers and lakes (2.1%), and areas regenerating from wildfire or site reclamation (<1%) (Frair et al. 2007, Wolf et al. 2009).

Over the study period, female elk were translocated to the study area from three source sites within Alberta: 1) Banff and Jasper National Parks, mountainous areas with the full suite of predators present in the study area but protected from hunting, 2) Cross Ranch Conservation Area (ca 20 km southwest of Calgary), a hunted area of foothills and agricultural lands largely without predators, and 3) Elk Island National Park, a flat aspen parkland without predators or hunters, see (Frair et al. 2007) for more details about these three sites. Collared animals included six females from the town site of Banff released in February 2001. Nine females were released from the Cross Area, six during December 2000 and three in December 2001, and six females were released from Elk Island between January and February 2002. The animals were captured primarily using corral traps baited with hay. These animals were transported to release areas in livestock trailers that held between 9 and 16 animals depending on the sex and age class composition. Elk were released directly from the trailers into the study area. The animals were released in a number of separate locations to increase independence between results from different individuals (Frair et al. 2007, Wolf et al. 2009).

Prior to release, translocated elk were fitted with GPS collars (LMRT4 and GPS2200, Lotek Wireless, ON, Canada) that collected locations every 2 h for up to 11 months. We used all locations of each collared animal during a season or until radio-contact was lost, the animal died, or GPS collars were retrieved via breakaway device (11 months post-release). All collars were equipped with mortality sensors that activated after 7 h of immobility. Collar tests across the range of cover and terrain conditions encountered within the study area indicated a high fix rate and positional accuracy of ≤50 *m* 80% of the time (Frair et al. 2007, Wolf et al. 2009).

We identified foraging patches for elk from a 27-class landcover grid developed for this region (see Frair et al. 2005). The grid had a 28.5 *m* cell size, and an overall classification accuracy of 82.7%. Using ArcGIS (Environmental Systems Research Institute, Redlands, California), we reclassified dry/mesic and wet meadows, shrubland, clearcuts, and reclaimed herbaceous (pipeline) classes into a single foraging habitat class and converted the grid to a polygon layer without simplifying lines, which is equivalent to an 8-cell neighborhood rules for patch definition. We eliminated polygons <0.27 ha in size (essentially <3 contiguous pixels), and retained 16,782 patches for analysis. The resulting foraging patches averaged 6.93 ± 29.4 ha in size. For each elk GPS location occurring within a patch, we recorded the unique number for that patch, which allowed us to derive information on the time spent moving between foraging patches, the residency time within patches, and the return time to previously visited patches. Thus, we transformed the original GPS trajectories into a time series of patch to patch visits which included the time spent in each patch and the time travelling between patches. We assumed that most foraging occurred in these high biomass patches.

### 2.2. Models

For each model below, we made the following assumptions:

- The animals were moving in a stationary 2*d* environment which consisted of a set of *N* available patches (resource sites), *N* is obtained from environmental data as detailed in the previous subsection. Patches were characterized by their area *a_n_*, with *n* in {1, …,*N*}, taken as a proxy for resource abundance. The euclidean distance between the centroids of the patches *n* and *m* is denoted by *d_n,m_*.
- We modeled discrete movement events: at each time step *t* → *t* + 1 an animal decides to move to another patch (patch-to-patch movement) following a set of rules that we will explain below. The model does not take into account the actual time spent in a patch or between patches.
- An animal will go from patch *n* to patch *m* with probability *P_n,m_*. This probability were computed in different ways for each model.
- All the parameters to estimate were positive numbers.

#### 2.2.1. Model I

The first model is Markovian as it assumes that the forager chooses to visit a patch (*m*) in the environment by considering the distance (*d_n,m_*) from its current patch (*n*) and the area (*a_m_*) of the patch *m*. We define a probability vector *k* = (*k*_1_,…,*k_N_*) whose *m*-th entry denotes the probability that the animal goes to patch *m* from patch *n*. Each entry is defined by:

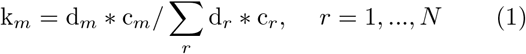

with,

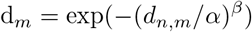

and

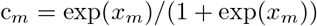

where *x_m_* = *λa_m_* + *κ*, i.e., we assume that the probability to visit patch *m* decays exponentially with the distance to patch *n* (*d_n,m_*) and increases with the area of patch *m* (*a_m_*). We aim at obtaining a hierarchical estimation for the parameters *α, β, λ* and *κ* (see Table I).

**TABLE I.** Prior distributions

| Parameter | Prior distribution | Interpretation |
| --- | --- | --- |
| $\alpha$ | normal(0,10) | Scale parameter for the exponential function that defines the probability decay with distance. |
| $\beta$ | normal(0,1) | Shape parameter for the exponential function that defines the probability decay with distance. |
| $\lambda$ | normal(0,1) | Slope parameter for the logit function that defines the probability increase as function of patch area. |
| $\kappa$ | normal(0,1) | Intercept parameter for the logit function that defines the probability increase as function of patch area. |
| $q$ | beta(1,1) | Parameter that defines the memory use frequency. |
| $\rho$ | normal(0,10) | Scale parameter for the exponential function that defines the increase of probability memory use as function of the number of unique visited sites. |
| $\epsilon$ | normal(0,1) | Shape parameter for the exponential function that defines the increase of probability memory use as function of the number of unique visited sites. |
| $\nu$ | normal(0,10) | Shape parameter for the exponential function that defines memory decay as function of time since last visit for each patch. |
| $\theta$ | normal(0,10) | Scale parameter for the exponential function that defines memory decay as function of time since last visit for each patch. |

#### 2.2.2. Model II

We next incorporate memory effects through a parameter *q* ∈ (0,1) that defines the probability with which an animal decides to use its experience to revisit a patch. In this Model II, we assume that the forager has infinite memory, i.e., is capable of remembering all previously visited sites. Linear reinforcement is implemented by setting that the probability to choose a particular site for revisit is proportional to the accumulated number of visits to that site. This model has two types of movement decisions:

- With probability *q* the forager moves from patch *n* to patch *m* considering, besides the distance and area, the number of visits that patch *m* has received in the past. The entry *m* of the probability vector **k** is now defined by:

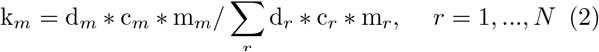

with d_*m*_ and c_*m*_ defined as in (1) and *m_m_* = *n_m_*, where *n_m_* is the number of visits at site *m* until the present time *t*. Hence, *m_m_* = 0 if the animal has never visited *m*.
- With probability 1 − *q* the forager does not use its memory and will choose a patch *m* using the probability vector **k** defined in (1). Hence the forager performs an exploratory movement.

#### 2.2.3. Model III

Given that the data trajectories belong to animals that were released in an unfamiliar environment, it is reasonable to hypothesize that movements were dominated by exploration at early times and by memory at later times. In such case, one may allow the memory parameter *q* to vary with time.

In this model, the memory parameter depends on the number of unique visited sites (UVS) of the forager up to time *t*. To this end, we define **u** = (*u*_1_,…, *u_T_*) as a vector of length *T*, with *T* the trajectory length and *u_T_* the number of distinct patches visited by the forager up to time *T* (*u*_1_ = 1). This vector is an observed data and *q* will depend on it as follows:

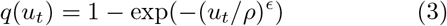

In this model the total number of parameters to estimate is six, four of them already considered in Model I, plus two parameters for the increase of memory use as i function of the UVS *(ρ* and *ϵ*. See Table I).

#### 2.2.4. Model IV

So far we have considered in Model II and III that foragers possess infinite memory. Besides, we have considered that reinforcement is linear, *i.e.*, that an animal chooses a site for revisit with probability proportional to the total number of visits to that site. To incorporate memory decay, we assume in Model IV that the weight of any visit decays exponentially in time, from the value unity. Hence, the animal will forget those visits that are far away in the past and will remember very well those that are recent. Therefore, the recently visited sites have a larger probability to be visited again.

The memory factor defined in Model II now takes the form:

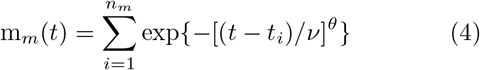

with *n_m_* the number of visits to patch *m* until time *t*, and *t_j_* the time at which the *i*-th visit to this patch occurred. It is important to note that m_*m*_ defined in Eq.(4) will be characterized by an exponential memory decay for *θ* = 1, a stretched exponential decay for *θ* < 1, and a super-exponential decay for *θ* > 1. In this model, one needs to estimate eight parameters. The six parameters already considered in Model III and two more describing memory decay (ν and *θ*. See Table I).

We fitted these four models to the data and then we performed a model comparison. We used two different tools to perform this comparison: a Posterior Predictive Check (PPC) to asses the model’s ability to “predict” the data used to parameterize it, and the Watanabe-Akaike Information Criterion (WAIC) (Watanabe and Opper 2010) as an approximation for out of sample predicting capacity of each model. These two tools help us to compare the four models above. Specifications about fitting and comparison are shown in the next sections.

### 2.3. Model fitting

For some parameters such as q, the frequency of memory use, we used non-informative priors while for other parameters we used weakly informative priors (Table I). All priors were truncated to take only non-negative values.

The models were fitted by using a two-stage approach as proposed by (Hooten and Hefley 2019). The first stage involves fitting the set of individual-level models independently using placeholder priors for all model parameters. Each individual has its own set of parameters for each model. This first-stage was achieved using Hamiltonian Monte Carlo (HMC) techniques implemented within the software Stan (Carpenter et al. 2017) and accessed via RStan (Team et al. 2018). For all models we ran three HMC chains with: 5 000 iterations each for Model I and II, and 10 000 iterations each for Model III and IV (with 2 500 (5 000) iterations for warmup, a Rhat < 1.1, and a reasonable number of effective samples (n_eff)), from which samples from the posterior distribution of all parameters were obtained.

The second stage involved a simple MCMC algorithm to fit the full hierarchical Gaussian model using the posteriors from the first stage as priors (Hooten and Hefley 2019). This second stage ran only one chain with 7 500 (15 000) iterations (the union of the three chains from the first stage) with 3 750 (7 500) iterations for warmup, a *p—*value *p_v_ >* 0.05 for the Geweke’s statistic and a reasonable n_eff for all the relevant parameters in the different model dynamics. With this second step, we obtained the posterior distributions at the individual-level for the parameters of each animal (this fit takes into account the variability between individuals) as well as the posterior distributions of the parameters at the population-level.

### 2.4. Model assessment and comparison

In order to assess and compare the descriptive and predictive capacity of the different models, we use two kinds of tools: one qualitative and the other quantitative.

As qualitative assessments, we performed PPC on the number of unique patches visited by the animals through time. That is, for each animal we determined the number of unique patches visited (or UVS) as a function of the number of between-patch movements and compared this quantity with the predictions of simulated trajectories from the different models. For each simulated trajectory, we used parameter combinations sampled from the joint posterior of each of the corresponding model. For each model and individual animal, we simulated 1 000 trajectories and we checked whether the observed change in number of UVS fell within the credible interval of the simulated ones. We thus could asses whether the observed pattern was consistent with the parametrized model.

As a quantitative assessment of model predictive capacity we used WAIC (Watanabe and Opper 2010). This quantity is computed from the log-pointwise-predictive-density of each model, which was calculated from the posterior distributions obtained from the second-stage algorithm. This quantity helped us to suggest the best model for each individual: we say that a model is the best when it obtained the lowest WAIC and when the difference between this and other model’s WAIC were grater than 2.

## 3. RESULTS

### 3.1. Model comparison

Considering the PPC for all individuals and models (Fig. 9 of the Supplemental Material (SM)), we found that 6 trajectories (out of the 21 individuals) were contained within the 95 percent credible interval (CI) of Model I, while 17 did so for Model II, 10 for Model III and 15 for Model IV.

The WAIC comparisons displayed in Table II show us that Model I was not the best model for any individual, *i.e.* the calculated WAIC for Model I was never the smallest one for any animal. Model II had the smallest WAIC for 12 individuals. Model III was the best for 9 animals, and Model IV was not the best for any individual. Therefore, in most cases, a constant rate of memory use and a linear reinforcement without memory decay provided a good description of their trajectories. These results agree qualitatively with those of the PPC.

**TABLE II.**
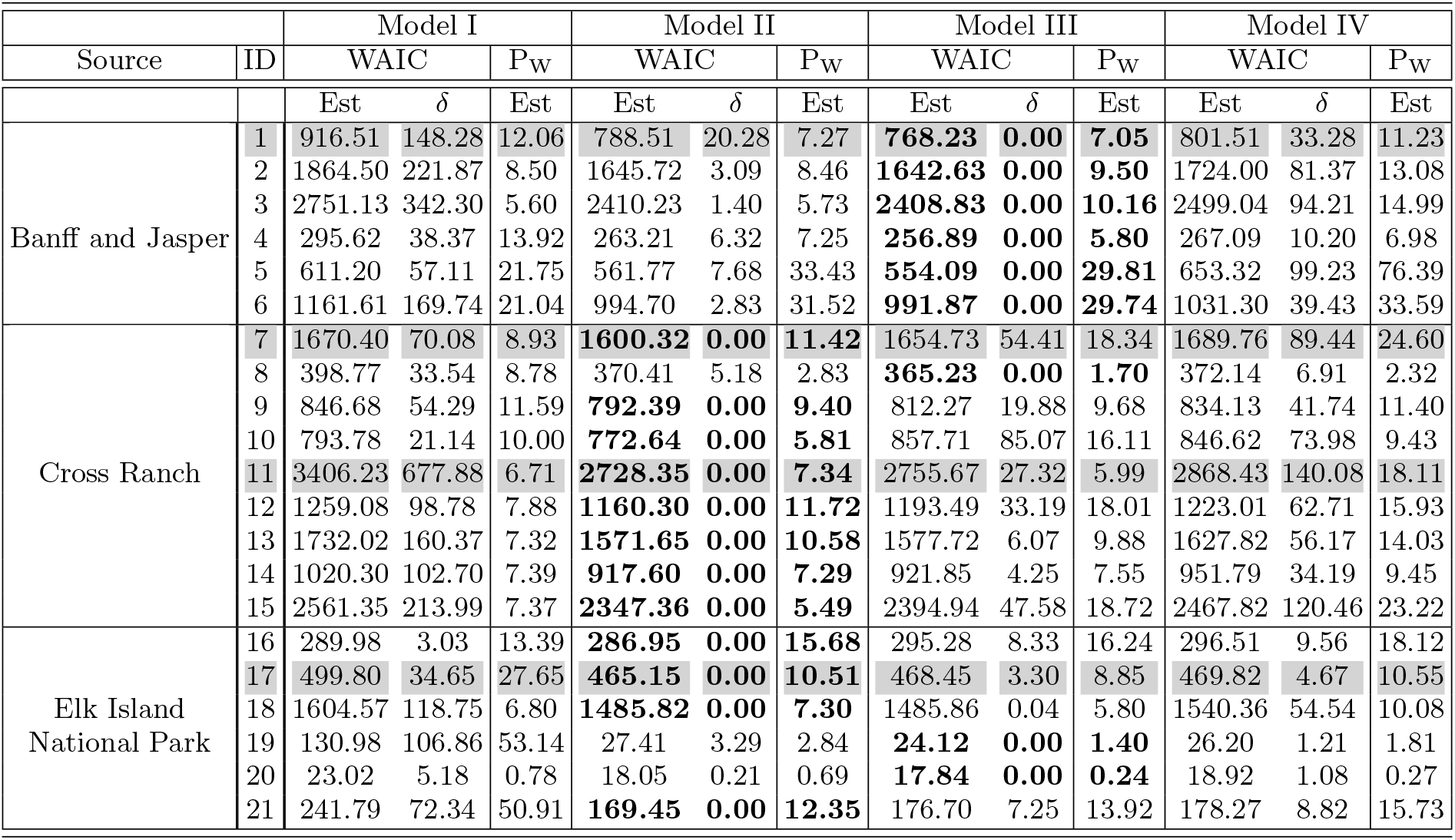
WAIC from the pointwise log-likelihood for each model and each individual. Table shows point estimates (Est) for information criterion WAIC, the effective number of parameters (P_w_) and difference between WAIC’s models as δ. In bold the lowest WAIC for each individual.

To illustrate these general results, we present a closer analysis of the PPC and WAIC for 4 representative individuals that portray different kinds of behaviors on a trajectory. Figure 1 displays the PPC for each model and animals 1, 7, 11 and 17. Table II shows WAIC for all models and the same 4 representative animals in grey). The lowest WAIC between models for each individual is indicated in bold. We denoted as *δ* the difference between the WAIC of each model and the lowest one, and *P_W_* as the effective number of parameters.

**FIG. 1.**
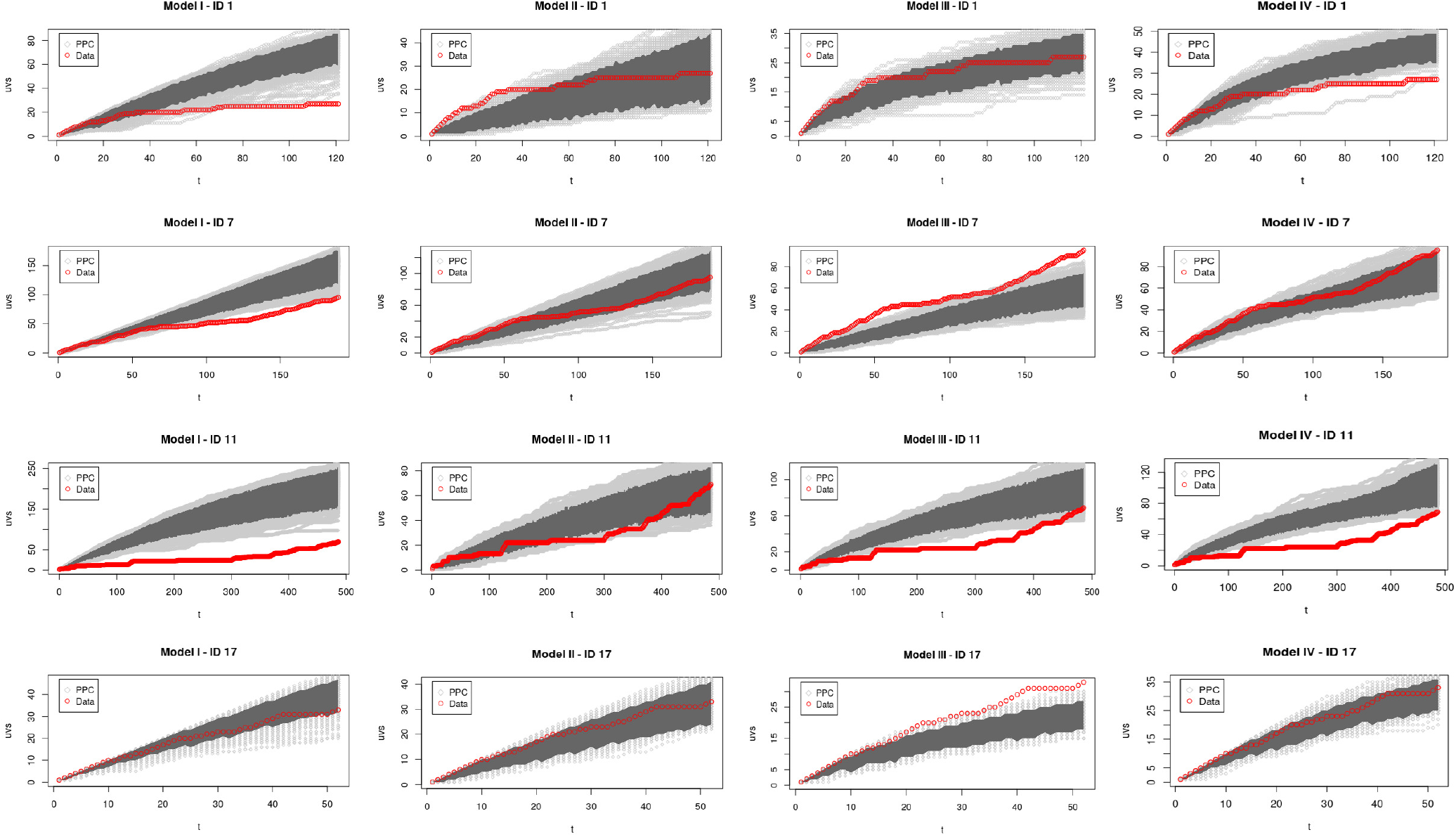
Posterior predictive check (PPC) of models I-IV for elk with ID 1 (1st row), 7 (2nd row), 11 (3rd row) and 17 (4th row). The number of UVS is shown as a function of time. The PPC curves (obtained from sampling the parameters posterior distributions) are in light grey, with the 95% CI in dark grey. The red curves were obtained from the real trajectories. The y-scale for each graph in the same row is different in order to show in a better way which parts of the real trajectory are inside of the CI.

Fig. 1-First Row shows the PPC results for individual 1 from Banff. Model I fitted well only the first steps of the trajectory, indicating that the animal was probably in exploration phase. Later on, the trajectory is no longer contained within Model I credible interval. Model II fitted well the final steps of the trajectory from this animal, suggesting that it followed an exploitation phase with *q* = 0.68 (from here on, all reported parameters values are the mean from their correspond posterior distribution). However, like Model I, neither Model II OR IV described the entire time series. Thus, Model III was the only acceptable model for animal 1, indicating that this particular individual increased its memory use as it explored the environment. In agreement with this finding, Model III had the lowest WAIC for this animal (Table II).

Fig. 1-Second Row displays the PPCs for animal 7 from Cross Ranch. Here, Models I and III fitted well just the first trajectory steps, indicating a exploration phase, but overall, they were not acceptable for animal 7. In contrast, Model IV contained all the observed trajectory within its CI, suggesting that this particular individual increased its memory use as it explored the space and its memory decayed over time. Model II was also acceptable for animal 7, with a constant rate of memory use of *q* = 0.38. Therefore, animal 7 had two possible acceptable models. However, the lowest WAIC for individual 7 was for Model II, and the δ for Model IV was quite large (Table II).

Fig. 1-Third Row corresponds to animal 11, also from Cross Ranch. Models I, III and IV fitted well only the first steps of the observed trajectory. Model II contained within its CI the entire observed data, indicating that this particular animal used its memory at a constant and very high rate (q = 0.80), being most of the time visiting known patches. Table II indicated the lowest WAIC for Model II, confirming the conclusion drawn from the PPC.

Fig. 1-Fourth Row corresponds to animal 17 from Elk Island. Model III fitted well just the first steps of the trajectory and it was not acceptable for this individual. Otherwise Models I, II and IV contained within their respective CI all the observed trajectory. This give us three possible interpretations for animal 17: i) The animal was always in exploratory phase. ii) The individual used its memory at constant rate *q* = 0.25. iii) The animal increased its memory use with time and its memory decayed over the time. Table II shows that Model II actually had the lowest WAIC. Therefore, Model II can be considered as fairly good to describe and predict the trajectory of animal 17.

Table II summarizes models fit to each elk by their source population. Elk from 1 to 6 belong to the Banff and Jasper Source, animals from 7 to 15 to Cross Ranch, and elk from 16 to 21 to Elk Island. Models having the lowest WAIC are bolded. We can see that for all animals from Banff and Jasper, Model III was the best according to WAIC. For animals from Cross Ranch, model II was the best for most of them. And for 66% of elk from Elk Island, Model II was the best. This suggests that animals from different source populations reacted differently to the new environment.

We discuss in the following the different parameters obtained from the fits.

### 3.2. Spatial Parameters

The spatial parameters *α, β, λ* and *κ* are present in all models. The estimated values for these parameters do not vary too much between the four different models. We present here a common interpretation for these parameters. From now on the analysis focuses on the individual-level estimate of each parameter (say *p_j_* (*j* = 1: 21)) as well as on the population-level parameter *p*.

Parameter *α*, which controls the scale of the exponential decay with distance between patches, (see Tables VI-IX of the SM) fluctuated little among individuals and across the four models (0.60 ≤ *α_j_* ≤ 2.57 (km)), with a population average between models of *α* = 1.71. Parameter *β*, which controls the shape of the exponential decay, varied between 0.64 and 1.50 among individuals, with a population average of *β* = 1.09, i.e., close to the exponential shape. These values mean that distance played an important role in patch selection; the animals did not choose patches beyond one or two kilometers from their actual positions (maybe due to the patchiness of the environment) as shown by the posterior curve in Fig. 2-Top. These results highlight the importance of “distance discounting” in movement choices, even when memory was involved.

**FIG. 2.**
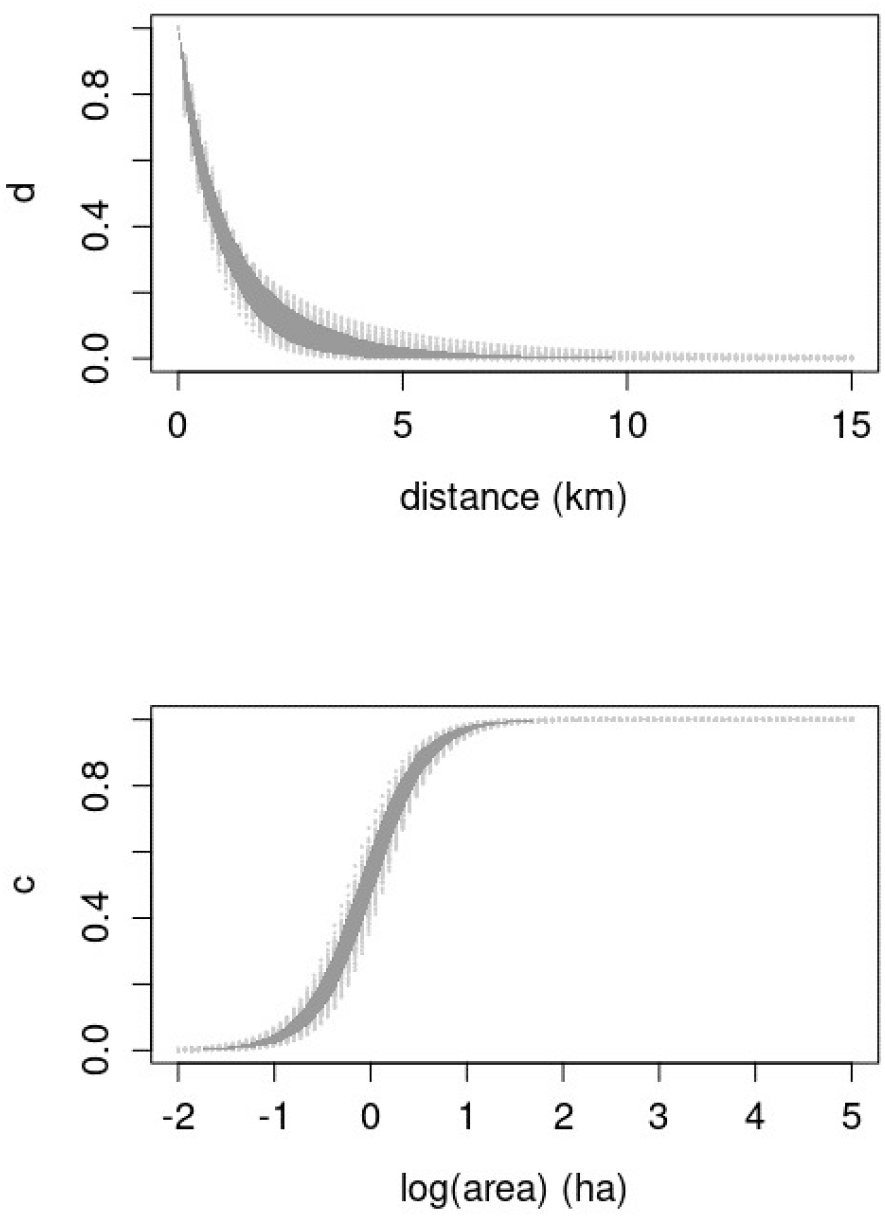
Probability weights (see Eq.1) for choosing a patch as a function of distance (**Top**) and area (**Bottom**) at the population-level for Model I. The curves (obtained from sampling the parameters posterior distributions) are in light grey, with the 95% CI in dark grey. Very similar results were obtained for Models II-IV, see Figs. 6-8 of the SM.

Parameter *λ*, which controls the slope of the logit increase with patch area, also fluctuated little among individuals and models (2.34 ≤ *λ_j_* ≤ 3.70 (ha)), with a population average of λ = 2.90. Whereas parameter *κ*, which controls the intercept of the logit increase, had fluctuations between 0.02 and 0.42 among individuals, and a population average of *κ* = 0.14. We conclude that patch area played a significant role during patch use: the probability increased rapidly for patches of area around 1 ha, and saturated for patches with area greater than 2 ha as shown by the posterior curve in Fig. 2-Bottom.

### 3.3. Memory Use

Figure 3 displays the marginal posterior distributions of the parameter *q*, that defines the probability of memory use in Model II. As mentioned earlier, this model was considered the best for 12 individuals (the ones in blue in Fig. 3). For these individuals *q* had a minimum value of 0.18 and a maximum of 0.80, but most of them had a *q* ≈ 0.5. Hence, according to this model, roughly half of the moves from patch to patch performed by most of the animals are informed by memory, while the other half can be considered as exploratory. For those animals with values of *q* far from 0.5, the trajectories are either dominated by memory (e.g. ID 11) or by exploratory movements (e.g. ID 10 and 21).

**FIG. 3.**
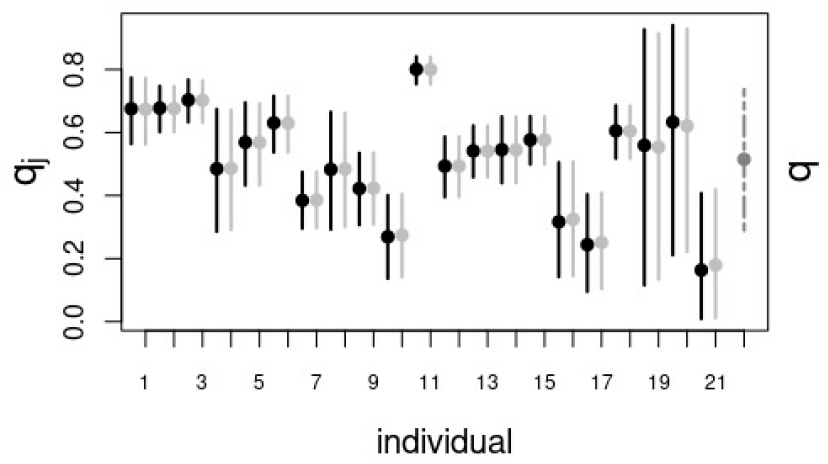
Mean marginal posteriors (points) and 95% CI (vertical lines) for individuals (denoted by ID number) and the population (denoted by ‘pop’) for the parameter *q* of Model II. The black intervals correspond to the results of the first-stage algorithm and the solid light gray intervals correspond to the results of the hierarchical, second-stage algorithm across all animals. Dashed gray interval correspond to the populationlevel.

Model III assumes that *q* grows from zero with the number of UVS at time *t* (*u_t_*), as defined by Eq. (3). Figure 4 displays the marginal posterior distributions of the parameter *ρ*. This parameter defines the number of visited sites needed for the onset of important memory effects. For those individuals for which this Model III was considered the best (in blue on Fig. 4) *ρ* had values between 12.87 and 12.93. Likewise, the shape parameter *ϵ* (Fig. 4) of the exponential ranged between 0.03 and 0.86. Fig. 4-Bottom displays the growth of memory use as a function of **u** at the population-level. Memory use increased rapidly to 0.5 when the unique visited sites were between 5 and 10, before slowly tending to its asymptotic value.

**FIG. 4.**
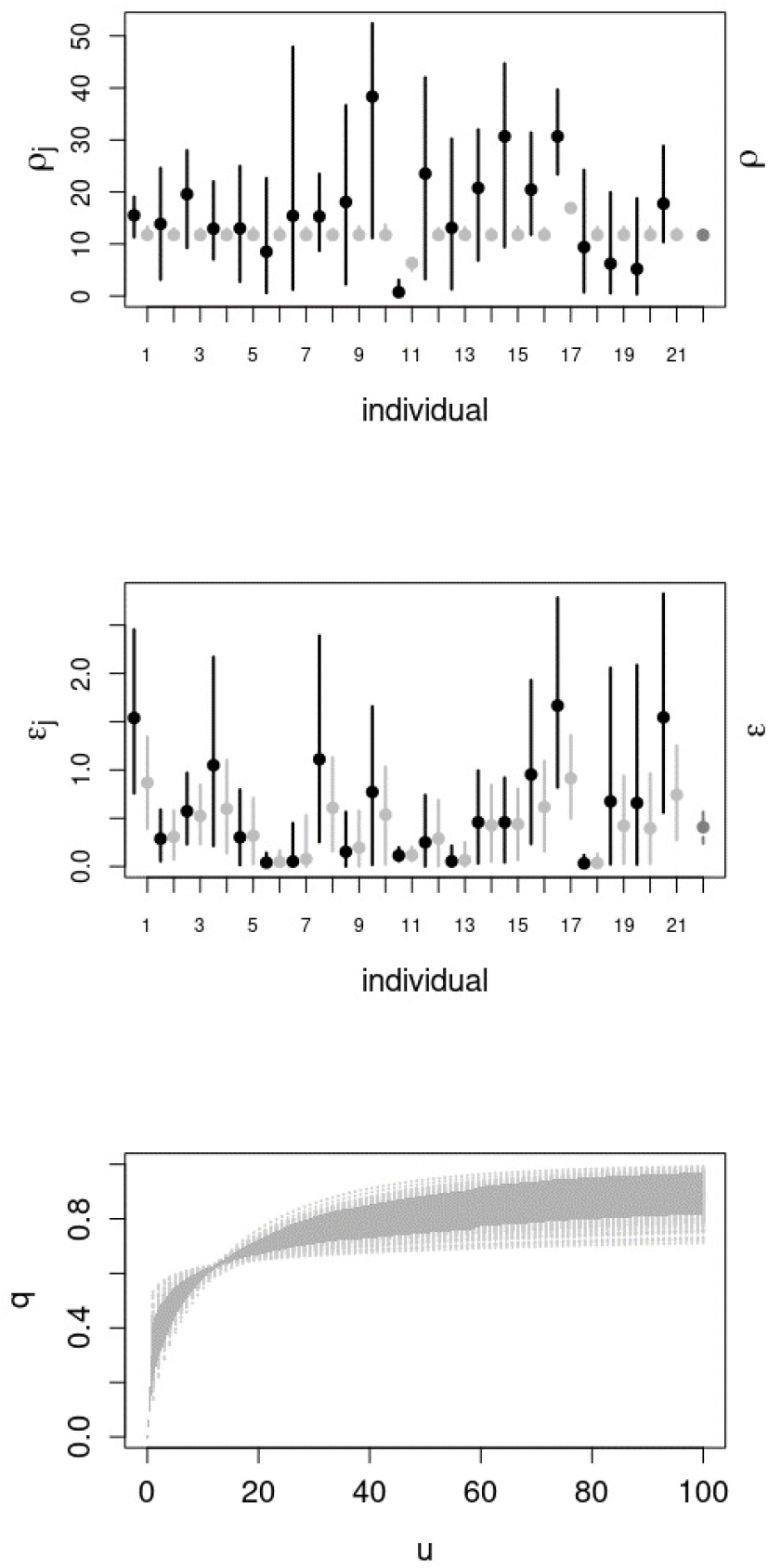
**Top and Center:** Same as Figure 3 but for parameters *ρ* and *ϵ* (resp.) of Model III. **Bottom:** Increase of the probability of memory use *q* as a function of the number of unique visited sites **u** at the population-level. The curves (obtained from sampling parameters from the joint posterior distributions) are in light grey, with the 95% CI in dark grey.

### 3.4. Memory Decay

Model IV takes into account all the assumptions of Model III, with the addition of a decay in memory. Figure 5 displays the marginal posterior distribution of the scale parameter *v* that defines the time scale of memory decay. For the population-level this parameter was estimated as *v* = 10.78 (Fig. 5 in dashed grey line). The shape parameter *θ* was estimated as *θ* = 0.30, thus, memory decayed as a stretched exponential.

**FIG. 5.**
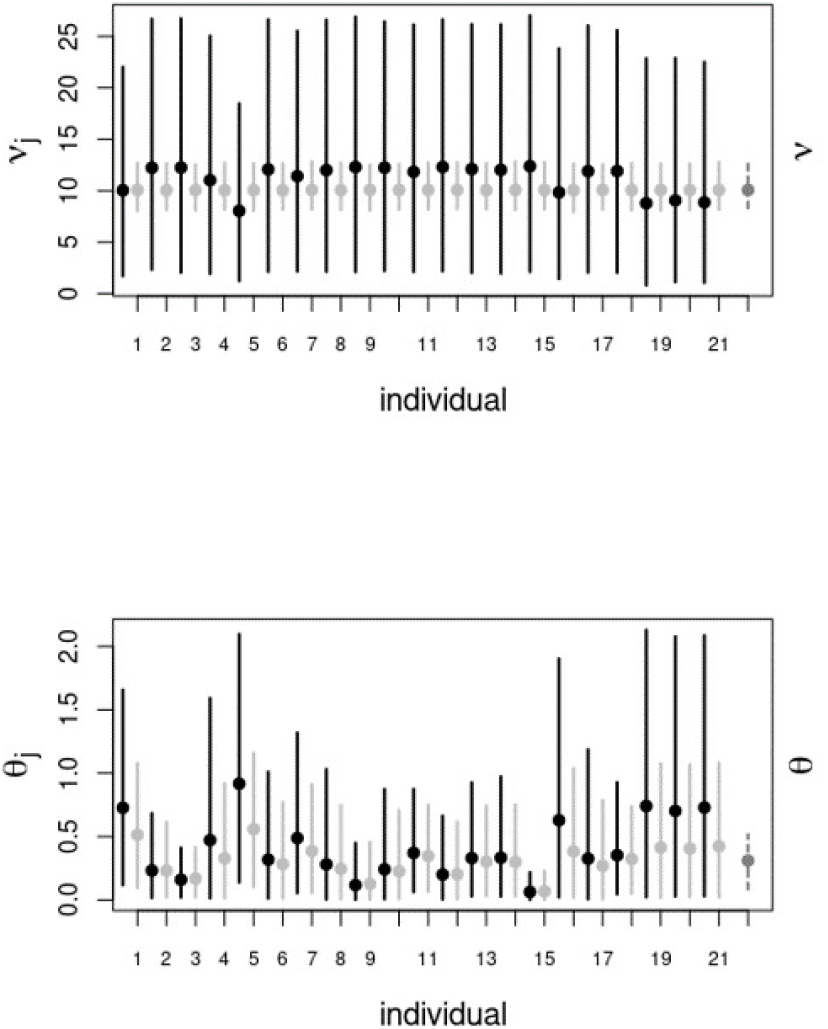
As same as Figure 3 for the parameters *v* (**Top**) and *θ* (**Bottom**) of Model IV.

## 4. DISCUSSION

We have presented four simple models to fit a set of movement data collected in western Canada for 21 elk relocated into a new environment. In a first stage, Bayesian estimates were carried out at the individual-level using Hamiltonian Monte Carlo sampling. A hierarchical analysis was next implemented following the algorithm proposed in (Hooten and Hefley 2019), allowing us to infer how the population as a whole is adapting to a new environment. All the results obtained at the individual-level can be found in the SM. To compare and evaluate these four different models we used two tools, one quantitative the other qualitative. We used, on the one hand the Watanabe Akaike Information Criterion (WAIC) and on the other hand, a Posterior Predictive Check (PPC) based on the number of unique patches visited by the animals. The results obtained by these two tests were in agreement. We found that the trajectories of all animals were far from being described by a memoryless random walk and rather exhibited patterns of recurrent revisits to patches. Our models II and III, that consider an infinite memory capability (with constant and dynamic rate of use, respectively) combined with a linear reinforcement of the visited patches, fitted and predicted well all the trajectories. This is consistent with the results exposed by Wolf et al. (2009) in which, after a thorough statistical study of habitat selection, found that elk have a strong tendency to select the most recently visited locations to forage instead of selecting locations only by their quality. Moreover, the values of our spatial parameters, and the curves that they defined, correspond well with resident elk movement scales reported in (Frair et al. 2005); foraging movements are on the order of hundreds of meters and relocating moves on the order of 1.6 km. Our fourth model, that considered a dynamic use of memory and memory decay, was not considered as the best model for any individual. It thus seems to be too sophisticated for this population over this time period of the data (11 months).

The exploitation-exploration paradigm is a well known concept in ecology. There are several models that have focused on identifying and predicting these two phases from single animal trajectories (Jonsen et al. 2007, Morales et al. 2004) but they are often based on memoryless dynamics and the exploitation-exploration phases are the result of different types of random walks movements. Our Model II is memory-based and the use of memory is represented by a constant parameter q. While the exploration phase is governed by random decisions, the exploitation phase is ruled by the use of memory and the reinforcement learning acquired by experience. These simple assumptions were enough to adequately represent the temporal changes in the number of unique patches visited (Fig. 9 in SM) by twelve animals and therefore to identify the presence of these two phases. It is important to note that those twelve individuals for which Model II was considered the best model, as well as the nine animals for which Model III gave better results, had a high value of *q* (near 1/2 on average). This suggests that these animals used memory intensively, instead of performing pure random walks (which correspond to the limit *q* → 0). A previous study on capuchin monkeys that used a similar model found a value of *q* near 0.12 over a 6-month period (Boyer and Solis-Salas 2014b). In that model the environment was represented as a regular discrete lattice in which each point was a site to visit. The high values of *q* observed here could be explained by frequent decisions to return to high-resource patches or safe places, for instance those where the predation risk (by wolves or humans) is lower. This is also consistent with the scale movement results exposed in (Frair et al. 2005) that shows that elk make use of certain patches and do not explore beyond them, possibly to reduce their mortality rate and predation risk.

We also found that animals from the same source population tended to behave similarly: for most of the animals from Banff and Jasper, Model III was considered the best model, whereas most of the elk from Cross Ranch and Elk Island were best described by Model II. These results might be explained by the experience animals had before translocation: we speculate that if the original environment was similar to the new one or the animal was not naive to predators, the animal relied more heavily on memory as they visited new patches (Model III). Conversely, if the original environment was very different or the animals naive to predators, then the they kept high rates of exploration (Model II). This hypothesis stems from the fact that Banff and Jasper are mountainous with similar kinds of valley meadows as the new habitat, and that the animals were familiar with predators, while Cross Ranch and Elk Island have quite different habitat backgrounds, mostly wide-open areas dominated by agriculture and flatland, respectively, and with animals naive to predators.

It is noteworthy that the model in which memory decays with time (Model IV) was not supported as the best model for any of the animals during the period of this study. This suggests that elk remember very well the places they have visited at least within one year. Similar findings have been reported for other species such as American bison (Merkle et al. 2014), sheep (Gautestad and Mysterud 2005), woodland caribou (Avgar et al. 2015) or chimpanzees (Janmaat et al. 2013). These works reported evidence of long-term or very slowly decaying memory, with individuals having the ability to return sometimes to locations which had not been visited for months, or even years.

The movement trajectories from translocated animals provides a way to study how animals use memory. Our findings are qualitatively consistent with those recently reported by Ranc et al. (2020) on reintroduced roe deer. The movements of those animals were described by a model including both memory and resource preferences, somehow similarly to ours in the memory mode, with a reinforcement that saturated to a limiting value instead of growing linearly as here. Their fitted model was able to predict the dynamics of home range formation observed in roe deer, thus bringing support to the hypothesis that memory constitutes an important mechanism for home range emergence (Börger et al. 2008, Van Moorter et al. 2009). Although not analyzed in detail here, it is very likely that the models that we have fitted would also predict several movement properties indicative of limited space use and home range behaviour in elk but it would be important to have longer observation periods to verify this.

Several extensions would make these models more realistic and complex. For example, the probability of moving from one patch to another could be affected not only by distance and patch area but also by more realistic estimates of movement costs due to topography and other landscape variables such as different habitat types and predation risk between patches. It would also be possible to consider continuous time modeling taking into account the time that an animal spend going from one patch to another, as well as the residence time within patches. Finally, our modelling approach ignored the fact that in a network of patches, nearby patches can compete as possible destinations due to their spatial configuration (Ovaskainen and Cornell 2003). This effect can be approximated by considered all possible ways in which an individual leaving a particular patch can eventually reach another patch in the network, although the computational costs are substantial (Morales et al. 2017).

Our models could capture features of the movement patterns of the study animals with a minimum number of parameters and rather simple dynamical rules. Such simplicity is advantageous if one wishes to apply the same models to other data sets. Particularly, a single parameter *q* quantifies the behaviour of an animal memory-wise, and can serve as a basis for comparisons between individuals or between species. Substantial variations of this parameter among individuals of a same species and in a same environment, as observed here, indicate that the movement strategies employed are quite flexible.

## 5. ACKNOWLEDGEMENTS

AFC thanks Conacyt (Mexico), PAEP-UNAM, and PostDoctoral Scholarship DGAPA-UNAM for financial support. AFC and JMM thanks to M Hooten and T Hefley for fruitful discussion. JMM was supported by a Leverhulme Visiting Professorship (VP2-2018-0630). Financial, logistical and technical support for elk relocation and monitoring was provided by the Alberta Conservation Association, Alberta Fish & Wildlife (Sustainable Resources Division), Banff National Park, Canadian Foundation for Innovation, Elk Island National Park, Jasper National Park, National Science Foundation (Grant 0078130), Rocky Mountain Elk Foundation, Sunpine Forest Products, University of Alberta, and Weyerhaeuser Ltd. Along with Ralph Schmidt and local ranchers.

