## Supplemental Material for "Hierarchical, Memory-based Movement Models for Translocated Elk (*Cervus canadensis*)"

In this supplemental material, we show all the results regarding the posterior parameter fits for each model and each observed animal, at both the individual and hierarchical levels. We considered estimates such as: the mean, standard deviation (SD), quantiles at 2.5% and 97.5%, sample effective size (n_eff), the statistical Rhat and the statistical Geweke to test the convergence of the estimations. For this last metric we computed the associated $p-$value ($p_v$). We also show the WAIC obtained for all models and all individuals. Finally, we show the PPC graphs.

#### INDIVIDUAL LEVEL

The estimates and computations shown here correspond to the fit obtained during the first-stage algorithm exposed briefly in the main paper.

##### Model I

| ID | Parameter | Mean | SD | 2.5% | 97.5% | n_eff | Rhat | ID | Parameter | Mean | SD | 2.5% | 97.5% | n_eff | Rhat |
|---|---|---|---|---|---|---|---|---|---|---|---|---|---|---|---|
| 1 | $\alpha$ | 1.42 | 0.23 | 1.00 | 1.88 | 3126 | 1.00 | 2 | $\alpha$ | 0.72 | 0.08 | 0.56 | 0.86 | 3703 | 1.00 |
| | $\beta$ | 1.13 | 0.12 | 0.91 | 1.38 | 3184 | 1.00 | | $\beta$ | 0.87 | 0.05 | 0.78 | 0.96 | 3773 | 1.00 |
| | $\lambda$ | 4.25 | 0.54 | 3.27 | 5.37 | 5076 | 1.00 | | $\lambda$ | 4.41 | 0.55 | 3.41 | 5.55 | 4850 | 1.00 |
| | $\kappa$ | 0.07 | 0.07 | 0.00 | 0.26 | 6234 | 1.00 | | $\kappa$ | 0.04 | 0.04 | 0.00 | 0.16 | 6174 | 1.00 |
| 3 | $\alpha$ | 1.13 | 0.09 | 0.97 | 1.30 | 4190 | 1.00 | 4 | $\alpha$ | 1.32 | 0.42 | 0.62 | 2.28 | 4185 | 1.00 |
| | $\beta$ | 1.28 | 0.07 | 1.15 | 1.43 | 4317 | 1.00 | | $\beta$ | 0.98 | 0.18 | 0.67 | 1.38 | 3976 | 1.00 |
| | $\lambda$ | 4.43 | 0.52 | 3.50 | 5.50 | 5274 | 1.00 | | $\lambda$ | 2.58 | 0.65 | 1.41 | 4.00 | 5322 | 1.00 |
| | $\kappa$ | 0.03 | 0.03 | 0.00 | 0.12 | 6896 | 1.00 | | $\kappa$ | 0.17 | 0.16 | 0.00 | 0.57 | 5572 | 1.00 |
| 5 | $\alpha$ | 1.17 | 0.39 | 0.54 | 2.06 | 4025 | 1.00 | 6 | $\alpha$ | 1.31 | 0.23 | 0.89 | 1.80 | 4159 | 1.00 |
| | $\beta$ | 0.71 | 0.09 | 0.55 | 0.91 | 4044 | 1.00 | | $\beta$ | 0.86 | 0.07 | 0.73 | 1.01 | 4096 | 1.00 |
| | $\lambda$ | 2.88 | 0.63 | 1.70 | 4.16 | 5636 | 1.00 | | $\lambda$ | 3.70 | 0.57 | 2.66 | 4.87 | 5198 | 1.00 |
| | $\kappa$ | 0.11 | 0.11 | 0.00 | 0.40 | 6228 | 1.00 | | $\kappa$ | 0.06 | 0.06 | 0.00 | 0.24 | 6679 | 1.00 |
| 7 | $\alpha$ | 0.57 | 0.10 | 0.40 | 0.78 | 3580 | 1.00 | 8 | $\alpha$ | 1.41 | 0.25 | 0.93 | 1.93 | 3958 | 1.00 |
| | $\beta$ | 0.64 | 0.03 | 0.58 | 0.72 | 3593 | 1.00 | | $\beta$ | 1.40 | 0.21 | 1.03 | 1.86 | 3925 | 1.00 |
| | $\lambda$ | 3.89 | 0.55 | 2.89 | 5.01 | 5697 | 1.00 | | $\lambda$ | 2.57 | 0.64 | 1.37 | 3.91 | 4790 | 1.00 |
| | $\kappa$ | 0.06 | 0.06 | 0.00 | 0.21 | 6320 | 1.00 | | $\kappa$ | 0.16 | 0.16 | 0.00 | 0.58 | 5285 | 1.00 |
| 9 | $\alpha$ | 1.29 | 0.24 | 0.87 | 1.79 | 3544 | 1.00 | 10 | $\alpha$ | 1.38 | 0.22 | 0.98 | 1.82 | 4109 | 1.00 |
| | $\beta$ | 1.07 | 0.12 | 0.86 | 1.32 | 3539 | 1.00 | | $\beta$ | 1.20 | 0.13 | 0.96 | 1.47 | 4141 | 1.00 |
| | $\lambda$ | 2.97 | 0.61 | 1.84 | 4.23 | 5064 | 1.00 | | $\lambda$ | 2.90 | 0.59 | 1.83 | 4.10 | 5302 | 1.00 |
| | $\kappa$ | 0.07 | 0.07 | 0.00 | 0.27 | 5528 | 1.00 | | $\kappa$ | 0.08 | 0.08 | 0.00 | 0.30 | 6192 | 1.00 |
| 11 | $\alpha$ | 0.82 | 0.05 | 0.71 | 0.92 | 3916 | 1.00 | 12 | $\alpha$ | 1.17 | 0.18 | 0.86 | 1.55 | 4037 | 1.00 |
| | $\beta$ | 1.03 | 0.04 | 0.95 | 1.10 | 3962 | 1.00 | | $\beta$ | 1.01 | 0.08 | 0.86 | 1.18 | 4041 | 1.00 |
| | $\lambda$ | 4.09 | 0.50 | 3.20 | 5.15 | 4653 | 1.00 | | $\lambda$ | 3.41 | 0.55 | 2.41 | 4.57 | 5695 | 1.00 |
| | $\kappa$ | 0.03 | 0.03 | 0.00 | 0.10 | 6565 | 1.00 | | $\kappa$ | 0.05 | 0.05 | 0.00 | 0.19 | 6553 | 1.00 |
| 13 | $\alpha$ | 0.97 | 0.12 | 0.88 | 1.21 | 3658 | 1.00 | 14 | $\alpha$ | 0.75 | 0.16 | 0.47 | 1.09 | 2905 | 1.00 |
| | $\beta$ | 0.87 | 0.05 | 0.84 | 0.98 | 3615 | 1.00 | | $\beta$ | 0.80 | 0.07 | 0.66 | 0.96 | 2786 | 1.00 |
| | $\lambda$ | 4.31 | 0.56 | 3.92 | 5.48 | 5450 | 1.00 | | $\lambda$ | 0.54 | 0.56 | 2.54 | 4.73 | 4602 | 1.00 |
| | $\kappa$ | 0.04 | 0.04 | 0.01 | 0.16 | 5753 | 1.00 | | $\kappa$ | 0.06 | 0.06 | 0.00 | 0.22 | 5066 | 1.00 |
| 15 | $\alpha$ | 1.18 | 0.13 | 0.94 | 1.44 | 3173 | 1.00 | 16 | $\alpha$ | 0.83 | 0.36 | 0.32 | 1.66 | 4077 | 1.00 |
| | $\beta$ | 1.14 | 0.08 | 0.99 | 1.32 | 3210 | 1.00 | | $\beta$ | 0.67 | 0.10 | 0.50 | 0.88 | 3916 | 1.00 |
| | $\lambda$ | 4.81 | 0.52 | 3.86 | 5.88 | 5195 | 1.00 | | $\lambda$ | 2.10 | 0.67 | 0.89 | 3.54 | 3614 | 1.00 |
| | $\kappa$ | 0.03 | 0.03 | 0.00 | 0.11 | 6492 | 1.00 | | $\kappa$ | 0.20 | 0.19 | 0.01 | 0.70 | 6206 | 1.00 |

| ID | Parameter | Mean | SD | 2.5% | 97.5% | n_eff | Rhat | ID | Parameter | Mean | SD | 2.5% | 97.5% | n_eff | Rhat |
|---|---|---|---|---|---|---|---|---|---|---|---|---|---|---|---|
| | $\alpha$ | 1.26 | 0.41 | 0.60 | 2.17 | 3642 | 1.00 | | $\alpha$ | 0.65 | 0.08 | 0.51 | 0.81 | 4422 | 1.00 |
| | $\beta$ | 0.88 | 0.14 | 0.63 | 1.18 | 3638 | 1.00 | | $\beta$ | 0.84 | 0.04 | 0.75 | 0.93 | 4577 | 1.00 |
| 17 | $\lambda$ | 2.25 | 0.64 | 1.12 | 3.59 | 4704 | 1.00 | 18 | $\lambda$ | 3.66 | 0.52 | 2.73 | 4.78 | 5168 | 1.00 |
| | $\kappa$ | 0.15 | 0.15 | 0.00 | 0.54 | 5938 | 1.00 | | $\kappa$ | 0.04 | 0.04 | 0.00 | 0.06 | 6759 | 1.00 |
| | $\alpha$ | 9.92 | 6.17 | 0.83 | 23.95 | 3747 | 1.00 | | $\alpha$ | 5.72 | 5.59 | 0.54 | 20.37 | 2618 | 1.00 |
| | $\beta$ | 0.72 | 0.62 | 0.02 | 2.23 | 3627 | 1.00 | | $\beta$ | 1.04 | 0.67 | 0.08 | 2.57 | 2826 | 1.00 |
| 19 | $\lambda$ | 0.97 | 0.63 | 0.09 | 2.46 | 4309 | 1.00 | 20 | $\lambda$ | 1.25 | 0.78 | 0.13 | 2.74 | 3722 | 1.00 |
| | $\kappa$ | 0.59 | 0.51 | 0.02 | 1.87 | 5222 | 1.00 | | $\kappa$ | 0.57 | 0.49 | 0.02 | 1.83 | 4943 | 1.00 |
| | $\boldsymbol{\alpha}$ | **0.99** | **0.61** | **0.23** | **2.62** | **3536** | **1.00** | | | | | | | | |
| | $\boldsymbol{\beta}$ | **0.67** | **0.14** | **0.44** | **0.98** | **3399** | **1.00** | | | | | | | | |
| **21** | $\boldsymbol{\lambda}$ | **1.90** | **0.66** | **0.73** | **3.28** | **4516** | **1.00** | | | | | | | | |
| | $\boldsymbol{\kappa}$ | **0.31** | **0.29** | **0.01** | **1.09** | **5756** | **1.00** | | | | | | | | |

TABLE I. Parameter estimates for Model I. These results are obtained in the first-stage algorithm at the individual level. The individuals for which this model is the best one, according to the comparison exposed in Table V, are shown in bold.

##### Model II

| **ID** | **Parameter** | **Mean** | **SD** | **2.5%** | **97.5%** | **n_eff** | **Rhat** | **ID** | **Parameter** | **Mean** | **SD** | **2.5%** | **97.5%** | **n_eff** | **Rhat** |
|---|---|---|---|---|---|---|---|---|---|---|---|---|---|---|---|
| | $\alpha$ | 2.56 | 0.35 | 1.92 | 3.27 | 4193 | 1.00 | | $\alpha$ | 1.50 | 0.18 | 1.18 | 1.87 | 4177 | 1.00 |
| | $\beta$ | 1.45 | 0.19 | 1.11 | 1.87 | 4228 | 1.00 | | $\beta$ | 1.01 | 0.06 | 0.89 | 1.14 | 4301 | 1.00 |
| 1 | $\lambda$ | 3.10 | 0.58 | 2.05 | 4.32 | 6145 | 1.00 | 2 | $\lambda$ | 3.27 | 0.61 | 2.18 | 4.50 | 7123 | 1.00 |
| | $\kappa$ | 0.15 | 0.15 | 0.00 | 0.55 | 7500 | 1.00 | | $\kappa$ | 0.09 | 0.09 | 0.00 | 0.34 | 7500 | 1.00 |
| | $q$ | 0.67 | 0.05 | 0.56 | 0.77 | 5902 | 1.00 | | $q$ | 0.68 | 0.04 | 0.60 | 0.75 | 6213 | 1.00 |
| | $\alpha$ | 1.69 | 0.12 | 1.46 | 1.94 | 3977 | 1.00 | | $\alpha$ | 2.58 | 0.71 | 1.36 | 4.14 | 4319 | 1.00 |
| | $\beta$ | 1.50 | 0.10 | 1.32 | 1.70 | 3981 | 1.00 | | $\beta$ | 1.28 | 0.29 | 0.81 | 1.92 | 4300 | 1.00 |
| 3 | $\lambda$ | 3.39 | 0.56 | 2.38 | 4.56 | 6999 | 1.00 | 4 | $\lambda$ | 2.06 | 0.70 | 0.78 | 3.50 | 4954 | 1.00 |
| | $\kappa$ | 0.09 | 0.09 | 0.00 | 0.33 | 7500 | 1.00 | | $\kappa$ | 0.26 | 0.25 | 0.01 | 0.89 | 7500 | 1.00 |
| | $q$ | 0.70 | 0.03 | 0.63 | 0.77 | 6646 | 1.00 | | $q$ | 0.48 | 0.10 | 0.29 | 0.67 | 5592 | 1.00 |
| | $\alpha$ | 3.78 | 1.09 | 2.06 | 6.32 | 4049 | 1.00 | | $\alpha$ | 3.54 | 0.52 | 2.60 | 4.67 | 3821 | 1.00 |
| | $\beta$ | 0.95 | 0.15 | 0.69 | 1.28 | 3619 | 1.00 | | $\beta$ | 1.16 | 0.11 | 0.97 | 1.38 | 3894 | 1.00 |
| 5 | $\lambda$ | 2.18 | 0.65 | 1.04 | 3.54 | 6943 | 1.00 | 6 | $\lambda$ | 2.83 | 0.63 | 1.68 | 4.15 | 6933 | 1.00 |
| | $\kappa$ | 0.20 | 0.19 | 0.00 | 0.71 | 7500 | 1.00 | | $\kappa$ | 0.13 | 0.13 | 0.00 | 0.47 | 7500 | 1.00 |
| | $q$ | 0.57 | 0.07 | 0.43 | 0.69 | 6516 | 1.00 | | $q$ | 0.63 | 0.05 | 0.54 | 0.66 | 6211 | 1.00 |
| | $\boldsymbol{\alpha}$ | **1.38** | **0.23** | **0.89** | **1.79** | **4482** | **1.00** | | $\alpha$ | 2.18 | 0.40 | 1.44 | 3.01 | 4229 | 1.00 |
| | $\boldsymbol{\beta}$ | **0.76** | **0.05** | **0.67** | **0.86** | **4781** | **1.00** | | $\beta$ | 1.72 | 0.34 | 1.16 | 2.49 | 4152 | 1.00 |
| **7** | $\boldsymbol{\lambda}$ | **3.51** | **0.56** | **2.49** | **4.65** | **6664** | **1.00** | 8 | $\lambda$ | 2.09 | 0.69 | 0.83 | 3.46 | 4699 | 1.00 |
| | $\boldsymbol{\kappa}$ | **0.08** | **0.08** | **0.00** | **0.30** | **7500** | **1.00** | | $\kappa$ | 0.23 | 0.22 | 0.01 | 0.83 | 7500 | 1.00 |
| | $\boldsymbol{q}$ | **0.38** | **0.05** | **0.30** | **0.47** | **6346** | **1.00** | | $q$ | 0.48 | 0.10 | 0.29 | 0.66 | 4938 | 1.00 |
| | $\boldsymbol{\alpha}$ | **2.34** | **0.34** | **1.69** | **3.05** | **4502** | **1.00** | | $\boldsymbol{\alpha}$ | **1.82** | **0.26** | **1.35** | **2.35** | **4119** | **1.00** |
| | $\boldsymbol{\beta}$ | **1.45** | **0.19** | **1.12** | **1.87** | **4489** | **1.00** | | $\boldsymbol{\beta}$ | **1.38** | **0.16** | **1.10** | **1.72** | **4026** | **1.00** |
| **9** | $\boldsymbol{\lambda}$ | **2.54** | **0.64** | **1.34** | **3.84** | **5820** | **1.00** | **10** | $\boldsymbol{\lambda}$ | **2.66** | **0.60** | **1.61** | **3.91** | **5645** | **1.00** |
| | $\boldsymbol{\kappa}$ | **0.11** | **0.11** | **0.00** | **0.40** | **7500** | **1.00** | | $\boldsymbol{\kappa}$ | **0.10** | **0.10** | **0.00** | **0.37** | **7500** | **1.00** |
| | $\boldsymbol{q}$ | **0.42** | **0.06** | **0.31** | **0.54** | **6405** | **1.00** | | $\boldsymbol{q}$ | **0.27** | **0.07** | **0.14** | **0.40** | **5914** | **1.00** |
| | $\boldsymbol{\alpha}$ | **2.07** | **0.16** | **1.78** | **2.40** | **4473** | **1.00** | | $\boldsymbol{\alpha}$ | **2.42** | **0.29** | **1.87** | **3.04** | **4414** | **1.00** |
| | $\boldsymbol{\beta}$ | **1.31** | **0.07** | **1.18** | **1.45** | **4565** | **1.00** | | $\boldsymbol{\beta}$ | **1.41** | **0.14** | **1.16** | **1.70** | **4424** | **1.00** |
| **11** | $\boldsymbol{\lambda}$ | **2.93** | **0.59** | **1.85** | **4.17** | **6577** | **1.00** | **12** | $\boldsymbol{\lambda}$ | **2.91** | **0.60** | **1.83** | **4.16** | **6144** | **1.00** |
| | $\boldsymbol{\kappa}$ | **0.08** | **0.08** | **0.00** | **0.31** | **7500** | **1.00** | | $\boldsymbol{\kappa}$ | **0.09** | **0.09** | **0.00** | **0.32** | **7500** | **1.00** |
| | $\boldsymbol{q}$ | **0.80** | **0.02** | **0.76** | **0.84** | **6445** | **1.00** | | $\boldsymbol{q}$ | **0.49** | **0.05** | **0.40** | **0.59** | **5862** | **1.00** |
| | $\boldsymbol{\alpha}$ | **1.88** | **0.21** | **1.50** | **2.30** | **4951** | **1.00** | | $\alpha$ | 1.59 | 0.29 | 1.09 | 2.21 | 4595 | 1.00 |
| | $\boldsymbol{\beta}$ | **1.07** | **0.07** | **0.94** | **1.21** | **5013** | **1.00** | | $\beta$ | 0.98 | 0.10 | 0.80 | 1.20 | 4560 | 1.00 |
| **13** | $\boldsymbol{\lambda}$ | **3.44** | **0.61** | **2.30** | **4.70** | **5987** | **1.00** | 14 | $\lambda$ | 2.90 | 0.59 | 1.83 | 4.16 | 6051 | 1.00 |
| | $\boldsymbol{\kappa}$ | **0.08** | **0.08** | **0.00** | **0.31** | **7500** | **1.00** | | $\kappa$ | 0.10 | 0.10 | 0.00 | 0.39 | 7500 | 1.00 |
| | $\boldsymbol{q}$ | **0.54** | **0.04** | **0.46** | **0.63** | **5781** | **1.00** | | $q$ | 0.55 | 0.05 | 0.44 | 0.65 | 6290 | 1.00 |
| | $\alpha$ | 1.61 | 0.14 | 1.35 | 1.90 | 4688 | 1.00 | | $\boldsymbol{\alpha}$ | **2.05** | **0.88** | **0.77** | **4.13** | **3998** | **1.00** |
| | $\beta$ | 1.30 | 0.09 | 1.13 | 1.49 | 4729 | 1.00 | | $\boldsymbol{\beta}$ | **0.83** | **0.15** | **0.58** | **1.17** | **3800** | **1.00** |
| 15 | $\lambda$ | 4.02 | 0.56 | 2.98 | 5.20 | 5989 | 1.00 | **16** | $\boldsymbol{\lambda}$ | **1.86** | **0.67** | **0.70** | **3.24** | **5959** | **1.00** |
| | $\kappa$ | 0.06 | 0.06 | 0.00 | 0.22 | 7500 | 1.00 | | $\boldsymbol{\kappa}$ | **0.25** | **0.24** | **0.01** | **0.90** | **7500** | **1.00** |
| | $q$ | 0.58 | 0.04 | 0.50 | 0.65 | 5758 | 1.00 | | $\boldsymbol{q}$ | **0.32** | **0.09** | **0.14** | **0.51** | **6488** | **1.00** |

| ID | Parameter | Mean | SD | 2.5% | 97.5% | n_eff | Rhat | ID | Parameter | Mean | SD | 2.5% | 97.5% | n_eff | Rhat |
|---|---|---|---|---|---|---|---|---|---|---|---|---|---|---|---|
| 17 | $\alpha$ | 1.74 | 0.47 | 0.91 | 2.77 | 4817 | 1.00 | 18 | $\alpha$ | 1.43 | 0.19 | 1.09 | 1.85 | 4221 | 1.00 |
| | $\beta$ | 0.99 | 0.16 | 0.72 | 1.33 | 4919 | 1.00 | | $\beta$ | 0.99 | 0.07 | 0.86 | 1.13 | 4551 | 1.00 |
| | $\lambda$ | 2.09 | 0.65 | 0.93 | 3.44 | 6230 | 1.00 | | $\lambda$ | 3.07 | 0.57 | 2.03 | 4.25 | 6083 | 1.00 |
| | $\kappa$ | 0.18 | 0.18 | 0.01 | 0.67 | 7500 | 1.00 | | $\kappa$ | 0.08 | 0.08 | 0.00 | 0.28 | 7500 | 1.00 |
| | $q$ | 0.25 | 0.08 | 0.10 | 0.40 | 5090 | 1.00 | | $q$ | 0.61 | 0.04 | 0.52 | 0.69 | 5730 | 1.00 |
| 19 | $\alpha$ | 11.24 | 6.07 | 1.80 | 25.05 | 4883 | 1.00 | 20 | $\alpha$ | 9.07 | 5.97 | 1.22 | 23.09 | 6179 | 1.00 |
| | $\beta$ | 0.92 | 0.64 | 0.04 | 2.39 | 4169 | 1.00 | | $\beta$ | 1.05 | 0.60 | 0.15 | 2.43 | 7500 | 1.00 |
| | $\lambda$ | 0.87 | 0.63 | 0.04 | 2.35 | 5030 | 1.00 | | $\lambda$ | 0.91 | 0.63 | 0.04 | 2.37 | 4219 | 1.00 |
| | $\kappa$ | 0.71 | 0.56 | 0.03 | 2.08 | 7500 | 1.00 | | $\kappa$ | 0.74 | 0.57 | 0.03 | 2.14 | 6296 | 1.00 |
| | $q$ | 0.56 | 0.22 | 0.11 | 0.93 | 5489 | 1.00 | | $q$ | 0.63 | 0.20 | 0.20 | 0.95 | 5559 | 1.00 |
| 21 | $\alpha$ | 1.39 | 0.88 | 0.30 | 3.64 | 3852 | 1.00 | | | | | | | | |
| | $\beta$ | 0.71 | 0.16 | 0.46 | 1.07 | 3956 | 1.00 | | | | | | | | |
| | $\lambda$ | 1.80 | 0.65 | 0.63 | 3.17 | 4380 | 1.00 | | | | | | | | |
| | $\kappa$ | 0.33 | 0.31 | 0.01 | 1.10 | 6667 | 1.00 | | | | | | | | |
| | $q$ | 0.16 | 0.11 | 0.01 | 0.41 | 5163 | 1.00 | | | | | | | | |
| | $1-q$ | 0.84 | 0.11 | 0.59 | 0.99 | 5163 | 1.00 | | | | | | | | |

TABLE II. Parameter estimates for Model II. These results are obtained in the first-stage algorithm at the individual level. The individuals for which this model is the best one, according to the comparison exposed in Table V, are shown in bold.

#### Model III

| ID | Parameter | Mean | SD | 2.5% | 97.5% | n_eff | Rhat | ID | Parameter | Mean | SD | 2.5% | 97.5% | n_eff | Rhat |
|---|---|---|---|---|---|---|---|---|---|---|---|---|---|---|---|
| **1** | $\boldsymbol{\alpha}$ | **2.66** | **0.36** | **2.00** | **3.39** | **8490** | **1.00** | **2** | $\boldsymbol{\alpha}$ | **1.53** | **0.18** | **1.20** | **1.89** | **7775** | **1.00** |
| | $\boldsymbol{\beta}$ | **1.47** | **0.20** | **1.12** | **1.90** | **8665** | **1.00** | | $\boldsymbol{\beta}$ | **1.02** | **0.06** | **0.90** | **1.15** | **7843** | **1.00** |
| | $\boldsymbol{\lambda}$ | **2.93** | **0.59** | **1.86** | **4.17** | **10898** | **1.00** | | $\boldsymbol{\lambda}$ | **3.19** | **0.60** | **2.10** | **4.45** | **11755** | **1.00** |
| | $\boldsymbol{\kappa}$ | **0.17** | **0.16** | **0.00** | **0.60** | **15224** | **1.00** | | $\boldsymbol{\kappa}$ | **0.09** | **0.10** | **0.00** | **0.35** | **13351** | **1.00** |
| | $\boldsymbol{\rho}$ | **15.53** | **1.99** | **11.26** | **19.12** | **6625** | **1.00** | | $\boldsymbol{\rho}$ | **13.87** | **5.48** | **3.30** | **24.51** | **7389** | **1.00** |
| | $\boldsymbol{\epsilon}$ | **1.54** | **0.44** | **0.75** | **2.47** | **8186** | **1.00** | | $\boldsymbol{\epsilon}$ | **0.29** | **0.14** | **0.05** | **0.59** | **6728** | **1.00** |
| **3** | $\boldsymbol{\alpha}$ | **1.71** | **0.12** | **1.48** | **1.96** | **9258** | **1.00** | **4** | $\boldsymbol{\alpha}$ | **2.69** | **0.72** | **1.45** | **4.29** | **8750** | **1.00** |
| | $\boldsymbol{\beta}$ | **1.50** | **0.10** | **1.32** | **1.70** | **9153** | **1.00** | | $\boldsymbol{\beta}$ | **1.30** | **0.30** | **0.83** | **1.98** | **8548** | **1.00** |
| | $\boldsymbol{\lambda}$ | **3.31** | **0.57** | **2.27** | **4.52** | **12103** | **1.00** | | $\boldsymbol{\lambda}$ | **1.99** | **0.70** | **0.72** | **3.45** | **10121** | **1.00** |
| | $\boldsymbol{\kappa}$ | **0.10** | **0.10** | **0.00** | **0.36** | **15339** | **1.00** | | $\boldsymbol{\kappa}$ | **0.27** | **0.26** | **0.01** | **0.96** | **14398** | **1.00** |
| | $\boldsymbol{\rho}$ | **19.57** | **4.71** | **9.23** | **28.01** | **6596** | **1.00** | | $\boldsymbol{\rho}$ | **12.97** | **3.70** | **7.08** | **22.17** | **8028** | **1.00** |
| | $\boldsymbol{\epsilon}$ | **0.57** | **0.19** | **0.22** | **0.97** | **7522** | **1.00** | | $\boldsymbol{\epsilon}$ | **1.05** | **0.50** | **0.22** | **2.17** | **9182** | **1.00** |
| 5 | $\alpha$ | 3.83 | 1.08 | 2.08 | 6.27 | 7535 | 1.00 | **6** | $\boldsymbol{\alpha}$ | **3.55** | **0.52** | **2.63** | **4.65** | **7909** | **1.00** |
| | $\beta$ | 0.95 | 0.15 | 0.69 | 1.28 | 7200 | 1.00 | | $\boldsymbol{\beta}$ | **1.16** | **0.11** | **0.96** | **1.38** | **7788** | **1.00** |
| | $\lambda$ | 2.14 | 0.66 | 0.99 | 3.52 | 10322 | 1.00 | | $\boldsymbol{\lambda}$ | **2.82** | **0.63** | **1.69** | **4.10** | **10990** | **1.00** |
| | $\kappa$ | 0.20 | 0.19 | 0.01 | 0.72 | 14711 | 1.00 | | $\boldsymbol{\kappa}$ | **0.13** | **0.13** | **0.00** | **0.47** | **15281** | **1.00** |
| | $\rho$ | 13.01 | 5.55 | 2.76 | 24.81 | 7476 | 1.00 | | $\boldsymbol{\rho}$ | **8.73** | **5.95** | **0.58** | **22.78** | **13169** | **1.00** |
| | $\epsilon$ | 0.30 | 0.21 | 0.01 | 0.79 | 7244 | 1.00 | | $\boldsymbol{\epsilon}$ | **0.04** | **0.04** | **0.00** | **0.14** | **14605** | **1.00** |
| 7 | $\alpha$ | 1.52 | 0.25 | 1.08 | 2.04 | 6916 | 1.00 | **8** | $\boldsymbol{\alpha}$ | **2.28** | **0.41** | **1.52** | **3.13** | **7960** | **1.00** |
| | $\beta$ | 0.79 | 0.05 | 0.70 | 0.89 | 6808 | 1.00 | | $\boldsymbol{\beta}$ | **1.75** | **0.36** | **1.17** | **2.57** | **7870** | **1.00** |
| | $\lambda$ | 3.41 | 0.59 | 2.32 | 4.62 | 10012 | 1.00 | | $\boldsymbol{\lambda}$ | **1.99** | **0.68** | **0.76** | **3.42** | **9750** | **1.00** |
| | $\kappa$ | 0.09 | 0.09 | 0.00 | 0.31 | 10702 | 1.00 | | $\boldsymbol{\kappa}$ | **0.24** | **0.23** | **0.01** | **0.84** | **13553** | **1.00** |
| | $\rho$ | 15.62 | 11.76 | 1.18 | 48.99 | 3073 | 1.00 | | $\boldsymbol{\rho}$ | **15.32** | **3.69** | **8.73** | **23.64** | **8089** | **1.00** |
| | $\epsilon$ | 0.05 | 0.12 | 0.00 | 0.46 | 2006 | 1.01 | | $\boldsymbol{\epsilon}$ | **1.10** | **0.54** | **0.24** | **2.36** | **9359** | **1.00** |
| 9 | $\alpha$ | 2.46 | 0.34 | 1.82 | 3.15 | 6768 | 1.00 | 10 | $\alpha$ | 1.97 | 0.26 | 1.47 | 2.51 | 6496 | 1.00 |
| | $\beta$ | 1.50 | 0.20 | 1.15 | 1.92 | 7160 | 1.00 | | $\beta$ | 1.44 | 0.16 | 1.14 | 1.78 | 6551 | 1.00 |
| | $\lambda$ | 2.46 | 0.64 | 1.30 | 3.79 | 9995 | 1.00 | | $\lambda$ | 2.55 | 0.62 | 1.43 | 3.86 | 7972 | 1.00 |
| | $\kappa$ | 0.12 | 0.13 | 0.00 | 0.46 | 11871 | 1.00 | | $\kappa$ | 0.11 | 0.11 | 0.00 | 0.42 | 10156 | 1.00 |
| | $\rho$ | 18.17 | 9.34 | 2.33 | 36.58 | 5665 | 1.00 | | $\rho$ | 38.28 | 9.71 | 10.86 | 52.38 | 1781 | 1.01 |
| | $\epsilon$ | 0.15 | 0.16 | 0.00 | 0.57 | 7317 | 1.00 | | $\epsilon$ | 0.77 | 0.42 | 0.02 | 1.66 | 3412 | 1.00 |

| ID | Parameter | Mean | SD | 2.5% | 97.5% | n_eff | Rhat | ID | Parameter | Mean | SD | 2.5% | 97.5% | n_eff | Rhat |
|---|---|---|---|---|---|---|---|---|---|---|---|---|---|---|---|
| 11 | $\alpha$ | 2.04 | 0.16 | 1.75 | 2.36 | 9338 | 1.00 | 12 | $\alpha$ | 2.55 | 0.30 | 1.99 | 3.18 | 6418 | 1.00 |
| | $\beta$ | 1.30 | 0.07 | 1.17 | 1.44 | 9190 | 1.00 | | $\beta$ | 1.45 | 0.15 | 1.19 | 1.77 | 6360 | 1.00 |
| | $\lambda$ | 2.94 | 0.58 | 1.89 | 4.13 | 12515 | 1.00 | | $\lambda$ | 2.83 | 0.61 | 1.74 | 4.12 | 9379 | 1.00 |
| | $\kappa$ | 0.08 | 0.08 | 0.00 | 0.30 | 15294 | 1.00 | | $\kappa$ | 0.09 | 0.09 | 0.00 | 0.35 | 13000 | 1.00 |
| | $\rho$ | 0.75 | 0.84 | 0.01 | 3.12 | 10498 | 1.00 | | $\rho$ | 23.61 | 10.73 | 3.27 | 42.07 | 3303 | 1.00 |
| | $\epsilon$ | 0.11 | 0.04 | 0.06 | 0.20 | 8253 | 1.00 | | $\epsilon$ | 0.25 | 0.21 | 0.00 | 0.73 | 4152 | 1.00 |
| 13 | $\alpha$ | 1.98 | 0.21 | 1.59 | 2.41 | 8572 | 1.00 | **14** | $\boldsymbol{\alpha}$ | **1.66** | **0.29** | **1.14** | **2.27** | **7723** | **1.00** |
| | $\beta$ | 1.08 | 0.07 | 0.95 | 1.23 | 8550 | 1.00 | | $\boldsymbol{\beta}$ | **0.99** | **0.10** | **0.80** | **1.20** | **7701** | **1.00** |
| | $\lambda$ | 3.33 | 0.61 | 2.21 | 4.60 | 11566 | 1.00 | | $\boldsymbol{\lambda}$ | **2.83** | **0.59** | **1.79** | **4.07** | **9873** | **1.00** |
| | $\kappa$ | 0.09 | 0.09 | 0.00 | 0.33 | 13518 | 1.00 | | $\boldsymbol{\kappa}$ | **0.11** | **0.11** | **0.00** | **0.41** | **12097** | **1.00** |
| | $\rho$ | 13.04 | 7.77 | 1.26 | 30.24 | 9648 | 1.00 | | $\boldsymbol{\rho}$ | **20.89** | **6.17** | **6.81** | **32.20** | **4391** | **1.00** |
| | $\epsilon$ | 0.05 | 0.06 | 0.00 | 0.22 | 9970 | 1.00 | | $\boldsymbol{\epsilon}$ | **0.46** | **0.25** | **0.03** | **0.99** | **4725** | **1.00** |
| **15** | $\boldsymbol{\alpha}$ | **1.66** | **0.14** | **1.39** | **1.95** | **7170** | **1.00** | 16 | $\alpha$ | 2.15 | 0.90 | 0.82 | 4.31 | 8617 | 1.00 |
| | $\boldsymbol{\beta}$ | **1.32** | **0.10** | **1.14** | **1.51** | **7210** | **1.00** | | $\beta$ | 0.84 | 0.16 | 0.58 | 1.19 | 8477 | 1.00 |
| | $\boldsymbol{\lambda}$ | **3.88** | **0.56** | **2.85** | **5.03** | **9352** | **1.00** | | $\lambda$ | 1.81 | 0.67 | 0.66 | 3.22 | 12426 | 1.00 |
| | $\boldsymbol{\kappa}$ | **0.07** | **0.07** | **0.00** | **0.27** | **13499** | **1.00** | | $\kappa$ | 0.25 | 0.24 | 0.01 | 0.91 | 16531 | 1.00 |
| | $\boldsymbol{\rho}$ | **30.66** | **8.67** | **9.48** | **44.70** | **3101** | **1.00** | | $\rho$ | 20.52 | 4.92 | 11.82 | 31.52 | 9499 | 1.00 |
| | $\boldsymbol{\epsilon}$ | **0.46** | **0.23** | **0.04** | **0.91** | **4290** | **1.00** | | $\epsilon$ | 0.96 | 0.43 | 0.24 | 1.95 | 10800 | 1.00 |
| **17** | $\boldsymbol{\alpha}$ | **1.87** | **0.48** | **1.04** | **2.91** | **8819** | **1.00** | **18** | $\boldsymbol{\alpha}$ | **1.46** | **0.19** | **1.12** | **1.86** | **8451** | **1.00** |
| | $\boldsymbol{\beta}$ | **1.02** | **0.16** | **0.74** | **1.35** | **8919** | **1.00** | | $\boldsymbol{\beta}$ | **0.99** | **0.07** | **0.87** | **1.13** | **8577** | **1.00** |
| | $\boldsymbol{\lambda}$ | **2.02** | **0.66** | **0.86** | **3.41** | **12324** | **1.00** | | $\boldsymbol{\lambda}$ | **3.04** | **0.57** | **1.99** | **4.23** | **11963** | **1.00** |
| | $\boldsymbol{\kappa}$ | **0.20** | **0.19** | **0.00** | **0.71** | **15471** | **1.00** | | $\boldsymbol{\kappa}$ | **0.08** | **0.08** | **0.00** | **0.29** | **14408** | **1.00** |
| | $\boldsymbol{\rho}$ | **30.74** | **4.15** | **23.37** | **39.71** | **11688** | **1.00** | | $\boldsymbol{\rho}$ | **9.43** | **6.28** | **0.65** | **24.15** | **11308** | **1.00** |
| | $\boldsymbol{\epsilon}$ | **1.66** | **0.50** | **0.81** | **2.75** | **12708** | **1.00** | | $\boldsymbol{\epsilon}$ | **0.03** | **0.03** | **0.00** | **0.11** | **11678** | **1.00** |
| **19** | $\boldsymbol{\alpha}$ | **11.17** | **5.99** | **1.97** | **25.02** | **10048** | **1.00** | **20** | $\boldsymbol{\alpha}$ | **9.04** | **5.94** | **1.25** | **23.01** | **11610** | **1.00** |
| | $\boldsymbol{\beta}$ | **0.90** | **0.65** | **0.03** | **2.40** | **9076** | **1.00** | | $\boldsymbol{\beta}$ | **1.03** | **0.61** | **0.11** | **2.43** | **9758** | **1.00** |
| | $\boldsymbol{\lambda}$ | **0.88** | **0.61** | **0.05** | **2.31** | **12080** | **1.00** | | $\boldsymbol{\lambda}$ | **0.92** | **0.65** | **0.05** | **2.42** | **1139** | **1.00** |
| | $\boldsymbol{\kappa}$ | **0.71** | **0.56** | **0.02** | **2.07** | **12244** | **1.00** | | $\boldsymbol{\kappa}$ | **0.73** | **0.57** | **0.03** | **2.11** | **11317** | **1.00** |
| | $\boldsymbol{\rho}$ | **6.24** | **5.32** | **0.54** | **20.08** | **11374** | **1.00** | | $\boldsymbol{\rho}$ | **5.12** | **4.98** | **0.39** | **18.60** | **9024** | **1.00** |
| | $\boldsymbol{\epsilon}$ | **0.67** | **0.54** | **0.02** | **2.04** | **9743** | **1.00** | | $\boldsymbol{\epsilon}$ | **0.66** | **0.56** | **0.02** | **2.09** | **9319** | **1.00** |
| 21 | $\alpha$ | 1.72 | 1.05 | 0.43 | 4.39 | 8474 | 1.00 | | | | | | | | |
| | $\beta$ | 0.75 | 0.17 | 0.48 | 1.13 | 8842 | 1.00 | | | | | | | | |
| | $\lambda$ | 1.75 | 0.67 | 0.55 | 3.17 | 11995 | 1.00 | | | | | | | | |
| | $\kappa$ | 0.35 | 0.32 | 0.01 | 1.19 | 15010 | 1.00 | | | | | | | | |
| | $\rho$ | 17.77 | 4.75 | 10.49 | 29.01 | 11078 | 1.00 | | | | | | | | |
| | $\epsilon$ | 1.54 | 0.59 | 0.55 | 2.83 | 12074 | 1.00 | | | | | | | | |

TABLE III. Parameter estimates for Model III. These results are obtained in the first-stage algorithm at the individual level. The individuals for which this model is the best one, according to the comparison exposed in Table V, are shown in bold.

**Model IV**

| ID | Parameter | Mean | SD | 2.5% | 97.5% | n_eff | Rhat | ID | Parameter | Mean | SD | 2.5% | 97.5% | n_eff | Rhat |
|---|---|---|---|---|---|---|---|---|---|---|---|---|---|---|---|
| 1 | $\alpha$ | 2.89 | 0.38 | 2.19 | 3.67 | 8147 | 1.00 | 2 | $\alpha$ | 1.35 | 0.17 | 1.04 | 1.70 | 7078 | 1.00 |
| | $\beta$ | 1.62 | 0.22 | 1.24 | 2.09 | 8233 | 1.00 | | $\beta$ | 1.01 | 0.06 | 0.89 | 1.13 | 7318 | 1.00 |
| | $\lambda$ | 3.43 | 0.57 | 2.38 | 4.60 | 12288 | 1.00 | | $\lambda$ | 3.58 | 0.58 | 2.51 | 4.79 | 11313 | 1.00 |
| | $\kappa$ | 0.15 | 0.14 | 0.00 | 0.53 | 15610 | 1.00 | | $\kappa$ | 0.08 | 0.08 | 0.00 | 0.29 | 14087 | 1.00 |
| | $\rho$ | 19.81 | 3.10 | 13.55 | 25.99 | 9191 | 1.00 | | $\rho$ | 25.42 | 9.74 | 4.67 | 41.34 | 3356 | 1.00 |
| | $\epsilon$ | 1.07 | 0.40 | 0.35 | 1.92 | 10002 | 1.00 | | $\epsilon$ | 0.32 | 0.23 | 0.01 | 0.80 | 4113 | 1.00 |
| | $\nu$ | 10.06 | 6.50 | 1.65 | 22.05 | 7004 | 1.00 | | $\nu$ | 12.36 | 6.40 | 2.26 | 26.83 | 8262 | 1.00 |
| | $\theta$ | 0.73 | 0.41 | 0.12 | 1.68 | 7412 | 1.00 | | $\theta$ | 0.24 | 0.18 | 0.02 | 0.70 | 8135 | 1.00 |
| 3 | $\alpha$ | 1.60 | 0.12 | 1.38 | 1.84 | 6953 | 1.00 | 4 | $\alpha$ | 2.45 | 0.73 | 1.22 | 4.07 | 8155 | 1.00 |
| | $\beta$ | 1.50 | 0.09 | 1.33 | 1.69 | 7432 | 1.00 | | $\beta$ | 1.24 | 0.28 | 0.79 | 1.89 | 8459 | 1.00 |
| | $\lambda$ | 3.73 | 0.54 | 2.76 | 4.86 | 10415 | 1.00 | | $\lambda$ | 2.12 | 0.72 | 0.80 | 3.61 | 11860 | 1.00 |
| | $\kappa$ | 0.07 | 0.07 | 0.00 | 0.26 | 13018 | 1.00 | | $\kappa$ | 0.26 | 0.25 | 0.01 | 0.91 | 16563 | 1.00 |
| | $\rho$ | 30.37 | 11.28 | 4.62 | 46.26 | 2132 | 1.00 | | $\rho$ | 15.58 | 4.40 | 8.53 | 25.88 | 8444 | 1.00 |
| | $\epsilon$ | 0.40 | 0.28 | 0.01 | 0.98 | 2892 | 1.00 | | $\epsilon$ | 1.06 | 0.49 | 0.24 | 2.13 | 9579 | 1.00 |
| | $\nu$ | 12.31 | 6.49 | 1.99 | 26.89 | 6787 | 1.00 | | $\nu$ | 11.02 | 6.10 | 1.93 | 25.24 | 13294 | 1.00 |
| | $\theta$ | 0.16 | 0.10 | 0.02 | 0.41 | 7047 | 1.00 | | $\theta$ | 0.47 | 0.43 | 0.01 | 1.55 | 1383 | 1.00 |
| **5** | $\boldsymbol{\alpha}$ | **5.52** | **1.70** | **2.43** | **9.42** | **8729** | **1.00** | 6 | $\alpha$ | 3.30 | 0.49 | 2.43 | 4.34 | 9169 | 1.00 |
| | $\boldsymbol{\beta}$ | **1.09** | **0.20** | **0.76** | **1.55** | **8665** | **1.00** | | $\beta$ | 1.16 | 0.10 | 0.97 | 1.37 | 9449 | 1.00 |
| | $\boldsymbol{\lambda}$ | **2.22** | **0.65** | **1.06** | **3.56** | **11969** | **1.00** | | $\lambda$ | 2.98 | 0.63 | 1.80 | 4.27 | 12391 | 1.00 |
| | $\boldsymbol{\kappa}$ | **0.21** | **0.19** | **0.01** | **0.71** | **14948** | **1.00** | | $\kappa$ | 0.12 | 0.12 | 0.00 | 0.45 | 15485 | 1.00 |
| | $\boldsymbol{\rho}$ | **13.43** | **5.82** | **2.54** | **25.67** | **6634** | **1.00** | | $\rho$ | 10.49 | 6.55 | 0.87 | 25.44 | 12042 | 1.00 |
| | $\boldsymbol{\epsilon}$ | **0.29** | **0.21** | **0.01** | **0.79** | **7649** | **1.00** | | $\epsilon$ | 0.04 | 0.04 | 0.00 | 0.15 | 12760 | 1.00 |
| | $\boldsymbol{\nu}$ | **8.09** | **4.58** | **1.22** | **18.71** | **7393** | **1.00** | | $\nu$ | 12.05 | 6.32 | 2.19 | 26.07 | 11456 | 1.00 |
| | $\boldsymbol{\theta}$ | **0.92** | **0.52** | **0.13** | **2.11** | **7285** | **1.00** | | $\theta$ | 0.32 | 0.27 | 0.01 | 1.01 | 10238 | 1.00 |
| 7 | $\alpha$ | 1.52 | 0.28 | 1.03 | 2.11 | 1598 | 1.00 | 8 | $\alpha$ | 2.09 | 0.40 | 1.37 | 2.94 | 8917 | 1.00 |
| | $\beta$ | 0.79 | 0.05 | 0.70 | 0.90 | 1940 | 1.00 | | $\beta$ | 1.72 | 0.35 | 1.15 | 2.52 | 9139 | 1.00 |
| | $\lambda$ | 3.52 | 0.58 | 2.49 | 4.75 | 2589 | 1.00 | | $\lambda$ | 2.15 | 0.67 | 0.93 | 3.54 | 12220 | 1.00 |
| | $\kappa$ | 0.09 | 0.09 | 0.00 | 0.32 | 3073 | 1.00 | | $\kappa$ | 0.23 | 0.21 | 0.01 | 0.79 | 16586 | 1.00 |
| | $\rho$ | 19.54 | 17.76 | 1.29 | 65.70 | 351 | 1.00 | | $\rho$ | 18.53 | 4.26 | 11.00 | 28.17 | 7802 | 1.00 |
| | $\epsilon$ | 0.12 | 0.28 | 0.00 | 1.02 | 286 | 1.01 | | $\epsilon$ | 1.23 | 0.57 | 0.29 | 2.51 | 8130 | 1.00 |
| | $\nu$ | 11.43 | 6.14 | 2.13 | 25.27 | 2230 | 1.00 | | $\nu$ | 12.07 | 6.39 | 2.09 | 26.61 | 13105 | 1.00 |
| | $\theta$ | 0.49 | 0.34 | 0.06 | 1.34 | 2863 | 1.00 | | $\theta$ | 0.28 | 0.28 | 0.01 | 1.04 | 11792 | 1.00 |
| 9 | $\alpha$ | 2.41 | 0.34 | 1.76 | 3.11 | 6956 | 1.00 | 10 | $\alpha$ | 1.86 | 0.28 | 1.35 | 2.43 | 7795 | 1.00 |
| | $\beta$ | 1.50 | 0.20 | 1.16 | 1.93 | 7091 | 1.00 | | $\beta$ | 1.40 | 0.17 | 1.10 | 1.76 | 8025 | 1.00 |
| | $\lambda$ | 2.55 | 0.64 | 1.40 | 3.88 | 11024 | 1.00 | | $\lambda$ | 2.66 | 0.61 | 1.57 | 3.94 | 13170 | 1.00 |
| | $\kappa$ | 0.12 | 0.12 | 0.00 | 0.44 | 14817 | 1.00 | | $\kappa$ | 0.10 | 0.11 | 0.00 | 0.39 | 14556 | 1.00 |
| | $\rho$ | 23.04 | 10.77 | 3.08 | 42.05 | 3866 | 1.00 | | $\rho$ | 47.13 | 6.33 | 34.10 | 58.32 | 2757 | 1.00 |
| | $\epsilon$ | 0.26 | 0.24 | 0.00 | 0.80 | 5111 | 1.00 | | $\epsilon$ | 1.39 | 0.53 | 0.47 | 2.54 | 6878 | 1.00 |
| | $\nu$ | 12.32 | 6.36 | 2.09 | 26.97 | 9642 | 1.00 | | $\nu$ | 12.23 | 6.50 | 2.18 | 26.18 | 10850 | 1.00 |
| | $\theta$ | 0.12 | 0.12 | 0.00 | 0.46 | 10317 | 1.00 | | $\theta$ | 0.24 | 0.23 | 0.01 | 0.87 | 11821 | 1.00 |
| 11 | $\alpha$ | 1.94 | 0.16 | 1.66 | 2.27 | 7955 | 1.00 | 12 | $\alpha$ | 2.37 | 0.29 | 1.83 | 2.97 | 7521 | 1.00 |
| | $\beta$ | 1.33 | 0.07 | 1.20 | 1.47 | 8172 | 1.00 | | $\beta$ | 1.41 | 0.14 | 1.16 | 1.70 | 7434 | 1.00 |
| | $\lambda$ | 3.28 | 0.55 | 2.29 | 4.44 | 10974 | 1.00 | | $\lambda$ | 2.92 | 0.59 | 1.84 | 4.18 | 10641 | 1.00 |
| | $\kappa$ | 0.06 | 0.06 | 0.00 | 0.23 | 15316 | 1.00 | | $\kappa$ | 0.09 | 0.09 | 0.00 | 0.33 | 14030 | 1.00 |
| | $\rho$ | 3.84 | 3.79 | 0.08 | 14.00 | 9824 | 1.00 | | $\rho$ | 25.96 | 13.54 | 2.59 | 48.40 | 2760 | 1.00 |
| | $\epsilon$ | 0.06 | 0.03 | 0.01 | 0.14 | 6593 | 1.00 | | $\epsilon$ | 0.27 | 0.26 | 0.00 | 0.87 | 3543 | 1.00 |
| | $\nu$ | 11.84 | 6.28 | 2.09 | 26.10 | 6994 | 1.00 | | $\nu$ | 12.31 | 6.45 | 2.17 | 26.62 | 9261 | 1.00 |
| | $\theta$ | 0.37 | 0.21 | 0.06 | 0.87 | 7170 | 1.00 | | $\theta$ | 0.20 | 0.18 | 0.01 | 0.66 | 8482 | 1.00 |
| 13 | $\alpha$ | 1.98 | 0.25 | 1.53 | 2.51 | 8152 | 1.00 | 14 | $\alpha$ | 1.59 | 0.30 | 1.06 | 2.23 | 7162 | 1.00 |
| | $\beta$ | 1.07 | 0.07 | 0.93 | 1.22 | 8405 | 1.00 | | $\beta$ | 0.99 | 0.10 | 0.80 | 1.20 | 7611 | 1.00 |
| | $\lambda$ | 3.63 | 0.59 | 2.52 | 4.85 | 13102 | 1.00 | | $\lambda$ | 3.08 | 0.58 | 2.01 | 4.29 | 12091 | 1.00 |
| | $\kappa$ | 0.08 | 0.08 | 0.00 | 0.31 | 15503 | 1.00 | | $\kappa$ | 0.10 | 0.10 | 0.00 | 0.36 | 13936 | 1.00 |
| | $\rho$ | 13.35 | 8.50 | 1.27 | 33.51 | 9463 | 1.00 | | $\rho$ | 25.38 | 8.89 | 5.19 | 40.04 | 3019 | 1.00 |
| | $\epsilon$ | 0.03 | 0.05 | 0.00 | 0.18 | 9245 | 1.00 | | $\epsilon$ | 0.40 | 0.27 | 0.01 | 0.96 | 4114 | 1.00 |
| | $\nu$ | 12.11 | 6.30 | 2.16 | 26.16 | 9360 | 1.00 | | $\nu$ | 12.08 | 6.34 | 2.11 | 26.36 | 9044 | 1.00 |
| | $\theta$ | 0.33 | 0.24 | 0.03 | 0.92 | 9520 | 1.00 | | $\theta$ | 0.34 | 0.25 | 0.03 | 0.96 | 9471 | 1.00 |
| 15 | $\alpha$ | 1.57 | 0.14 | 1.31 | 1.85 | 5311 | 1.00 | 16 | $\alpha$ | 2.09 | 0.94 | 0.75 | 4.37 | 8716 | 1.00 |
| | $\beta$ | 1.32 | 0.09 | 1.15 | 1.51 | 5319 | 1.00 | | $\beta$ | 0.83 | 0.16 | 0.57 | 1.20 | 9186 | 1.00 |
| | $\lambda$ | 4.34 | 0.55 | 3.32 | 5.49 | 7206 | 1.00 | | $\lambda$ | 1.81 | 0.68 | 0.63 | 3.27 | 12338 | 1.00 |
| | $\kappa$ | 0.06 | 0.07 | 0.00 | 0.24 | 10028 | 1.00 | | $\kappa$ | 0.25 | 0.24 | 0.01 | 0.88 | 17282 | 1.00 |
| | $\rho$ | 32.71 | 20.56 | 2.05 | 62.54 | 1025 | 1.01 | | $\rho$ | 21.41 | 4.98 | 12.92 | 32.42 | 8974 | 1.00 |
| | $\epsilon$ | 0.38 | 0.41 | 0.00 | 1.23 | 1166 | 1.01 | | $\epsilon$ | 1.09 | 0.47 | 0.31 | 2.14 | 11795 | 1.00 |
| | $\nu$ | 12.41 | 6.50 | 2.16 | 26.87 | 5935 | 1.00 | | $\nu$ | 9.83 | 6.00 | 1.37 | 23.80 | 13237 | 1.00 |
| | $\theta$ | 0.06 | 0.06 | 0.00 | 0.22 | 5988 | 1.00 | | $\theta$ | 0.63 | 0.52 | 0.02 | 1.92 | 12035 | 1.00 |

| ID | Parameter | Mean | SD | 2.5% | 97.5% | n_eff | Rhat | ID | Parameter | Mean | SD | 2.5% | 97.5% | n_eff | Rhat |
|---|---|---|---|---|---|---|---|---|---|---|---|---|---|---|---|
| 17 | $\alpha$ | 1.85 | 0.49 | 0.99 | 2.90 | 10067 | 1.00 | 18 | $\alpha$ | 1.45 | 0.19 | 1.10 | 1.86 | 9641 | 1.00 |
| | $\beta$ | 1.01 | 0.16 | 0.73 | 1.35 | 9916 | 1.00 | | $\beta$ | 1.02 | 0.07 | 0.89 | 1.15 | 9744 | 1.00 |
| | $\lambda$ | 2.07 | 0.66 | 0.92 | 3.46 | 13907 | 1.00 | | $\lambda$ | 3.19 | 0.57 | 2.15 | 4.37 | 14942 | 1.00 |
| | $\kappa$ | 0.19 | 0.19 | 0.01 | 0.71 | 16683 | 1.00 | | $\kappa$ | 0.07 | 0.07 | 0.00 | 0.28 | 16737 | 1.00 |
| | $\rho$ | 31.98 | 4.29 | 24.45 | 41.47 | 14377 | 1.00 | | $\rho$ | 11.00 | 6.83 | 0.90 | 26.42 | 11822 | 1.00 |
| | $\epsilon$ | 1.71 | 0.52 | 0.83 | 2.87 | 17568 | 1.00 | | $\epsilon$ | 0.03 | 0.03 | 0.00 | 0.12 | 13276 | 1.00 |
| | $\nu$ | 11.91 | 6.35 | 2.03 | 26.20 | 11923 | 1.00 | | $\nu$ | 11.94 | 6.30 | 2.04 | 25.99 | 9127 | 1.00 |
| | $\theta$ | 0.33 | 0.32 | 0.01 | 1.17 | 12553 | 1.00 | | $\theta$ | 0.36 | 0.23 | 0.05 | 0.92 | 9524 | 1.00 |
| 19 | $\alpha$ | 11.18 | 6.04 | 1.91 | 25.11 | 9951 | 1.00 | 20 | $\alpha$ | 9.02 | 5.88 | 1.21 | 22.94 | 11626 | 1.00 |
| | $\beta$ | 0.91 | 0.66 | 0.03 | 2.43 | 8012 | 1.00 | | $\beta$ | 1.04 | 0.62 | 0.11 | 2.44 | 10971 | 1.00 |
| | $\lambda$ | 0.88 | 0.62 | 0.05 | 2.33 | 14272 | 1.00 | | $\lambda$ | 0.94 | 0.65 | 0.05 | 2.46 | 12841 | 1.00 |
| | $\kappa$ | 0.70 | 0.57 | 0.03 | 2.10 | 16495 | 1.00 | | $\kappa$ | 0.72 | 0.58 | 0.03 | 2.14 | 13590 | 1.00 |
| | $\rho$ | 6.58 | 5.41 | 0.57 | 20.04 | 12450 | 1.00 | | $\rho$ | 5.27 | 5.07 | 0.40 | 18.94 | 10298 | 1.00 |
| | $\epsilon$ | 0.68 | 0.55 | 0.03 | 2.04 | 11015 | 1.00 | | $\epsilon$ | 0.67 | 0.56 | 0.03 | 2.08 | 10358 | 1.00 |
| | $\nu$ | 8.72 | 5.94 | 0.81 | 22.91 | 14189 | 1.00 | | $\nu$ | 9.04 | 5.90 | 1.10 | 22.94 | 12648 | 1.00 |
| | $\theta$ | 0.74 | 0.58 | 0.02 | 2.13 | 13548 | 1.00 | | $\theta$ | 0.71 | 0.56 | 0.03 | 2.04 | 12502 | 1.00 |
| 21 | $\alpha$ | 1.78 | 1.12 | 0.45 | 4.61 | 9166 | 1.00 | | | | | | | | |
| | $\beta$ | 0.75 | 0.17 | 0.49 | 1.14 | 9372 | 1.00 | | | | | | | | |
| | $\lambda$ | 1.76 | 0.67 | 0.59 | 3.16 | 15167 | 1.00 | | | | | | | | |
| | $\kappa$ | 0.34 | 0.32 | 0.01 | 1.18 | 16825 | 1.00 | | | | | | | | |
| | $\rho$ | 18.31 | 4.90 | 10.75 | 29.70 | 11854 | 1.00 | | | | | | | | |
| | $\epsilon$ | 1.57 | 0.59 | 0.57 | 2.84 | 12237 | 1.00 | | | | | | | | |
| | $\nu$ | 8.89 | 5.78 | 1.05 | 22.71 | 12967 | 1.00 | | | | | | | | |
| | $\theta$ | 0.73 | 0.56 | 0.03 | 2.11 | 12615 | 1.00 | | | | | | | | |

TABLE IV. Parameter estimates for Model IV. These results are obtained in the first-stage algorithm at the individual level. This model was the best model for none of the individuals.

#### WAIC

| | | Model I | | | Model II | | | Model III | | | Model IV | | |
|---|---|---|---|---|---|---|---|---|---|---|---|---|---|
| Source | ID | WAIC | | $P_W$ | WAIC | | $P_W$ | WAIC | | $P_W$ | WAIC | | $P_W$ |
| | | Est | $\delta$ | Est | Est | $\delta$ | Est | Est | $\delta$ | Est | Est | $\delta$ | Est |
| Banff Park | 1 | 891.80 | 137.29 | 1.99 | 778.55 | 24.04 | 3.20 | **754.51** | **0.00** | **3.64** | 781.21 | 26.70 | 4.98 |
| | 2 | 1853.59 | 219.42 | 5.03 | 1639.01 | 4.84 | 5.88 | **1634.17** | **0.00** | **6.07** | 1705.45 | 71.28 | 7.77 |
| | 3 | 2739.68 | 351.31 | 2.84 | 2404.02 | 15.65 | 4.04 | **2388.37** | **0.00** | **4.75** | 2463.30 | 74.93 | 6.85 |
| | 4 | 275.76 | 23.43 | 1.87 | 257.40 | 5.07 | 2.99 | **252.33** | **0.00** | **3.05** | 263.37 | 11.04 | 3.53 |
| | 5 | 578.03 | 77.05 | 2.61 | 501.76 | 0.78 | 3.77 | 501.28 | 0.30 | 3.81 | **500.98** | **0.00** | **4.06** |
| | 6 | 1128.17 | 189.78 | 4.14 | 938.73 | 0.34 | 5.76 | **938.39** | **0.00** | **4.73** | 974.36 | 35.97 | 5.62 |
| Cross Ranch | 7 | 1658.74 | 76.84 | 3.11 | **1581.90** | **0.00** | **4.56** | 1615.96 | 34.06 | 3.94 | 1641.48 | 59.58 | 7.27 |
| | 8 | 388.13 | 22.68 | 1.59 | 371.02 | 5.57 | 2.60 | **365.45** | **0.00** | **2.72** | 372.61 | 7.16 | 3.00 |
| | 9 | 830.40 | 48.43 | 1.89 | **781.97** | **0.00** | **3.40** | 794.71 | 12.74 | 3.28 | 817.23 | 35.26 | 4.46 |
| | 10 | 780.99 | 12.51 | 2.02 | **768.48** | **0.00** | **3.62** | 789.60 | 21.12 | 5.75 | 794.64 | 26.16 | 5.25 |
| | 11 | 3400.63 | 676.57 | 4.61 | **2724.06** | **0.00** | **4.97** | 2731.12 | 7.06 | 5.17 | 2841.78 | 117.72 | 7.77 |
| | 12 | 1248.45 | 103.88 | 2.10 | **1144.57** | **0.00** | **3.82** | 1150.78 | 6.21 | 4.37 | 1192.20 | 47.63 | 5.63 |
| | 13 | 1722.04 | 160.84 | 4.42 | **1561.20** | **0.00** | **6.35** | 1566.64 | 5.44 | 5.68 | 1610.65 | 49.45 | 6.25 |
| | 14 | 1012.85 | 102.93 | 3.02 | 911.82 | 1.90 | 4.42 | **909.92** | **0.00** | **4.90** | 941.88 | 31.96 | 6.80 |
| | 15 | 2539.97 | 212.93 | 2.35 | 2334.23 | 7.19 | 3.48 | **2327.04** | **0.00** | **4.69** | 2395.85 | 68.81 | 8.91 |
| Elk Island National Park | 16 | 273.84 | 6.26 | 2.74 | **267.58** | **0.00** | **3.88** | 269.03 | 1.45 | 3.98 | 272.59 | 5.01 | 4.50 |
| | 17 | 454.51 | 9.23 | 2.56 | 445.67 | 3.39 | 3.83 | **442.28** | **0.00** | **3.81** | 446.28 | 4.00 | 4.40 |
| | 18 | 1599.71 | 121.46 | 3.72 | 1478.28 | 0.03 | 4.19 | **1478.25** | **0.00** | **2.96** | 1528.43 | 50.18 | 4.49 |
| | 19 | 20.96 | 4.64 | 0.11 | 17.35 | 1.03 | 0.88 | **16.32** | **0.00** | **0.54** | 17.58 | 1.26 | 0.57 |
| | 20 | 26.16 | 8.21 | 0.47 | 19.03 | 1.08 | 0.98 | **17.95** | **0.00** | **0.61** | 19.34 | 1.39 | 0.66 |
| | 21 | **151.95** | **0.00** | **3.12** | 154.23 | 2.28 | 3.78 | 153.25 | 1.30 | 3.50 | 154.96 | 3.01 | 3.61 |

TABLE V. WAIC from the pointwise log-likelihood for each model and individual, at the individual level. The point estimate for the information criterion WAIC is denoted as Est, the effective number of parameters as $P_W$, and the difference between the WAIC and that of the best model for a given individual as $\delta$.

#### Posterior Predictive Check (PPC)

Each row of the Figure below displays the PPCs (here, the number of unique visited sites as a function of time) of all models, for a same individual. Thus, each row allows us to compare the different models for animal $i$, with $i = 1 : 21$. The blue symbols show the observed number of unique visited sites, whereas the light grey symbols represent the PPCs generated by 1000 independent samples, and the dark grey symbols the 95% confidence intervals (CIs).

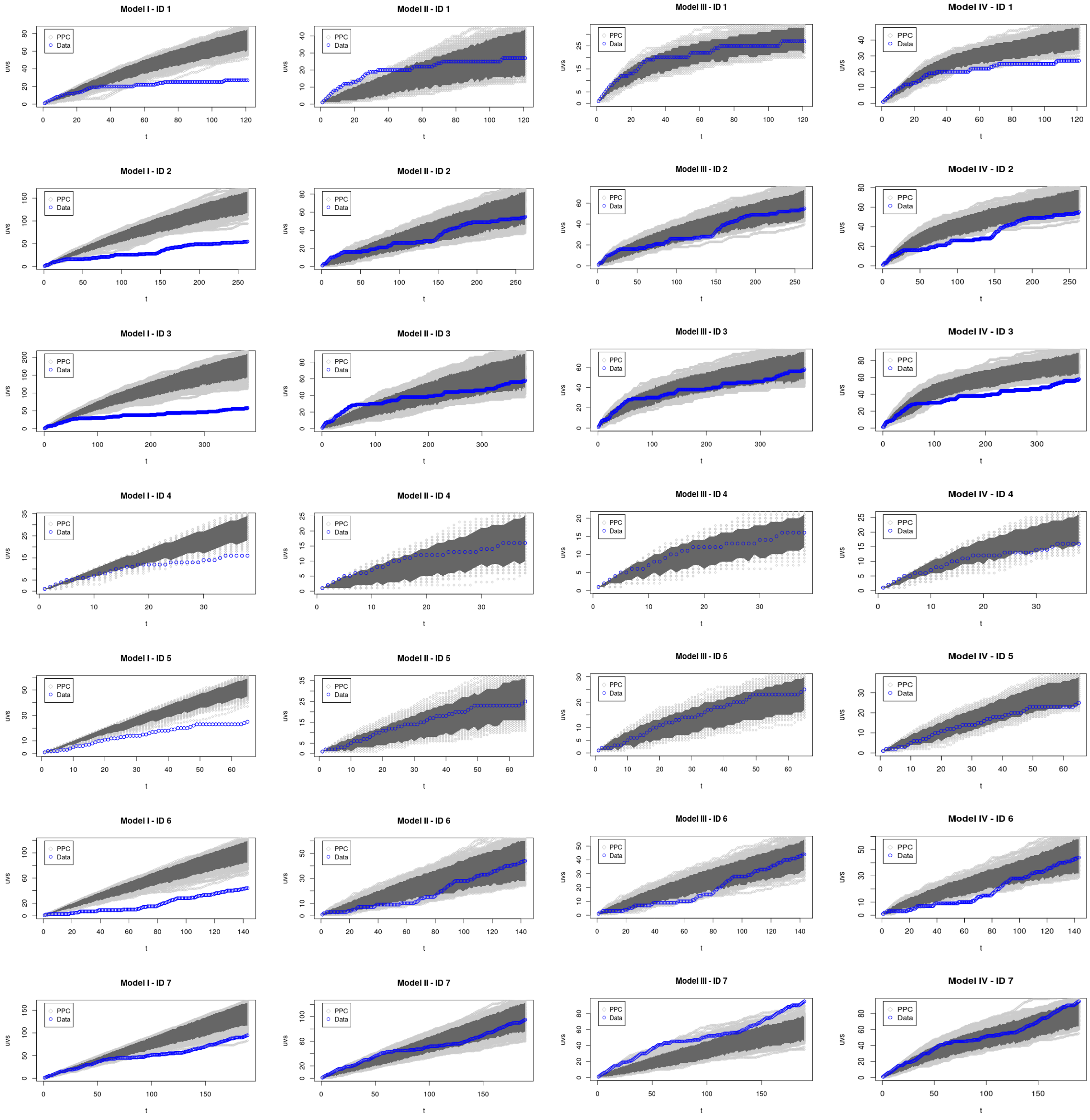

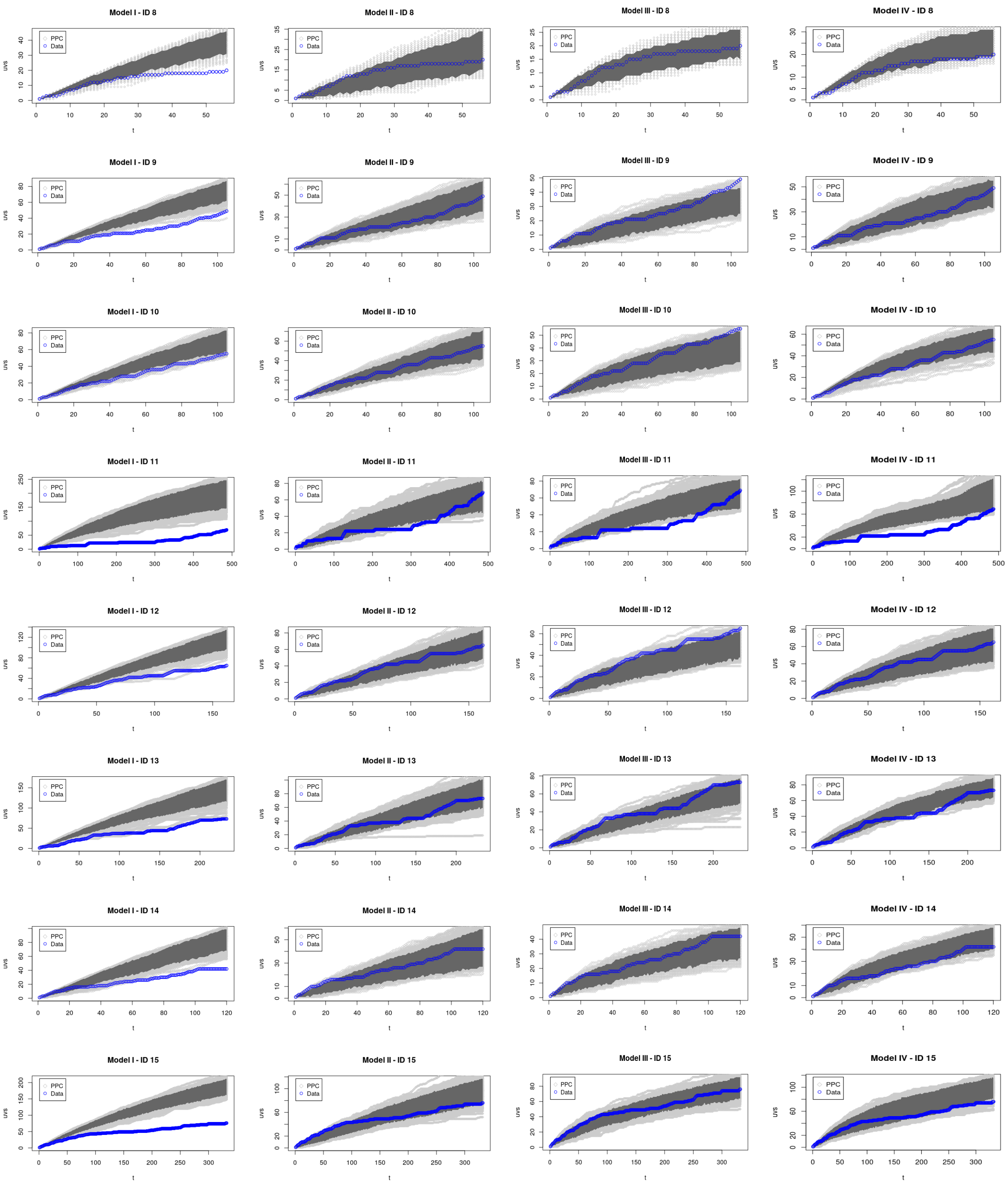
Model I - ID 8
Model II - ID 8
Model III - ID 8
Model IV - ID 8
Model I - ID 9
Model II - ID 9
Model III - ID 9
Model IV - ID 9
Model I - ID 10
Model II - ID 10
Model III - ID 10
Model IV - ID 10
Model I - ID 11
Model II - ID 11
Model III - ID 11
Model IV - ID 11
Model I - ID 12
Model II - ID 12
Model III - ID 12
Model IV - ID 12
Model I - ID 13
Model II - ID 13
Model III - ID 13
Model IV - ID 13
Model I - ID 14
Model II - ID 14
Model III - ID 14
Model IV - ID 14
Model I - ID 15
Model II - ID 15
Model III - ID 15
Model IV - ID 15
PPC
Data
uvs
t

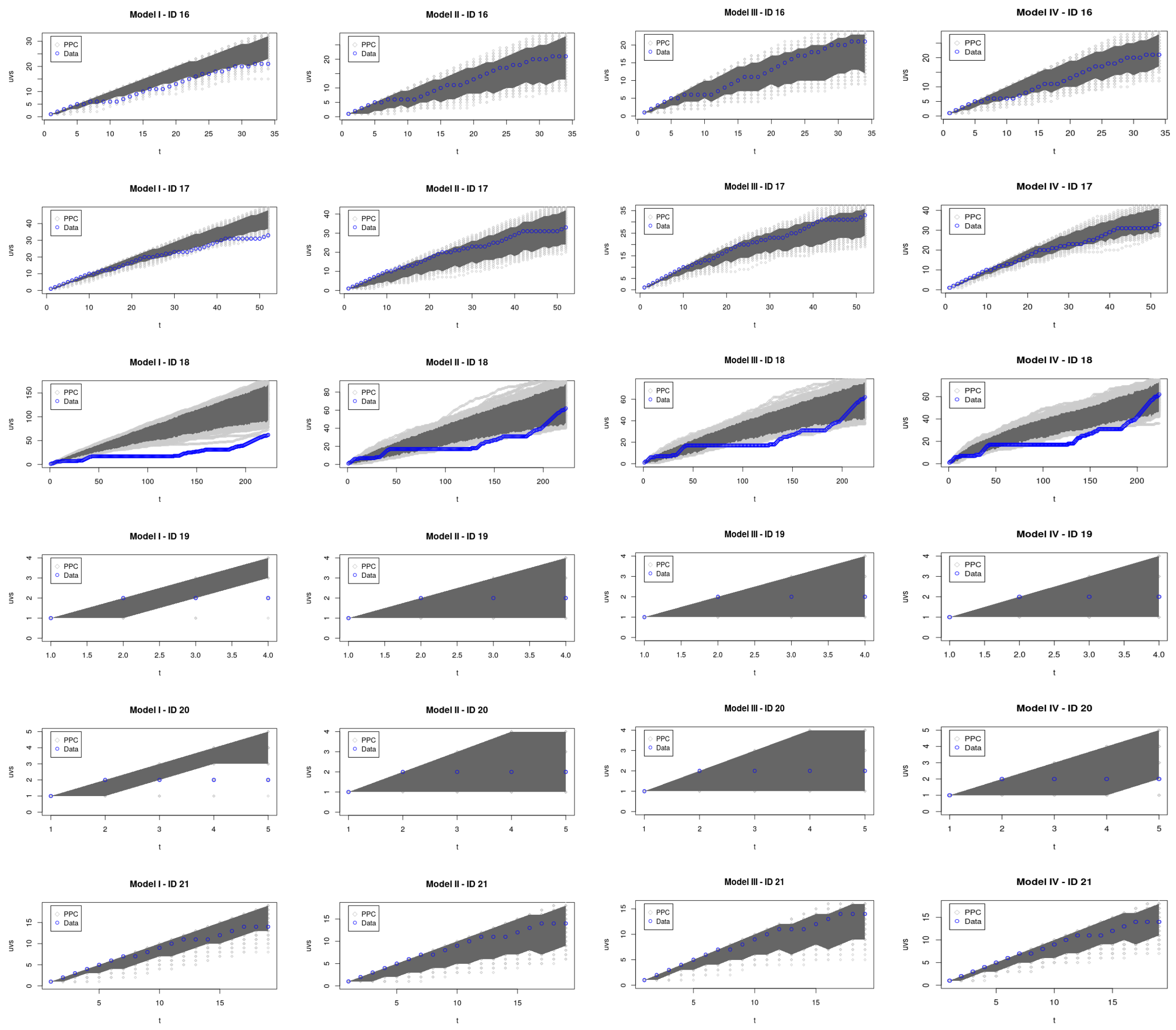

FIG. 1. Posterior predictive check (PPC) of Models I-IV (columns) for each observed animal (rows), at the individual level. The PPC is the number of unique visited sites (UVS) as a function of time $t$. The curves of the PPC are shown in light grey, the CIs at 95% in dark grey, and the data from the real trajectories in blue. Note that the $y-$scale for each graph in a same row may vary in order to better display which parts of the real trajectory are inside of the CI.

#### HIERARCHICAL LEVEL

##### Model I

| Parameter | $\alpha$ | | | | | | $\beta$ | | | | | |
|---|---|---|---|---|---|---|---|---|---|---|---|---|
| ID | Mean | SD | 2.5% | 97.5% | $p_v$ | n_eff | Mean | SD | 2.5% | 97.5% | $p_v$ | n_eff |
| 1 | 1.29 | 0.18 | 1.17 | 1.66 | 0.16 | 3048 | 1.11 | 0.11 | 1.03 | 1.35 | 0.68 | 5676 |
| 2 | 0.71 | 0.07 | 0.66 | 0.87 | 0.94 | 4842 | 0.86 | 0.04 | 0.83 | 0.95 | 0.70 | 7500 |
| 3 | 1.12 | 0.08 | 1.07 | 1.28 | 0.46 | 6426 | 1.27 | 0.06 | 1.22 | 1.40 | 0.83 | 5360 |
| 4 | 1.13 | 0.25 | 0.95 | 1.68 | 0.87 | 4066 | 0.96 | 0.16 | 0.85 | 1.31 | 0.78 | 6445 |
| 5 | 1.06 | 0.25 | 0.88 | 1.58 | 0.53 | 4783 | 0.72 | 0.09 | 0.66 | 0.90 | 0.98 | 5690 |
| 6 | 1.21 | 0.18 | 1.08 | 1.60 | 0.90 | 4082 | 0.86 | 0.07 | 0.81 | 1.00 | 0.34 | 7204 |
| 7 | 0.60 | 0.09 | 0.53 | 0.81 | 0.31 | 3318 | 0.64 | 0.03 | 0.62 | 0.71 | 0.39 | 6281 |
| 8 | 1.27 | 0.19 | 1.13 | 1.68 | 0.96 | 3392 | 1.30 | 0.17 | 1.18 | 1.67 | 0.21 | 3628 |
| 9 | 1.20 | 0.18 | 1.06 | 1.58 | 0.34 | 3951 | 1.05 | 0.11 | 0.97 | 1.28 | 0.91 | 6272 |
| 10 | 1.28 | 0.18 | 1.15 | 1.65 | 0.18 | 3734 | 1.16 | 0.11 | 1.08 | 1.41 | 0.71 | 5181 |
| 11 | 0.82 | 0.05 | 0.74 | 0.92 | 0.96 | 5498 | 1.02 | 0.03 | 0.99 | 1.10 | 0.72 | 6679 |
| 12 | 1.14 | 0.15 | 1.03 | 1.45 | 0.63 | 5856 | 1.00 | 0.07 | 0.94 | 1.17 | 0.20 | 7087 |
| 13 | 0.97 | 0.11 | 0.89 | 1.20 | 0.80 | 6002 | 0.87 | 0.05 | 0.84 | 0.97 | 0.52 | 6756 |
| 14 | 0.79 | 0.15 | 0.68 | 1.11 | 0.73 | 4759 | 0.80 | 0.07 | 0.75 | 0.96 | 0.22 | 6534 |
| 15 | 1.16 | 0.11 | 1.08 | 1.41 | 0.86 | 5749 | 1.13 | 0.07 | 1.07 | 1.30 | 0.47 | 6270 |
| 16 | 0.88 | 0.26 | 0.69 | 1.46 | 0.97 | 4170 | 0.68 | 0.09 | 0.61 | 0.89 | 0.67 | 5422 |
| 17 | 1.10 | 0.25 | 0.92 | 1.62 | 0.56 | 4244 | 0.87 | 0.12 | 0.78 | 1.15 | 0.37 | 6756 |
| 18 | 0.66 | 0.07 | 0.61 | 0.83 | 0.90 | 4616 | 0.83 | 0.04 | 0.80 | 0.92 | 0.82 | 7165 |
| 19 | 1.03 | 0.38 | 0.78 | 1.64 | 0.42 | 266 | 0.76 | 0.39 | 0.48 | 1.58 | 0.24 | 3601 |
| 20 | 1.04 | 0.29 | 0.82 | 1.68 | 0.80 | 1375 | 0.91 | 0.33 | 0.65 | 1.65 | 0.80 | 3931 |
| 21 | 0.93 | 0.30 | 0.71 | 1.57 | 0.79 | 3694 | 0.69 | 0.13 | 0.59 | 1.00 | 0.27 | 4494 |
| pop | 1.02 | 0.09 | 0.96 | 1.20 | 0.78 | 2350 | 0.93 | 0.09 | 0.86 | 1.12 | 0.90 | 5142 |

| Parameter | $\lambda$ | | | | | | $\kappa$ | | | | | |
|---|---|---|---|---|---|---|---|---|---|---|---|---|
| ID | Mean | SD | 2.5% | 97.5% | $p_v$ | n_eff | Mean | SD | 2.5% | 97.5% | $p_v$ | n_eff |
| 1 | 3.60 | 0.26 | 3.40 | 4.17 | 0.51 | 757 | 0.07 | 0.07 | 0.02 | 0.26 | 0.09 | 7055 |
| 2 | 3.67 | 0.28 | 3.47 | 4.25 | 0.44 | 535 | 0.04 | 0.04 | 0.01 | 0.16 | 0.77 | 6995 |
| 3 | 3.70 | 0.26 | 3.53 | 4.23 | 0.59 | 586 | 0.03 | 0.03 | 0.00 | 0.12 | 0.21 | 7500 |
| 4 | 3.25 | 0.32 | 3.04 | 3.91 | 0.08 | 1013 | 0.15 | 0.14 | 0.04 | 0.50 | 0.36 | 6262 |
| 5 | 3.30 | 0.29 | 3.10 | 3.92 | 0.20 | 1416 | 0.10 | 0.10 | 0.03 | 0.38 | 0.02 | 6989 |
| 6 | 3.45 | 0.28 | 3.27 | 4.02 | 0.67 | 1020 | 0.06 | 0.06 | 0.01 | 0.22 | 0.26 | 7500 |
| 7 | 3.50 | 0.28 | 3.30 | 4.08 | 0.31 | 1039 | 0.05 | 0.05 | 0.01 | 0.21 | 0.54 | 7500 |
| 8 | 3.24 | 0.29 | 3.05 | 3.79 | 0.40 | 1001 | 0.14 | 0.13 | 0.04 | 0.50 | 0.91 | 6961 |
| 9 | 3.31 | 0.29 | 3.12 | 3.87 | 0.08 | 1566 | 0.07 | 0.07 | 0.02 | 0.25 | 0.23 | 6857 |
| 10 | 3.29 | 0.28 | 3.09 | 3.84 | 0.71 | 1144 | 0.07 | 0.07 | 0.02 | 0.29 | 0.22 | 7072 |
| 11 | 3.61 | 0.25 | 3.44 | 4.12 | 0.59 | 1019 | 0.02 | 0.02 | 0.00 | 0.09 | 0.38 | 7212 |
| 12 | 3.38 | 0.28 | 3.20 | 3.96 | 0.63 | 1353 | 0.05 | 0.05 | 0.01 | 0.18 | 0.06 | 7109 |
| 13 | 3.62 | 0.27 | 3.43 | 4.19 | 0.21 | 615 | 0.04 | 0.04 | 0.01 | 0.16 | 0.49 | 6636 |
| 14 | 3.41 | 0.27 | 3.23 | 3.96 | 0.61 | 1211 | 0.05 | 0.05 | 0.01 | 0.22 | 0.54 | 7306 |
| 15 | 3.81 | 0.27 | 3.60 | 3.45 | 0.59 | 141 | 0.02 | 0.02 | 0.00 | 0.10 | 0.45 | 7212 |
| 16 | 3.19 | 0.33 | 2.98 | 3.80 | 0.72 | 358 | 0.17 | 0.15 | 0.06 | 0.56 | 0.63 | 6598 |
| 17 | 3.22 | 0.30 | 3.02 | 3.82 | 0.89 | 570 | 0.13 | 0.12 | 0.04 | 0.45 | 0.59 | 6543 |
| 18 | 3.46 | 0.27 | 3.27 | 4.01 | 0.12 | 1336 | 0.04 | 0.04 | 0.01 | 0.15 | 0.51 | 7160 |
| 19 | 3.02 | 0.38 | 2.74 | 3.82 | 0.71 | 52 | 0.37 | 0.30 | 0.13 | 1.16 | 0.14 | 3949 |
| 20 | 3.10 | 0.32 | 2.87 | 3.73 | 0.33 | 90 | 0.36 | 0.29 | 0.14 | 1.09 | 0.08 | 4429 |
| 21 | 3.19 | 0.30 | 2.98 | 3.77 | 0.85 | 262 | 0.24 | 0.21 | 0.08 | 0.79 | 0.39 | 6013 |
| pop | 3.41 | 0.13 | 3.31 | 3.68 | 0.85 | 207 | 0.11 | 0.14 | 0.03 | 0.38 | 0.90 | 7134 |

TABLE VI. Parameter estimates for Model I. These results are obtained from the second-stage algorithm at the hierarchical level.

#### Model II

| Parameter | $\alpha$ | | | | | | $\beta$ | | | | | | $\lambda$ | | | | | |
|---|---|---|---|---|---|---|---|---|---|---|---|---|---|---|---|---|---|---|
| ID | Mean | SD | 2.5% | 97.5% | $p_v$ | n_eff | Mean | SD | 2.5% | 97.5% | $p_v$ | n_eff | Mean | SD | 2.5% | 97.5% | $p_v$ | n_eff |
| 1 | 2.22 | 0.22 | 2.07 | 2.67 | 0.52 | 1451 | 1.37 | 0.16 | 1.26 | 1.71 | 0.90 | 4399 | 2.77 | 0.29 | 2.57 | 3.36 | 0.07 | 1116 |
| 2 | 1.60 | 0.15 | 1.49 | 1.93 | 0.81 | 1983 | 1.01 | 0.06 | 0.97 | 1.14 | 0.11 | 6598 | 2.81 | 0.29 | 2.60 | 3.42 | 0.27 | 1251 |
| 3 | 1.72 | 0.11 | 1.64 | 1.96 | 0.80 | 4534 | 1.47 | 0.09 | 1.41 | 1.66 | 0.56 | 4588 | 2.87 | 0.29 | 2.66 | 3.45 | 0.68 | 1282 |
| 4 | 2.02 | 0.28 | 1.82 | 2.60 | 0.96 | 1770 | 1.21 | 0.21 | 1.05 | 1.67 | 0.09 | 5931 | 2.58 | 0.31 | 2.37 | 3.21 | 0.13 | 1083 |
| 5 | 2.16 | 0.29 | 1.99 | 2.78 | 0.81 | 277 | 0.98 | 0.14 | 0.88 | 1.28 | 0.77 | 5561 | 2.59 | 0.31 | 2.38 | 3.20 | 0.55 | 1237 |
| 6 | 2.49 | 0.22 | 2.32 | 2.98 | 0.91 | 108 | 1.15 | 0.10 | 1.08 | 1.36 | 0.31 | 6539 | 2.70 | 0.29 | 2.50 | 3.29 | 0.50 | 869 |
| **7** | **1.53** | **0.22** | **1.37** | **1.98** | **0.27** | **944** | **0.77** | **0.04** | **0.73** | **0.86** | **0.90** | **5886** | **2.91** | **0.28** | **2.73** | **3.46** | **0.28** | **853** |
| 8 | 2.00 | 0.24 | 1.83 | 2.49 | 0.23 | 3371 | 1.48 | 0.21 | 1.33 | 1.93 | 0.10 | 2445 | 2.60 | 0.33 | 2.38 | 3.26 | 0.93 | 734 |
| **9** | **2.09** | **0.22** | **1.94** | **2.54** | **0.20** | **1907** | **1.37** | **0.15** | **1.26** | **1.71** | **0.08** | **4211** | **2.66** | **0.31** | **2.45** | **3.27** | **0.96** | **1561** |
| **10** | **1.85** | **0.20** | **1.72** | **2.26** | **0.85** | **3856** | **1.33** | **0.13** | **1.24** | **1.62** | **0.09** | **4952** | **2.66** | **0.29** | **2.44** | **3.29** | **0.09** | **1286** |
| **11** | **2.03** | **0.13** | **1.94** | **2.31** | **0.57** | **5034** | **1.29** | **0.06** | **1.25** | **1.43** | **0.23** | **6236** | **2.74** | **0.30** | **2.53** | **3.33** | **0.97** | **1529** |
| **12** | **2.18** | **0.20** | **2.05** | **2.60** | **0.21** | **2099** | **1.37** | **0.12** | **1.28** | **1.63** | **0.36** | **5139** | **2.73** | **0.30** | **2.52** | **3.33** | **0.82** | **1193** |
| **13** | **1.88** | **0.17** | **1.76** | **2.24** | **0.06** | **4223** | **1.06** | **0.06** | **1.02** | **1.20** | **0.55** | **6659** | **2.86** | **0.29** | **2.65** | **3.48** | **0.78** | **1061** |
| **14** | **1.74** | **0.22** | **1.59** | **2.18** | **0.25** | **2752** | **0.99** | **0.10** | **0.92** | **1.20** | **0.45** | **5874** | **2.73** | **0.28** | **2.53** | **3.31** | **0.64** | **1424** |
| **15** | **1.66** | **0.12** | **1.57** | **1.91** | **0.67** | **3490** | **1.29** | **0.08** | **1.23** | **1.46** | **0.60** | **6365** | **3.08** | **0.33** | **2.85** | **3.66** | **0.09** | **308** |
| **16** | **1.89** | **0.29** | **1.70** | **245** | **0.55** | **2224** | **0.87** | **0.14** | **0.77** | **1.20** | **0.19** | **4027** | **2.55** | **0.32** | **2.31** | **3.21** | **0.18** | **858** |
| **17** | **1.85** | **0.27** | **1.66** | **2.38** | **0.84** | **2997** | **1.01** | **0.14** | **0.90** | **1.32** | **0.74** | **5804** | **2.58** | **0.29** | **2.37** | **3.20** | **0.96** | **1153** |
| **18** | **1.57** | **0.17** | **1.44** | **1.93** | **0.17** | **1777** | **0.99** | **0.06** | **0.94** | **1.13** | **0.20** | **6378** | **2.78** | **0.33** | **2.59** | **3.39** | **0.20** | **1358** |
| 19 | 1.91 | 0.37 | 1.77 | 2.60 | 0.87 | 110 | 1.03 | 0.35 | 0.79 | 1.71 | 0.62 | 3009 | 2.42 | 0.36 | 2.15 | 3.15 | 0.17 | 179 |
| 20 | 1.90 | 0.33 | 1.71 | 2.52 | 0.97 | 368 | 1.06 | 0.34 | 0.82 | 1.76 | 0.19 | 3978 | 2.42 | 0.33 | 2.21 | 3.04 | 0.32 | 165 |
| **21** | **1.81** | **0.32** | **1.59** | **2.45** | **0.60** | **1080** | **0.78** | **0.15** | **0.66** | **1.12** | **0.41** | **3224** | **2.55** | **0.32** | **2.33** | **3.19** | **0.30** | **801** |
| pop | 1.91 | 0.10 | 1.85 | 2.11 | 0.35 | 520 | 1.14 | 0.09 | 1.08 | 1.31 | 0.90 | 4312 | 2.70 | 0.14 | 2.60 | 2.99 | 0.34 | 349 |

| Parameter | $\kappa$ | | | | | | $q$ | | | | | |
|---|---|---|---|---|---|---|---|---|---|---|---|---|
| ID | Mean | SD | 2.5% | 97.5% | $p_v$ | n_eff | Mean | SD | 2.5% | 97.5% | $p_v$ | n_eff |
| 1 | 0.14 | 0.13 | 0.04 | 0.49 | 0.83 | 7221 | 0.67 | 0.05 | 0.63 | 0.77 | 0.38 | 7108 |
| 2 | 0.09 | 0.09 | 0.02 | 0.33 | 0.38 | 7500 | 0.67 | 0.03 | 0.65 | 0.74 | 0.34 | 7253 |
| 3 | 0.08 | 0.08 | 0.02 | 0.32 | 0.10 | 6640 | 0.70 | 0.03 | 0.67 | 0.76 | 0.39 | 7452 |
| 4 | 0.21 | 0.18 | 0.07 | 0.69 | 0.41 | 6696 | 0.48 | 0.09 | 0.41 | 0.66 | 0.59 | 7500 |
| 5 | 0.17 | 0.15 | 0.05 | 0.59 | 0.43 | 5743 | 0.56 | 0.06 | 0.51 | 0.69 | 0.36 | 7383 |
| 6 | 0.12 | 0.11 | 0.03 | 0.43 | 0.07 | 6878 | 0.62 | 0.04 | 0.59 | 0.71 | 0.03 | 7176 |
| **7** | **0.08** | **0.07** | **0.02** | **0.29** | **0.90** | **6953** | **0.38** | **0.04** | **0.35** | **0.47** | **0.36** | **7060** |
| 8 | 0.19 | 0.17 | 0.06 | 0.65 | 0.25 | 6869 | 0.48 | 0.09 | 0.42 | 0.66 | 0.96 | 7096 |
| **9** | **0.10** | **0.10** | **0.03** | **0.36** | **0.99** | **7020** | **0.42** | **0.05** | **0.38** | **0.53** | **0.30** | **6957** |
| **10** | **0.09** | **0.09** | **0.03** | **0.34** | **0.16** | **6622** | **0.27** | **0.06** | **0.22** | **0.40** | **0.27** | **6473** |
| **11** | **0.08** | **0.08** | **0.02** | **0.30** | **0.83** | **7322** | **0.80** | **0.02** | **0.78** | **0.84** | **0.21** | **7243** |
| **12** | **0.08** | **0.08** | **0.02** | **0.31** | **0.81** | **6990** | **0.49** | **0.04** | **0.45** | **0.58** | **0.99** | **7826** |
| **13** | **0.08** | **0.08** | **0.02** | **0.30** | **0.30** | **6955** | **0.54** | **0.04** | **0.51** | **0.62** | **0.25** | **7500** |
| **14** | **0.10** | **0.09** | **0.03** | **0.37** | **0.37** | **6611** | **0.54** | **0.05** | **0.50** | **0.64** | **0.24** | **7500** |
| **15** | **0.06** | **0.06** | **0.01** | **0.22** | **0.62** | **7002** | **0.57** | **0.03** | **0.54** | **0.64** | **0.90** | **7113** |
| **16** | **0.21** | **0.19** | **0.07** | **0.69** | **0.90** | **6318** | **0.32** | **0.09** | **0.25** | **0.50** | **0.69** | **6157** |
| **17** | **0.16** | **0.15** | **0.05** | **0.57** | **0.46** | **6269** | **0.25** | **0.07** | **0.20** | **0.40** | **0.72** | **6127** |
| **18** | **0.07** | **0.07** | **0.02** | **0.28** | **0.71** | **6704** | **0.60** | **0.04** | **0.57** | **0.68** | **0.31** | **7488** |
| 19 | 0.40 | 0.30 | 0.17 | 1.13 | 0.96 | 3849 | 0.55 | 0.20 | 0.40 | 0.91 | 0.55 | 6651 |
| 20 | 0.42 | 0.31 | 0.17 | 1.15 | 0.12 | 3845 | 0.61 | 0.18 | 0.49 | 0.93 | 0.00 | 6510 |
| **21** | **0.26** | **0.21** | **0.09** | **0.79** | **0.73** | **6004** | **0.18** | **0.11** | **0.09** | **0.42** | **0.51** | **5226** |
| pop | 0.15 | 0.12 | 0.08 | 0.40 | 0.66 | 6487 | 0.51 | 0.11 | 0.44 | 0.73 | 0.86 | 7214 |

TABLE VII. Parameter estimates for Model II. These results are obtained from the second-stage algorithm at the hierarchical level. The individuals for which this model was the best one, according to the comparison of Table II in the main paper, are highlighted in bold.

#### Model III

| Parameter | $\alpha$ | | | | | | $\beta$ | | | | | | $\lambda$ | | | | | |
|---|---|---|---|---|---|---|---|---|---|---|---|---|---|---|---|---|---|---|
| ID | Mean | SD | 2.5% | 97.5% | $p_v$ | n_eff | Mean | SD | 2.5% | 97.5% | $p_v$ | n_eff | Mean | SD | 2.5% | 97.5% | $p_v$ | n_eff |
| **1** | **2.29** | **0.22** | **2.14** | **2.74** | **0.32** | **2496** | **1.39** | **0.16** | **1.27** | **1.73** | **0.29** | **8290** | **2.66** | **0.29** | **2.46** | **3.37** | **0.18** | **2054** |
| **2** | **1.64** | **0.16** | **1.53** | **1.98** | **0.76** | **3568** | **1.02** | **0.06** | **0.97** | **1.14** | **0.76** | **12880** | **2.73** | **0.29** | **2.52** | **3.31** | **0.67** | **2237** |
| **3** | **1.75** | **0.11** | **1.67** | **1.99** | **0.54** | **7123** | **1.48** | **0.09** | **1.41** | **1.66** | **0.51** | **9412** | **2.77** | **0.28** | **2.58** | **3.36** | **0.52** | **1765** |
| **4** | **2.09** | **0.28** | **1.90** | **2.66** | **0.68** | **3673** | **1.22** | **0.21** | **1.07** | **1.69** | **0.36** | **10532** | **2.50** | **0.32** | **2.29** | **3.13** | **0.95** | **1800** |
| **5** | **2.21** | **0.29** | **2.00** | **2.78** | **0.98** | **604** | **0.97** | **0.13** | **0.88** | **1.27** | **0.45** | **10620** | **2.50** | **0.32** | **2.27** | **3.14** | **0.59** | **1751** |
| **6** | **2.52** | **0.25** | **2.32** | **3.02** | **0.66** | **186** | **1.15** | **0.10** | **1.09** | **1.36** | **0.48** | **13497** | **2.62** | **0.30** | **2.42** | **3.23** | **0.36** | **2676** |
| 7 | 1.70 | 0.21 | 1.55 | 2.13 | 0.17 | 3921 | 0.79 | 0.05 | 0.76 | 0.89 | 0.19 | 11062 | 2.80 | 0.30 | 2.59 | 3.40 | 0.44 | 1322 |
| **8** | **2.08** | **0.25** | **1.90** | **2.59** | **0.93** | **5184** | **1.48** | **0.22** | **1.32** | **1.94** | **0.55** | **4586** | **2.49** | **0.32** | **2.28** | **3.09** | **0.06** | **1542** |
| 9 | 2.21 | 0.22 | 2.05 | 2.65 | 0.87 | 4084 | 1.42 | 0.15 | 1.31 | 1.76 | 0.45 | 8146 | 2.56 | 0.31 | 2.34 | 3.19 | 0.23 | 1815 |
| 10 | 1.97 | 0.20 | 1.83 | 2.38 | 0.32 | 7109 | 1.38 | 0.14 | 1.29 | 1.68 | 0.54 | 8687 | 2.57 | 0.31 | 2.35 | 3.18 | 0.31 | 2515 |
| 11 | 2.02 | 0.13 | 1.93 | 2.31 | 0.42 | 10623 | 1.29 | 0.06 | 1.25 | 1.43 | 0.09 | 12949 | 2.67 | 0.29 | 2.47 | 3.27 | 0.08 | 2416 |
| 12 | 2.28 | 0.21 | 2.14 | 2.72 | 0.59 | 3084 | 1.40 | 0.12 | 1.31 | 1.67 | 0.90 | 9531 | 2.63 | 0.30 | 2.43 | 3.25 | 0.17 | 1748 |
| 13 | 1.98 | 0.17 | 1.86 | 2.33 | 0.46 | 9540 | 1.08 | 0.06 | 1.03 | 1.23 | 0.31 | 14127 | 2.75 | 0.29 | 2.54 | 3.35 | 0.51 | 1364 |
| 14 | 1.81 | 0.23 | 1.65 | 2.27 | 0.73 | 4991 | 0.99 | 0.10 | 0.92 | 1.20 | 0.77 | 12154 | 2.64 | 0.29 | 2.45 | 3.24 | 0.08 | 2468 |
| 15 | 1.72 | 0.13 | 1.62 | 1.98 | 0.51 | 6574 | 1.30 | 0.09 | 1.24 | 1.49 | 0.40 | 12096 | 2.96 | 0.27 | 2.78 | 3.52 | 0.57 | 600 |
| 16 | 1.96 | 0.30 | 1.75 | 2.56 | 0.26 | 3717 | 0.89 | 0.15 | 0.78 | 1.22 | 0.45 | 8636 | 2.45 | 0.33 | 2.23 | 3.10 | 0.48 | 1586 |
| 17 | 1.93 | 0.27 | 1.74 | 2.47 | 0.65 | 5341 | 1.03 | 0.14 | 0.92 | 1.34 | 0.60 | 11650 | 2.49 | 0.33 | 2.26 | 3.14 | 0.92 | 1699 |
| 18 | 1.62 | 0.18 | 1.49 | 1.99 | 0.37 | 2869 | 0.99 | 0.06 | 0.95 | 1.14 | 0.89 | 12533 | 2.70 | 0.29 | 2.50 | 3.31 | 0.12 | 3068 |
| **19** | **2.04** | **0.39** | **1.81** | **2.74** | **0.35** | **237** | **1.05** | **0.35** | **0.82** | **1.75** | **0.77** | **5696** | **2.33** | **0.37** | **2.06** | **3.12** | **0.15** | **319** |
| **20** | **1.99** | **0.31** | **1.78** | **2.60** | **0.10** | **667** | **1.07** | **0.33** | **0.83** | **1.72** | **0.50** | **7290** | **2.35** | **0.33** | **2.14** | **2.96** | **0.30** | **418** |
| 21 | 1.90 | 0.31 | 1.69 | 2.53 | 0.33 | 3102 | 0.82 | 0.16 | 0.70 | 1.20 | 0.42 | 6068 | 2.44 | 0.32 | 2.23 | 3.07 | 0.30 | 1713 |
| pop | 1.99 | 0.10 | 1.92 | 2.19 | 0.70 | 924 | 1.15 | 0.09 | 1.09 | 1.33 | 0.55 | 8805 | 2.60 | 0.15 | 2.49 | 2.91 | 0.24 | 435 |

| Parameter | $\kappa$ | | | | | | $\rho$ | | | | | | $\epsilon$ | | | | | |
|---|---|---|---|---|---|---|---|---|---|---|---|---|---|---|---|---|---|---|
| ID | Mean | SD | 2.5% | 97.5% | $p_v$ | n_eff | Mean | SD | 2.5% | 97.5% | $p_v$ | n_eff | Mean | SD | 2.5% | 97.5% | $p_v$ | n_eff |
| **1** | **0.15** | **0.14** | **0.05** | **0.51** | **0.48** | **13910** | **12.89** | **0.39** | **12.71** | **13.96** | **0.26** | **21** | **0.86** | **0.23** | **0.70** | **1.34** | **0.48** | **779** |
| **2** | **0.09** | **0.09** | **0.02** | **0.33** | **0.85** | **13908** | **12.88** | **0.40** | **12.68** | **13.92** | **0.33** | **30** | **0.30** | **0.13** | **0.20** | **0.58** | **0.48** | **10975** |
| **3** | **0.09** | **0.09** | **0.02** | **0.33** | **0.62** | **14113** | **12.89** | **0.41** | **12.71** | **13.86** | **0.45** | **24** | **0.52** | **0.15** | **0.41** | **0.84** | **0.24** | **10185** |
| **4** | **0.22** | **0.19** | **0.07** | **0.72** | **013** | **13064** | **12.88** | **0.40** | **12.72** | **14.01** | **0.25** | **33** | **0.60** | **0.24** | **0.43** | **1.10** | **0.91** | **3404** |
| **5** | **0.17** | **0.15** | **0.05** | **0.57** | **0.88** | **13626** | **12.88** | **0.40** | **12.69** | **13.81** | **0.30** | **33** | **0.31** | **0.18** | **0.18** | **0.71** | **0.94** | **10585** |
| **6** | **0.12** | **0.11** | **0.03** | **0.42** | **0.45** | **14237** | **12.88** | **0.42** | **12.71** | **13.89** | **0.37** | **32** | **0.04** | **0.04** | **0.01** | **0.15** | **0.77** | **8557** |
| 7 | 0.08 | 0.08 | 0.02 | 0.31 | 0.03 | 14071 | 12.89 | 0.41 | 12.70 | 14.01 | 0.27 | 33 | 0.08 | 0.14 | 0.00 | 0.54 | 0.52 | 4682 |
| **8** | **0.20** | **0.18** | **0.06** | **0.66** | **0.58** | **12997** | **12.89** | **0.38** | **12.71** | **13.94** | **0.25** | **33** | **0.59** | **0.24** | **0.42** | **1.10** | **0.50** | **3586** |
| 9 | 0.12 | 0.11 | 0.03 | 0.42 | 0.51 | 13636 | 12.90 | 0.43 | 12.73 | 13.99 | 0.23 | 35 | 0.19 | 0.16 | 0.05 | 0.58 | 0.10 | 7399 |
| 10 | 0.11 | 0.10 | 0.03 | 0.39 | 0.07 | 12660 | 12.93 | 0.86 | 12.77 | 13.74 | 0.05 | 290 | 0.53 | 0.25 | 0.37 | 1.03 | 0.34 | 6288 |
| 11 | 0.07 | 0.07 | 0.02 | 0.29 | 0.51 | 13731 | 7.32 | 0.70 | 7.65 | 7.65 | 0.01 | 11 | 0.11 | 0.03 | 0.09 | 0.20 | 0.37 | 10265 |
| 12 | 0.09 | 0.09 | 0.02 | 0.34 | 0.98 | 13828 | 12.90 | 0.40 | 12.76 | 14.11 | 0.30 | 23 | 0.28 | 0.19 | 0.11 | 0.68 | 0.11 | 9362 |
| 13 | 0.08 | 0.08 | 0.02 | 0.32 | 0.55 | 13796 | 12.88 | 0.41 | 12.72 | 15.00 | 0.27 | 32 | 0.06 | 0.07 | 0.01 | 0.25 | 0.76 | 6503 |
| 14 | 0.10 | 0.10 | 0.03 | 0.38 | 0.72 | 13995 | 12.90 | 0.42 | 12.70 | 14.11 | 0.26 | 27 | 0.42 | 0.19 | 0.28 | 0.83 | 0.84 | 10856 |
| 15 | 0.07 | 0.07 | 0.02 | 0.26 | 0.62 | 13959 | 12.90 | 0.50 | 12.65 | 13.94 | 0.26 | 26 | 0.43 | 0.18 | 0.31 | 0.80 | 0.54 | 10305 |
| 16 | 0.21 | 0.18 | 0.06 | 0.69 | 0.84 | 13112 | 12.89 | 0.40 | 12.79 | 14.01 | 0.33 | 19 | 0.61 | 0.23 | 0.45 | 1.09 | 0.77 | 4253 |
| 17 | 0.17 | 0.15 | 0.05 | 0.58 | 0.63 | 13773 | 18.17 | 0.92 | 18.29 | 21.04 | 0.10 | 21 | 0.91 | 0.21 | 0.76 | 1.35 | 0.38 | 730 |
| 18 | 0.08 | 0.07 | 0.02 | 0.28 | 0.71 | 14088 | 12.87 | 0.38 | 12.71 | 13.88 | 0.30 | 35 | 0.03 | 0.03 | 0.01 | 0.12 | 0.83 | 8563 |
| **19** | **0.40** | **0.30** | **0.16** | **1.13** | **0.06** | **7386** | **12.89** | **0.38** | **12.75** | **13.94** | **0.35** | **25** | **0.41** | **0.24** | **0.22** | **0.95** | **0.95** | **7431** |
| **20** | **0.41** | **0.31** | **0.17** | **1.15** | **0.99** | **7151** | **12.89** | **0.40** | **12.70** | **14.12** | **0.25** | **73** | **0.39** | **0.24** | **0.21** | **0.93** | **0.18** | **7664** |
| 21 | 0.26 | 0.22 | 0.09 | 0.82 | 0.31 | 11423 | 12.88 | 0.40 | 12.68 | 14.01 | 0.27 | 34 | 0.75 | 0.24 | 0.58 | 1.24 | 0.08 | 1094 |
| pop | 0.16 | 0.12 | 0.09 | 0.40 | 0.12 | 12604 | 12.88 | 0.37 | 12.71 | 13.94 | 0.39 | 23 | 0.40 | 0.08 | 0.35 | 0.56 | 0.24 | 3631 |

TABLE VIII. Parameter estimates for Model III. These results are obtained from the second-stage algorithm at the hierarchical level. The individuals for which this model was the best one, according to the comparison of Table II in the main paper, are highlighted in bold.

**Model IV**

| Parameter | $\alpha$ | | | | | | $\beta$ | | | | | | $\lambda$ | | | | | |
|---|---|---|---|---|---|---|---|---|---|---|---|---|---|---|---|---|---|---|
| ID | Mean | SD | 2.5% | 97.5% | $p_v$ | n_eff | Mean | SD | 2.5% | 97.5% | $p_v$ | n_eff | Mean | SD | 2.5% | 97.5% | $p_v$ | n_eff |
| 1 | 2.35 | 0.22 | 2.18 | 2.81 | 0.67 | 970 | 1.50 | 0.16 | 1.38 | 1.85 | 0.33 | 6377 | 2.99 | 0.28 | 2.79 | 3.57 | 0.07 | 1357 |
| 2 | 1.49 | 0.16 | 1.37 | 1.83 | 0.25 | 2443 | 1.01 | 0.06 | 0.97 | 1.13 | 0.66 | 12928 | 3.02 | 0.30 | 2.81 | 3.64 | 0.06 | 1508 |
| 3 | 1.65 | 0.11 | 1.57 | 1.87 | 0.96 | 6460 | 1.48 | 0.08 | 1.42 | 1.65 | 0.45 | 9131 | 3.11 | 0.26 | 2.94 | 3.68 | 0.08 | 1889 |
| 4 | 1.98 | 0.28 | 1.79 | 2.56 | 0.52 | 3802 | 1.19 | 0.21 | 1.04 | 1.64 | 0.49 | 11299 | 2.75 | 0.31 | 2.54 | 3.37 | 0.46 | 1321 |
| 5 | 2.25 | 0.27 | 2.08 | 2.79 | 0.37 | 79 | 1.09 | 0.17 | 0.96 | 1.48 | 74 | 12315 | 2.74 | 0.31 | 2.53 | 3.37 | 0.29 | 1979 |
| 6 | 2.43 | 0.23 | 2.26 | 2.91 | 0.53 | 372 | 1.15 | 0.09 | 1.09 | 1.35 | 0.78 | 13790 | 2.88 | 0.30 | 2.67 | 3.49 | 0.06 | 2227 |
| 7 | 1.70 | 0.21 | 1.55 | 2.13 | 0.99 | 4109 | 0.80 | 0.05 | 0.76 | 0.91 | 0.78 | 10722 | 3.02 | 0.27 | 2.83 | 3.60 | 0.07 | 2125 |
| 8 | 1.97 | 0.24 | 1.80 | 2.45 | 0.79 | 5352 | 1.47 | 0.21 | 1.32 | 1.92 | 0.86 | 5168 | 2.75 | 0.31 | 2.54 | 3.39 | 0.08 | 2041 |
| 9 | 2.14 | 0.22 | 1.99 | 2.60 | 0.98 | 4182 | 1.42 | 0.16 | 1.31 | 1.76 | 0.09 | 8359 | 2.79 | 0.31 | 2.58 | 3.43 | 0.02 | 1765 |
| 10 | 1.88 | 0.20 | 1.74 | 2.30 | 0.47 | 8236 | 1.35 | 0.14 | 1.25 | 1.66 | 0.47 | 10695 | 2.81 | 0.30 | 2.61 | 3.41 | 0.14 | 1941 |
| 11 | 1.93 | 0.13 | 1.84 | 2.22 | 0.22 | 11336 | 1.32 | 0.06 | 1.27 | 1.45 | 0.28 | 12685 | 2.97 | 0.28 | 2.77 | 3.54 | 0.52 | 2423 |
| 12 | 2.15 | 0.20 | 2.01 | 2.56 | 0.11 | 4512 | 1.37 | 0.12 | 1.28 | 1.63 | 0.77 | 10361 | 2.86 | 0.30 | 2.65 | 3.43 | 0.06 | 2049 |
| 13 | 1.95 | 0.18 | 1.82 | 2.33 | 0.27 | 7747 | 1.07 | 0.07 | 1.02 | 1.22 | 0.84 | 12726 | 3.03 | 0.30 | 2.82 | 3.63 | 0.51 | 1492 |
| 14 | 1.75 | 0.23 | 1.59 | 2.23 | 0.76 | 3838 | 1.00 | 0.08 | 0.93 | 1.21 | 0.10 | 11835 | 2.90 | 0.29 | 2.69 | 3.49 | 0.07 | 1644 |
| 15 | 1.63 | 0.13 | 1.54 | 1.89 | 0.49 | 5737 | 1.30 | 0.09 | 1.24 | 1.49 | 0.47 | 12558 | 3.30 | 0.28 | 3.15 | 3.89 | 0.00 | 223 |
| 16 | 1.89 | 0.29 | 1.69 | 2.49 | 0.06 | 4172 | 0.88 | 0.15 | 0.77 | 1.22 | 0.17 | 7911 | 2.70 | 0.31 | 2.49 | 3.36 | 0.19 | 1164 |
| 17 | 1.88 | 0.27 | 1.70 | 2.42 | 0.44 | 4538 | 1.03 | 0.15 | 0.92 | 1.34 | 0.28 | 11033 | 2.73 | 0.32 | 2.51 | 3.39 | 0.68 | 1620 |
| 18 | 1.59 | 0.18 | 1.46 | 1.95 | 0.62 | 3082 | 1.01 | 0.03 | 0.97 | 1.15 | 0.00 | 12335 | 2.92 | 0.29 | 2.73 | 3.50 | 0.01 | 1852 |
| 19 | 2.00 | 0.36 | 1.75 | 2.69 | 0.49 | 162 | 1.06 | 0.35 | 0.82 | 1.74 | 0.88 | 5757 | 2.60 | 0.37 | 2.35 | 3.42 | 0.68 | 164 |
| 20 | 1.95 | 0.33 | 1.73 | 2.60 | 0.98 | 543 | 1.08 | 0.33 | 0.86 | 1.75 | 0.06 | 6780 | 2.64 | 0.35 | 2.41 | 3.30 | 0.19 | 239 |
| 21 | 1.86 | 0.31 | 1.66 | 2.46 | 0.50 | 2917 | 0.82 | 0.17 | 0.70 | 1.20 | 0.58 | 6160 | 2.69 | 0.32 | 2.48 | 3.30 | 0.14 | 1048 |
| pop | 1.93 | 0.10 | 1.86 | 2.12 | 0.47 | 528 | 1.16 | 0.09 | 1.10 | 1.34 | 0.18 | 8465 | 2.87 | 0.15 | 2.76 | 3.17 | 0.33 | 295 |

| Parameter | $\kappa$ | | | | | | $\rho$ | | | | | | $\epsilon$ | | | | | |
|---|---|---|---|---|---|---|---|---|---|---|---|---|---|---|---|---|---|---|
| ID | Mean | SD | 2.5% | 97.5% | $p_v$ | n_eff | Mean | SD | 2.5% | 97.5% | $p_v$ | n_eff | Mean | SD | 2.5% | 97.5% | $p_v$ | n_eff |
| 1 | 0.14 | 0.12 | 0.04 | 0.47 | 0.97 | 14853 | 21.02 | 0.46 | 20.69 | 21.93 | 0.63 | 101 | 0.67 | 0.22 | 0.51 | 1.12 | 0.40 | 2782 |
| 2 | 0.07 | 0.07 | 0.02 | 0.28 | 0.96 | 14136 | 21.00 | 0.52 | 20.66 | 22.03 | 0.66 | 119 | 0.33 | 0.19 | 0.18 | 0.72 | 0.69 | 9678 |
| 3 | 0.06 | 0.06 | 0.01 | 0.25 | 0.94 | 13726 | 21.05 | 0.51 | 20.71 | 22.08 | 0.78 | 61 | 0.39 | 0.22 | 0.21 | 0.83 | 0.26 | 9599 |
| 4 | 0.22 | 0.19 | 0.07 | 0.71 | 0.68 | 12822 | 21.04 | 0.49 | 20.72 | 21.96 | 0.16 | 89 | 0.61 | 0.24 | 0.44 | 1.09 | 0.84 | 3159 |
| 5 | 0.18 | 0.15 | 0.06 | 0.60 | 0.59 | 13474 | 21.00 | 0.49 | 20.68 | 21.85 | 0.70 | 73 | 0.30 | 0.17 | 0.17 | 0.69 | 0.33 | 9385 |
| 6 | 0.12 | 0.11 | 0.03 | 0.42 | 0.82 | 14093 | 20.96 | 0.51 | 20.69 | 20.72 | 0.38 | 113 | 0.04 | 0.04 | 0.01 | 0.17 | 0.89 | 7815 |
| 7 | 0.08 | 0.08 | 0.02 | 0.30 | 0.78 | 14512 | 20.99 | 0.45 | 20.71 | 22.12 | 0.63 | 76 | 0.12 | 0.23 | 0.00 | 0.81 | 0.14 | 6012 |
| 8 | 0.19 | 0.17 | 0.06 | 0.63 | 0.57 | 11360 | 20.99 | 0.47 | 20.68 | 22.01 | 0.21 | 95 | 0.63 | 0.25 | 0.46 | 1.14 | 0.74 | 2286 |
| 9 | 0.11 | 0.10 | 0.03 | 0.39 | 0.14 | 14562 | 21.07 | 0.47 | 20.81 | 22.03 | 0.69 | 90 | 0.28 | 0.21 | 0.09 | 0.72 | 0.34 | 9042 |
| 10 | 0.10 | 0.09 | 0.03 | 0.37 | 0.44 | 14268 | 21.10 | 1.00 | 20.28 | 23.45 | 0.19 | 125 | 0.70 | 0.27 | 0.55 | 1.23 | 0.29 | 1321 |
| 11 | 0.06 | 0.06 | 0.01 | 0.26 | 0.10 | 13954 | 21.11 | 0.80 | 20.87 | 21.83 | 0.27 | 64 | 0.06 | 0.03 | 0.04 | 0.14 | 0.96 | 10617 |
| 12 | 0.08 | 0.08 | 0.02 | 0.30 | 0.54 | 13628 | 21.04 | 0.45 | 20.70 | 22.06 | 0.94 | 88 | 0.29 | 0.23 | 0.06 | 0.76 | 0.78 | 8416 |
| 13 | 0.08 | 0.08 | 0.02 | 0.30 | 0.16 | 13312 | 21.02 | 0.46 | 20.73 | 21.91 | 0.97 | 87 | 0.04 | 0.06 | 0.00 | 0.24 | 0.20 | 5730 |
| 14 | 0.10 | 0.09 | 0.03 | 0.34 | 0.57 | 14017 | 21.05 | 0.47 | 20.73 | 22.19 | 0.89 | 83 | 0.38 | 0.21 | 0.22 | 0.81 | 0.63 | 10255 |
| 15 | 0.06 | 0.06 | 0.01 | 0.24 | 0.49 | 13450 | 21.01 | 0.56 | 20.73 | 22.06 | 0.37 | 83 | 0.31 | 0.30 | 0.01 | 0.92 | 0.66 | 7598 |
| 16 | 0.20 | 0.18 | 0.06 | 0.68 | 0.51 | 13340 | 21.07 | 0.47 | 20.76 | 22.03 | 0.74 | 96 | 0.64 | 0.22 | 0.49 | 1.10 | 0.74 | 3125 |
| 17 | 0.17 | 0.15 | 0.05 | 0.58 | 0.64 | 13454 | 21.30 | 0.44 | 21.16 | 22.20 | 0.87 | 65 | 0.86 | 0.22 | 0.70 | 1.33 | 0.23 | 570 |
| 18 | 0.07 | 0.07 | 0.02 | 0.27 | 0.36 | 14172 | 20.99 | 0.49 | 20.72 | 22.13 | 0.53 | 51 | 0.03 | 0.03 | 0.00 | 0.13 | 0.39 | 8109 |
| 19 | 0.40 | 0.30 | 0.16 | 1.15 | 0.43 | 7734 | 20.98 | 0.58 | 20.51 | 21.79 | 0.83 | 42 | 0.41 | 0.24 | 0.22 | 0.94 | 0.87 | 7101 |
| 20 | 0.40 | 0.31 | 0.17 | 1.16 | 0.52 | 7398 | 21.09 | 0.59 | 20.95 | 22.04 | 0.23 | 74 | 0.39 | 0.24 | 0.21 | 0.94 | 0.70 | 7085 |
| 21 | 0.26 | 0.22 | 0.08 | 0.81 | 0.52 | 11560 | 21.01 | 0.48 | 20.69 | 22.03 | 0.97 | 87 | 0.74 | 0.24 | 0.57 | 1.24 | 0.79 | 1128 |
| pop | 0.15 | 0.13 | 0.08 | 0.39 | 0.88 | 13073 | 21.04 | 0.35 | 20.82 | 21.93 | 0.98 | 41 | 0.39 | 0.08 | 0.33 | 0.57 | 0.71 | 3416 |

| Parameter | $\nu$ | | | | | | $\theta$ | | | | | |
|---|---|---|---|---|---|---|---|---|---|---|---|---|
| ID | Mean | SD | 2.5% | 97.5% | $p_v$ | n_eff | Mean | SD | 2.5% | 97.5% | $p_v$ | n_eff |
| 1 | 10.78 | 1.30 | 10.17 | 12.61 | 0.06 | 62 | 0.51 | 0.25 | 0.31 | 1.06 | 0.83 | 7063 |
| 2 | 10.77 | 1.31 | 10.23 | 12.47 | 0.12 | 36 | 0.23 | 0.16 | 0.10 | 0.62 | 0.26 | 12431 |
| 3 | 10.79 | 1.30 | 10.24 | 12.59 | 0.06 | 43 | 0.16 | 0.10 | 0.09 | 0.41 | 0.56 | 12266 |
| 4 | 10.78 | 1.30 | 10.22 | 12.54 | 0.09 | 35 | 0.32 | 0.24 | 0.13 | 0.91 | 0.13 | 9510 |
| 5 | 10.76 | 1.28 | 10.10 | 12.44 | 0.08 | 36 | 0.55 | 0.28 | 0.34 | 1.16 | 0.61 | 5464 |
| 6 | 10.77 | 1.31 | 10.24 | 12.50 | 0.06 | 46 | 0.28 | 0.20 | 0.12 | 0.75 | 0.16 | 11643 |
| 7 | 10.82 | 1.31 | 10.22 | 12.61 | 0.10 | 35 | 0.38 | 0.22 | 0.21 | 0.91 | 0.35 | 10373 |
| 8 | 10.78 | 1.31 | 10.18 | 12.66 | 0.10 | 33 | 0.24 | 0.20 | 0.08 | 0.75 | 0.28 | 11858 |
| 9 | 10.77 | 1.32 | 10.19 | 12.65 | 0.13 | 37 | 0.12 | 0.12 | 0.03 | 0.44 | 0.28 | 11260 |
| 10 | 10.77 | 1.30 | 10.17 | 12.64 | 0.05 | 39 | 0.22 | 0.18 | 0.08 | 0.68 | 0.51 | 12393 |
| 11 | 10.77 | 1.30 | 10.22 | 12.51 | 0.05 | 30 | 0.34 | 0.17 | 0.21 | 0.74 | 0.64 | 12120 |
| 12 | 10.77 | 1.31 | 10.18 | 12.57 | 0.11 | 37 | 0.20 | 0.16 | 0.07 | 0.60 | 0.91 | 12778 |
| 13 | 10.79 | 1.31 | 10.30 | 12.68 | 0.07 | 48 | 0.30 | 0.19 | 0.15 | 0.73 | 0.82 | 12115 |
| 14 | 10.78 | 1.31 | 10.20 | 12.58 | 0.10 | 27 | 0.30 | 0.18 | 0.15 | 0.74 | 0.81 | 11795 |
| 15 | 10.78 | 1.30 | 10.11 | 12.56 | 0.10 | 46 | 0.06 | 0.06 | 0.02 | 0.22 | 0.51 | 1233 |
| 16 | 10.77 | 1.30 | 10.24 | 12.58 | 0.08 | 36 | 0.38 | 0.27 | 0.16 | 1.00 | 0.35 | 8011 |
| 17 | 10.80 | 1.30 | 10.16 | 12.71 | 0.08 | 41 | 0.27 | 0.21 | 0.09 | 0.80 | 0.39 | 11298 |
| 18 | 10.78 | 1.32 | 10.15 | 12.58 | 0.09 | 37 | 0.32 | 0.18 | 0.18 | 0.75 | 0.60 | 12968 |
| 19 | 10.78 | 1.31 | 10.20 | 12.54 | 0.05 | 33 | 0.41 | 0.28 | 0.18 | 1.08 | 0.21 | 7167 |
| 20 | 10.76 | 1.32 | 10.14 | 12.61 | 0.10 | 50 | 0.40 | 0.27 | 0.18 | 1.04 | 0.46 | 7375 |
| 21 | 10.76 | 1.32 | 10.15 | 12.69 | 0.08 | 38 | 0.41 | 0.27 | 0.19 | 1.03 | 0.22 | 7096 |
| pop | 10.78 | 1.27 | 10.31 | 12.46 | 0.31 | 9 | 0.30 | 0.10 | 0.24 | 0.50 | 0.24 | 8629 |

TABLE IX. Parameter estimates for Model IV. These results are obtained from the second-stage algorithm at the hierarchical level.

##### Marginal posteriors of the spatial parameters

The figures 2-5 display the mean values and the 95% CIs of the spatial parameters common to Models I-IV obtained from the marginal posterior distributions, for each animal (denoted by individual ID number) and for the population (denoted by 'pop'). The black intervals (to the left for each individual) correspond to the results from the first-stage algorithm, *i.e.*, the individual-level fit of each parameter $p_j$ for each animal. The light gray intervals to the right result from the second-stage algorithm, *i.e.*, the fit at the hierarchical level of each parameter $p_j$. The dark gray interval above 'pop' corresponds to the second-stage algorithm for the fit of the population parameter $p$. In blue are indicated the individuals for which the corresponding model is the best according to the results shown in Table V.

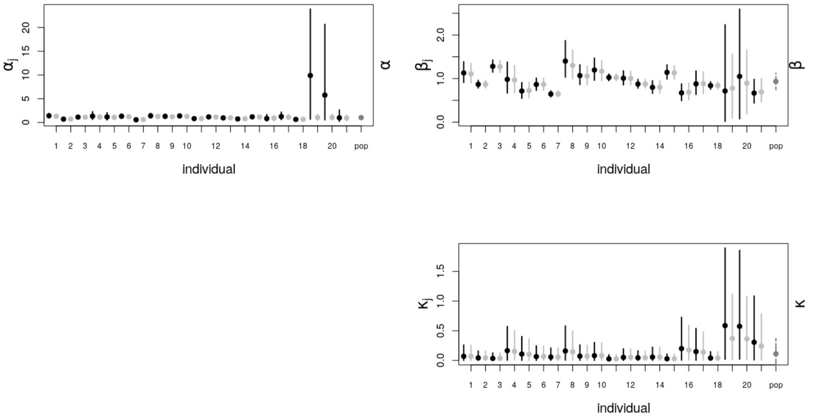


FIG. 2. Means (points) and 95% CIs (vertical lines) of the parameters of Model I, obtained from the marginal posterior distributions of the individuals (denoted by individual ID number). The black intervals (to the left of each individual) correspond to the results of the first-stage algorithm, and the light gray intervals (to the right) to the results of the second-stage algorithm. The dark gray interval above 'pop' corresponds to the results of second-stage algorithm at the population-level.

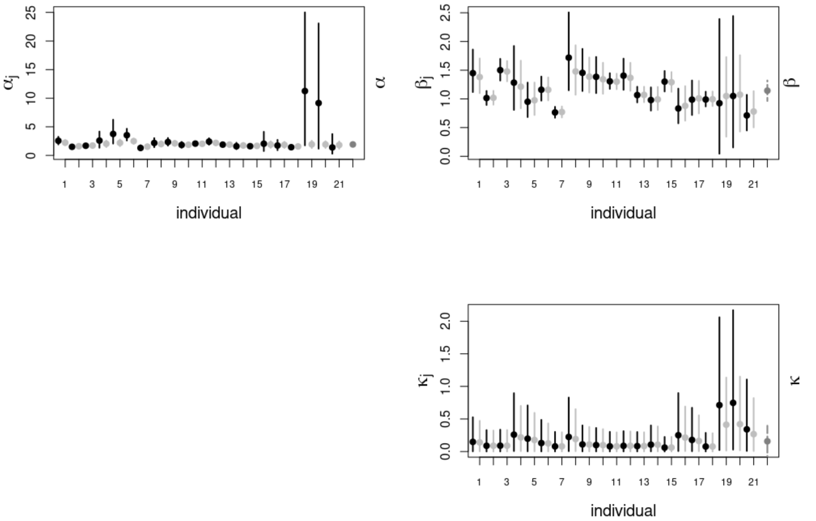


FIG. 3. As same as Fig.2, for Model II.

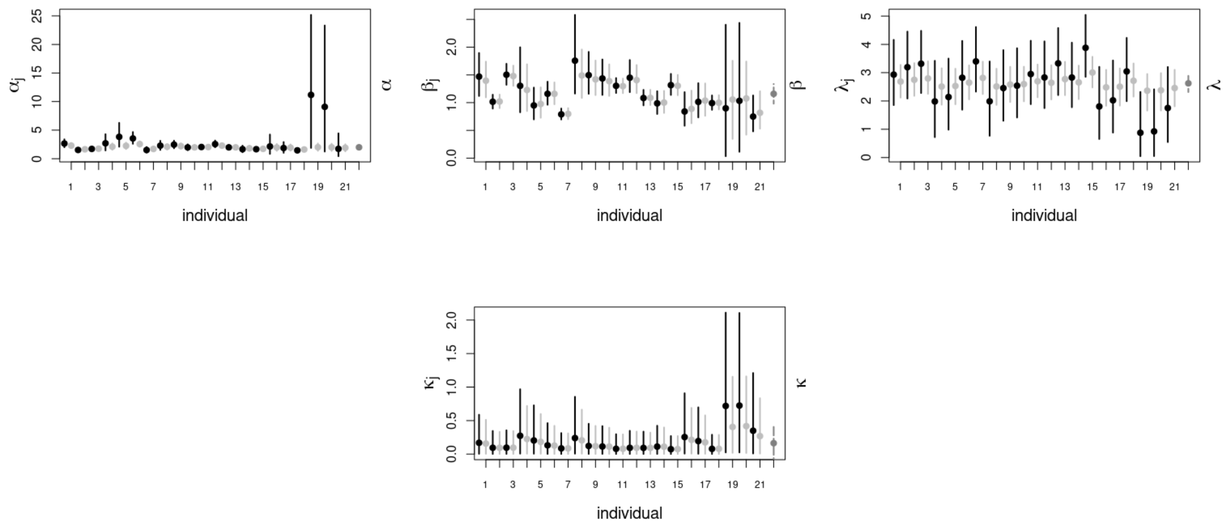

FIG. 4. As same as Fig.2, for Model III.

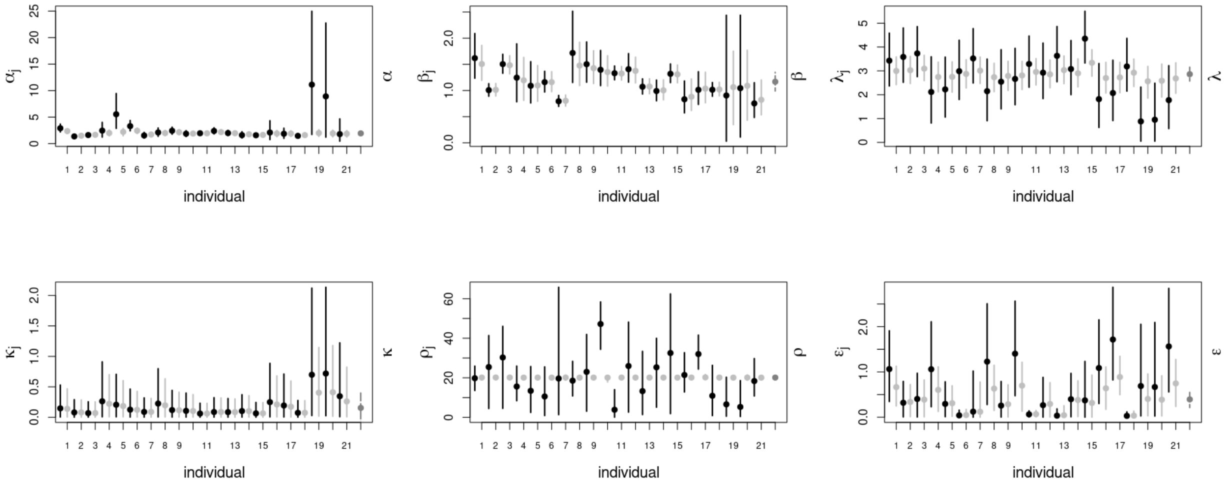

FIG. 5. As same as Fig.2, for Model IV.

The next three figures display the variations of the probability weight d with the distance to the patch, the variations the probability weight c with the patch area, and the memory use parameter ($q$) as function of the number of unique visit sites ($\mathbf{u}$) (in the case of Model III and IV). These results correspond to the population level. To obtain these graphs, many curves were generated (in light grey) by sampling the marginal distribution of each parameter. The CIs at 95% are indicated in dark grey.

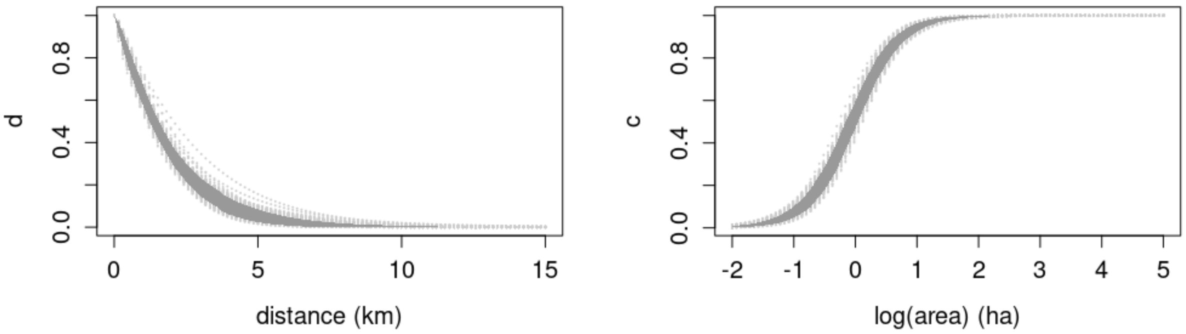


FIG. 6. Model II: Population-level probability weights for choosing a patch as a function of the distance (**Left**) and of the area (**Right**). The curves are obtained from sampling the posterior distributions, with the 95% CIs shown in dark grey.

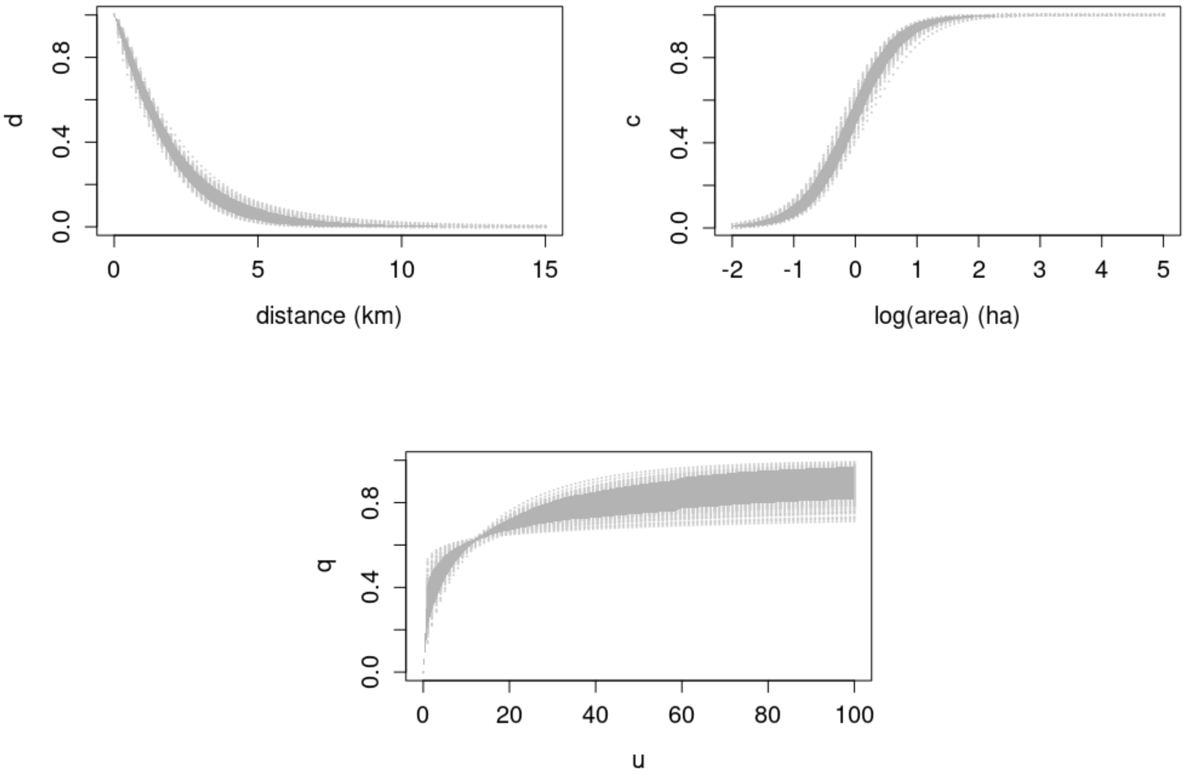


FIG. 7. Model III: Population-level probability weights for choosing a patch as a function of the distance (**Top Left**) and of the area (**Top Right**). Increase of the memory parameter as a function of the number of unique visited sites (**Bottom**). The curves are obtained from sampling the posterior distributions, with the 95% CIs shown in dark grey.

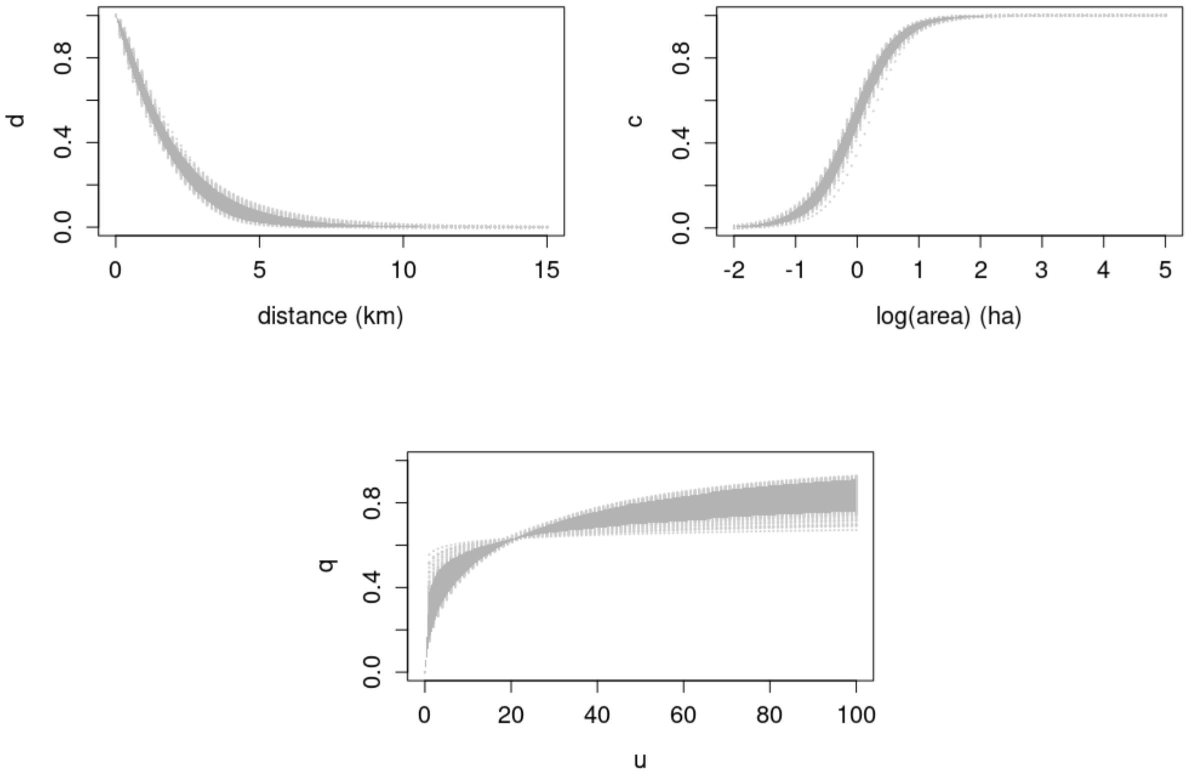


FIG. 8. Same as Figure 7, for Model IV.

### PPC

Each row of the Figure below displays the PPCs of all models, for a same individual. Thus, each row allows us to compare the different models for animal $i$, with $i = 1 : 21$. The red symbols show the observed number of unique visited sites, whereas the light grey symbols represent the PPCs generated by 1000 independent samples, and the dark grey symbols the 95% CIs.

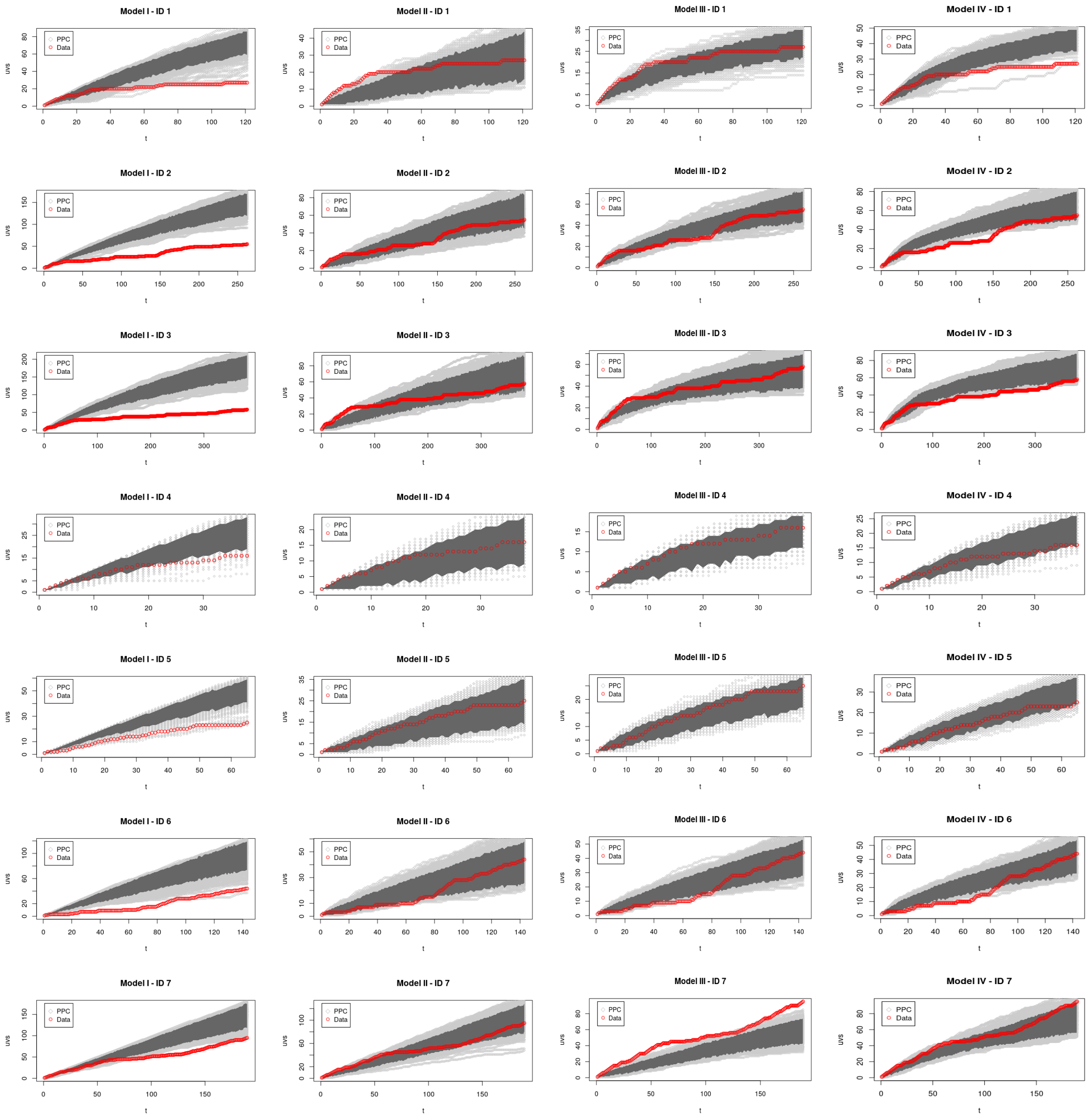

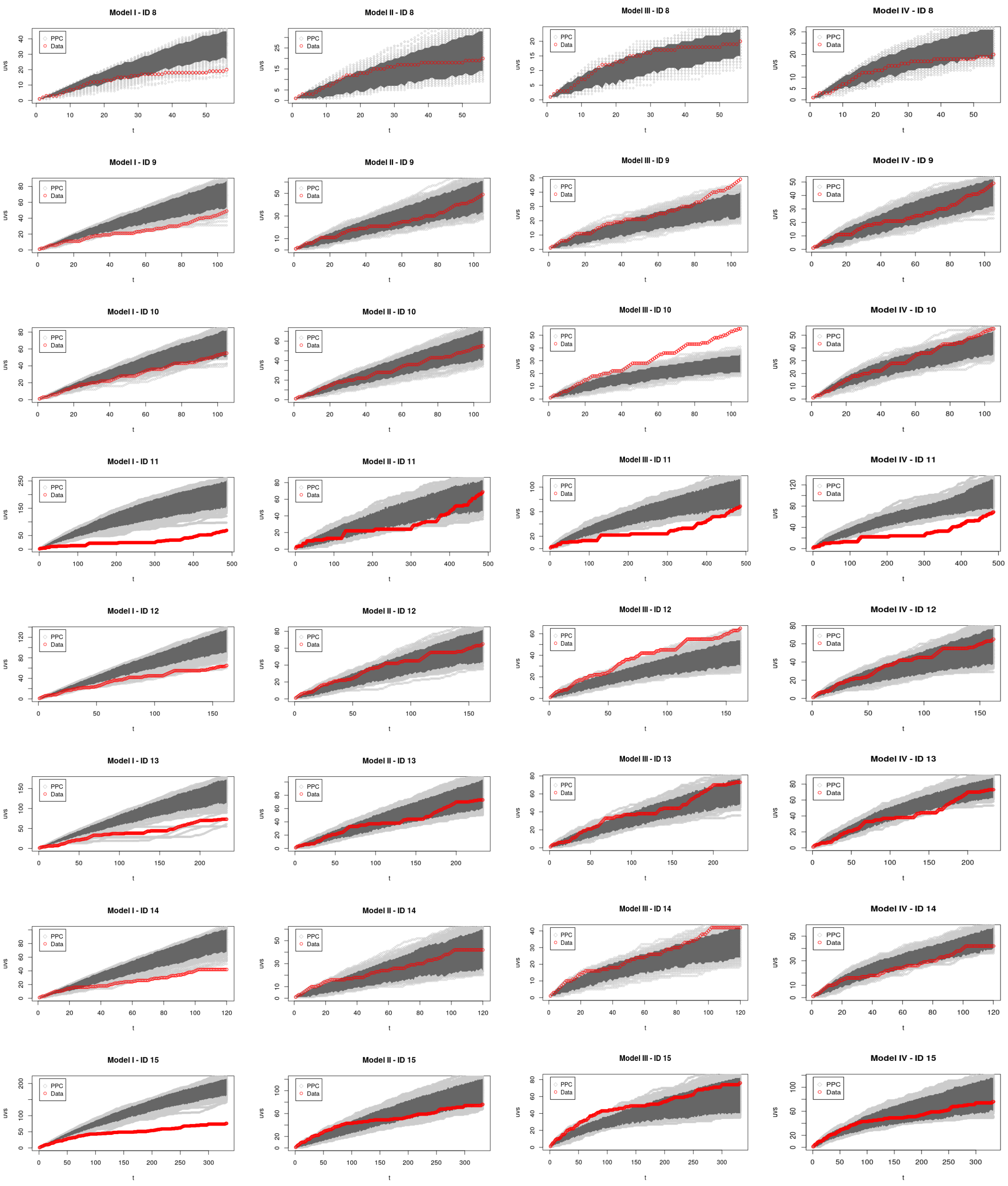
Model I - ID 8
Model II - ID 8
Model III - ID 8
Model IV - ID 8
Model I - ID 9
Model II - ID 9
Model III - ID 9
Model IV - ID 9
Model I - ID 10
Model II - ID 10
Model III - ID 10
Model IV - ID 10
Model I - ID 11
Model II - ID 11
Model III - ID 11
Model IV - ID 11
Model I - ID 12
Model II - ID 12
Model III - ID 12
Model IV - ID 12
Model I - ID 13
Model II - ID 13
Model III - ID 13
Model IV - ID 13
Model I - ID 14
Model II - ID 14
Model III - ID 14
Model IV - ID 14
Model I - ID 15
Model II - ID 15
Model III - ID 15
Model IV - ID 15
PPC
Data
uvs
t

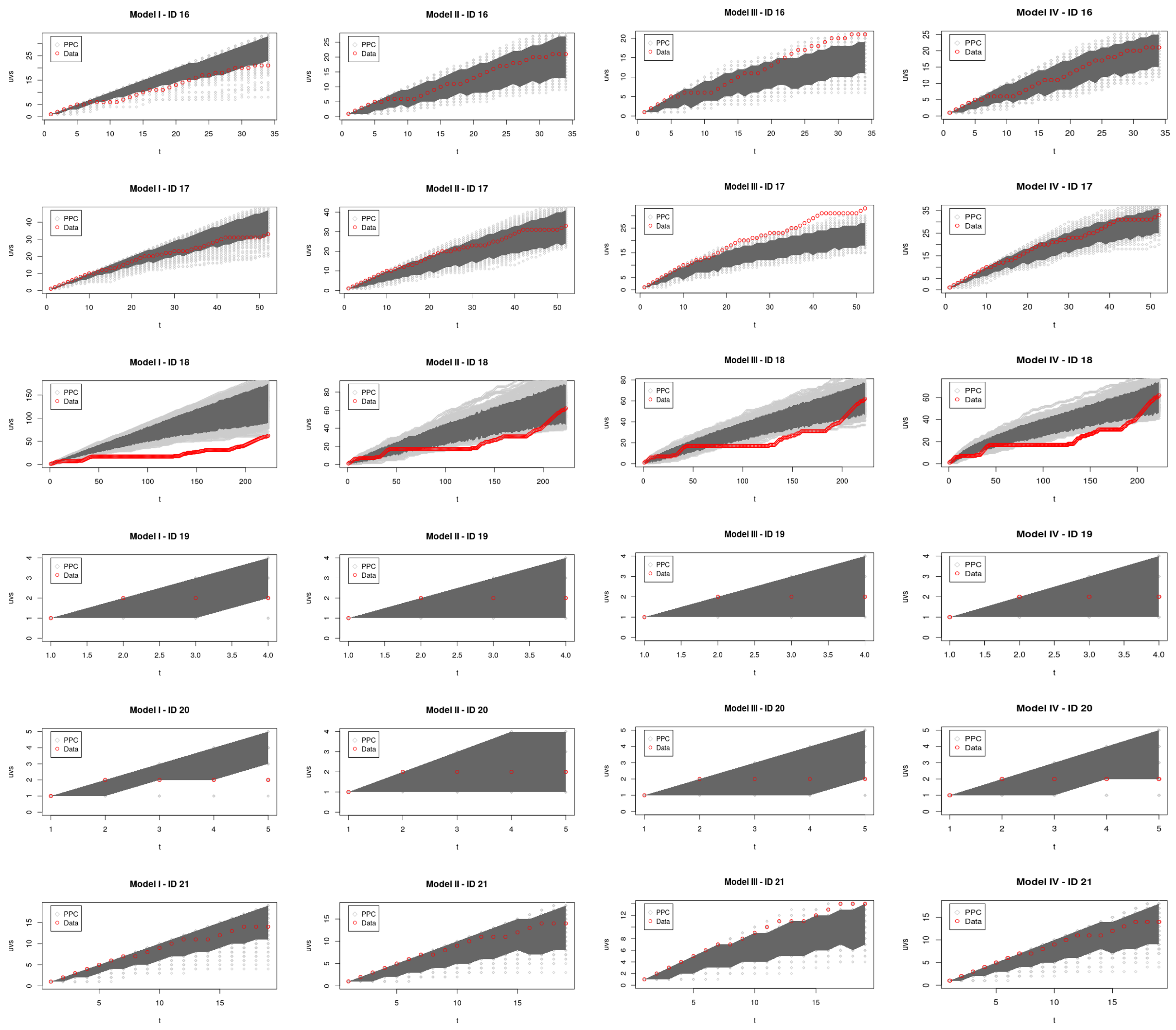

FIG. 9. Posterior predictive checks of Models I-IV (columns) for all the observed animals (rows), at the hierarchical level. Each panel shows the number of unique visited sites (UVS) as a function of time $t$. The PPC curves are shown in light grey, the CIs at 95% in dark grey and the data from the real trajectories in red. The $y$-scale of the graphs in a same row (individual) may vary in order to better show which parts of the real trajectory are inside of the CI.
